# Efficient Inference of Macrophylogenies: Insights from the Avian Tree of Life

**DOI:** 10.1101/2025.02.17.638170

**Authors:** Min Zhao, Gregory Thom, Brant C. Faircloth, Michael J. Andersen, F. Keith Barker, Brett W. Benz, Michael J. Braun, Gustavo A. Bravo, Robb T. Brumfield, R. Terry Chesser, Elizabeth P. Derryberry, Travis C. Glenn, Michael G. Harvey, Peter A. Hosner, Tyler S. Imfeld, Leo Joseph, Joseph D. Manthey, John E. McCormack, Jenna M. McCullough, Robert G. Moyle, Carl H. Oliveros, Noor D. White Carreiro, Kevin Winker, Daniel J. Field, Daniel T. Ksepka, Edward L. Braun, Rebecca T. Kimball, Brian Tilston Smith

## Abstract

The exponential growth of molecular sequence data over the past decade has enabled the construction of numerous clade-specific phylogenies encompassing hundreds or thousands of taxa. These independent studies often include overlapping data, presenting a unique opportunity to build macrophylogenies (phylogenies sampling > 1,000 taxa) for entire classes across the Tree of Life. However, the inference of large trees remains constrained by logistical, computational, and methodological challenges. The Avian Tree of Life provides an ideal model for evaluating strategies to robustly infer macrophylogenies from intersecting datasets derived from smaller studies. In this study, we leveraged a comprehensive resource of sequence capture datasets to evaluate the phylogenetic accuracy and computational costs of four methodological approaches: (1) supermatrix approaches using concatenation, including the “fast” maximum likelihood (ML) methods, (2) filtering datasets to reduce heterogeneity, (3) supertree estimation based on published phylogenomic trees, and (4) a “divide-and-conquer” strategy, wherein smaller ML trees were estimated and subsequently combined using a supertree approach. Additionally, we examined the impact of these methods on divergence time estimation using a dataset that includes newly vetted fossil calibrations for the Avian Tree of Life. Our findings highlight that recently developed fast tree search approaches offer a reasonable compromise between computational efficiency and phylogenetic accuracy, facilitating inference of macrophylogenies.

## Introduction

Completing the Tree of Life remains a significant bottleneck to addressing a wide range of questions in comparative biology (Cracraft and Donoghue 2004). Advances in sequencing technologies (e.g., see McCormack et al. (2013) for review), computational methods (e.g., Kozlov et al. 2019), and user-friendly bioinformatic pipelines (e.g., Faircloth 2016) have made the production and analysis of phylogenomic datasets involving hundreds of taxa increasingly routine. However, scaling these techniques to datasets with thousands of loci and thousands of taxa presents substantial logistical, computational, and methodological challenges (Delsuc et al. 2005; Philippe et al. 2011; Kapli et al. 2020). The construction of such “macrophylogenies” (Title and Rabosky 2017) often relies on combining independently produced datasets, which frequently have limited overlap and substantial missing data (Sanderson et al. 2010).

Past attempts to infer macrophylogenies from independently-produced datasets typically used two general approaches: supermatrix and supertree methods. Supermatrix methods infer phylogenies directly from orthologous loci, often compiled from multiple studies. However, these methods are negatively affected by large amounts of missing data (Driskell et al. 2004; Philippe et al. 2004; Goloboff et al. 2009; Hosner et al. 2016) and varying standards of data quality (Philippe et al. 2011). Analyses of supermatrices are also vulnerable to common issues in phylogenetic analyses, such as alignment errors (Ogden and Rosenberg 2006) and the inclusion of non-orthologous sequences (Koonin 2005), which are often exacerbated in supermatrices due to the heterogeneous nature of the data. Additionally, supermatrix methods face escalating computational demands that increase nonlinearly (Bader et al. 2006) as both the width (number of sites) and height (number of taxa) of the matrix expand (Delsuc et al. 2005). Some challenges, such as data quality and alignment issues, can be mitigated to an extent by analyzing multiple datasets filtered to remove “noise” in different ways and comparing the results (Kuhl et al. 2021). However, this approach is limited by the significant computational costs of performing multiple analyses on large datasets. Supertree methods, by contrast, generate phylogenies by combining existing tree topologies (Sanderson et al. 1998; Bininda-Emonds 2004; Cotton and Wilkinson 2009). These methods are more computationally efficient and can effectively incorporate trees built with heterogeneous data (Liu et al. 2001; Hinchliff et al. 2015; Redelings and Holder 2017). However, most supertree methods cannot directly estimate meaningful branch lengths. Despite the strengths and limitations of these methods, rigorous comparisons of the ability of supermatrix and supertree methods to estimate macrophylogenies using phylogenomic data remain rare. This gap largely reflects the limited availability of large-scale genomic datasets for most taxonomic groups.

Class Aves (birds) is one taxonomic group with sufficient data to perform these types of comparative analyses. As the most species-rich terrestrial vertebrate group, with 11,140 species recognized (Gill et al. 2023), birds have received extensive attention from phylogenetic systematists (e.g., Hackett et al. 2008; Jetz et al. 2012; McCormack, Harvey, et al. 2013; Jarvis et al. 2014; Burleigh et al. 2015; Prum et al. 2015; Moyle et al. 2016; Reddy et al. 2017; Oliveros et al. 2019; Harvey et al. 2020; Stiller et al. 2024). Many relationships among birds are now strongly corroborated across studies, providing a reliable framework for evaluating the accuracy of alternative approaches to estimate macrophylogenies.

Another advantage of birds as a model system is the partial standardization of phylogenomic data collection through the widespread use of targeted enrichment of nuclear loci, such as ultraconserved elements (UCEs *sensu* Faircloth et al. 2012). Over a quarter of all avian species now have UCE data available (see below). These data have been used to resolve phylogenetic relationships among birds at both deep (e.g., McCormack et al. 2013; Jarvis et al. 2014; Oliveros et al. 2019; Harvey et al. 2020) and shallow (e.g., Smith et al. 2014; Winker et al. 2018) timescales. Most UCE studies of birds target a large, uniform set of loci (the publicly available uce-5k-probe-set (https://github.com/faircloth-lab/uce-probe-sets; e.g., Sun et al. 2014). Some studies instead use a smaller, nested subset of these loci (uce-2.5k-probe-set) that is sometimes combined with exons commonly used in avian phylogenetics (e.g., Smith et al. 2014; Harvey et al. 2020). Although these datasets exhibit some heterogeneity – stemming from the use of different bait sets and variability in the quality of input DNA templates – extensive overlap facilitates integration into a single comprehensive dataset.

In this study, we use phylogenomic data from birds to empirically evaluate the accuracy and computational cost of alternative tree estimation approaches. By assembling orthologous UCE loci from the primary literature, we aim to better understand the factors influencing the estimation of macrophylogenies. Specifically, we address the following questions: 1) Does filtering datasets to reduce size and heterogeneity result in topologies that recover expected clades corroborated in prior studies? 2) Do computationally efficient methods, such as “fast” maximum likelihood (ML) estimation, supertrees, or a divide-and-conquer strategy that combines many small trees using a supertree method, recover similar numbers of expected relationships as traditional ML methods? 3) Does the use of different methods, which may bias branch length estimation and produce distinct topologies, affect divergence time estimation? By combining phylogenomic data from independent studies, we constructed a large-scale phylogeny for a major vertebrate group (Class: Aves), encompassing 2,756 ingroup taxa, 2 outgroup taxa and 5,121 loci. Our findings demonstrate that it is possible to infer an accurate macrophylogeny with moderate computational cost. Moreover, the strategies identified as most effective in this study are likely applicable to other taxonomic groups with sufficient phylogenomic data.

## Materials and Methods

### Assembling the phylogenomic data

We took multiple approaches to create a database of UCE loci from existing studies of birds. We downloaded much of the data as individual UCE alignments from 22 phylogenomic studies (Zhang et al. 2014; Bryson et al. 2016; Hosner et al. 2016; Manthey et al. 2016; McCormack et al. 2016; Burga et al. 2017; Campillo et al. 2018; Andermann et al. 2019; Andersen et al. 2019; Everson et al. 2019; Jenna M McCullough et al. 2019; Jenna M. McCullough et al. 2019; Oliveros et al. 2019; Sackton et al. 2019; White and Braun 2019; Harvey et al. 2020; Imfeld et al. 2020; Oliveros et al. 2020; Salter et al. 2020; Smith et al. 2023; Braun et al. 2024; for details, see Supplementary Table S1 & Supplementary Information). When downloading data from previous phylogenomic studies, we noticed that several studies had overlapping or nested taxon sampling. For example, Moyle et al. (2016) collected UCE data for 104 songbird species (Moyle et al., 2016), and these data had all been included in a later study with broader taxon sampling (Oliveros et al., 2019). Therefore, we used the “Oliveros2019” dataset for downstream analyses.

All studies targeted UCEs as the main genetic markers (some also targeted a small number of legacy markers), and we preferentially downloaded alignments with as little filtering as possible (e.g., no missing data cut-offs). For studies where individual alignments were unavailable, we downloaded concatenated matrices and partition files, which we converted into alignments using the “split” function of AMAS (Borowiec 2016). Finally, we extracted UCE loci from genome assemblies available at NCBI (that were not under embargo; data downloaded on October 14, 2020) for species that were not represented by UCE sequences. Genomes were retrieved based on the BioProject number available in publications using Reutils v0.2.2 (https://github.com/gschofl/reutils), and we extracted UCE loci and 500 bp flanking sequence following Tutorial III of PHYLUCE (Faircloth 2016) with the 5k probe set.

We converted the sequences of individual loci from different studies to FASTA format using PHYLUCE, standardized locus names across datasets, and combined homologous sequences from the different sources. We processed the sequences to retain only one individual per species, according to the IOC World Bird List v13.1 (Gill et al. 2023), although there were a few exceptions that resulted from taxonomic changes that occasionally led to a small number of duplicates or lumping some species together as subspecies (see Data Availability). When multiple individuals of the same species were present in our alignments or the same sample was used in different studies, we arbitrarily selected the representative sample based on the alphabetical order of the studies (Supplementary Table S1). After verifying taxa, we performed sequence alignment with MAFFT (Katoh and Standley 2013) using default settings and the --adjustdirection option to correct for sequence orientation. Then, we filtered raw alignments with trimAl (Capella-Gutiérrez et al. 2009) using the “gappyout” method to remove sites based on the gap distribution within each alignment. We refer to these alignments as the “full” dataset.

### Evaluating data heterogeneity

The starting quality of the sample, bioinformatic processing, and filtering schemes applied in different studies can bias the distribution of phylogenetic information within and between loci, directly impacting topology and branch length estimation (Molloy and Warnow 2018). We anticipated substantial heterogeneity in the original datasets used to generate our supermatrix, as the data were collected using different sequencing techniques (e.g., target enrichment vs. whole genome sequencing), bait sets (uce-5k-probe-set vs. uce-2.5k-probe-set), taxonomic samples (e.g., studies focused on all parrots (Smith et al. 2023) vs. all suboscine passerines (Harvey et al. 2020)), and sample sources (modern tissue samples vs. toe pads from historical museum specimens). We examined if data obtained from distinct studies could lead to a biased distribution of phylogenetic information by testing if samples could be classified according to their source study based on summary statistics. In total, we included 11 individual-based summary statistics for each sample (see Data Availability). We computed locus length and number of gaps using ape v5.6 (Paradis and Schliep 2019), GC content using seqinr v4.2 (Charif and Lobry 2007), and the number of singletons using vcftools v0.1.15 (Danecek et al. 2011) after converting alignments from FASTA to VCF (variant call format) format using SNP-sites (Page et al. 2016). We also used vcftools to compute the mean and standard deviation in the count of individual-based parsimony informative (IPI) sites present for an individual (out of all IPI sites identified across the entire data set). The number of IPI sites per individual was obtained by counting the number of alternative alleles that an individual carries that were present in two or more individuals (non-singleton), assuming *Alligator mississippiensis* (one of the outgroups) as a reference. To visualize the variation in summary statistics among studies, we built a heatmap of normalized summary statistics per individual using *gplots* v3.1.3.1 (https://github.com/talgalili/gplots) in R (R Core Team 2023).

### Filtering loci and subsetting datasets

Along with the full dataset, we wanted to investigate the effects of different locus filtering schemes on the topologies inferred, so we created 27 additional datasets that resulted from the application of three filtering schemes applied serially to the full dataset (Fig. 1a). To begin, we wanted to control for missing sequence data by taxon, so we prepared two datasets where we removed taxa from alignments when they were shorter than 50% or 75% of the longest sequence in the alignment for each locus (Fig. 1a, Step I). This step helps to control for the effects of partial sequences, i.e., “type II” missing data (*sensu* Hosner et al. 2016). Then we ran these two datasets, plus the full dataset, through a second stage of filtering to control for gappyness by retaining alignment positions with 90%, 70%, and 50% occupancy (Fig. 1a, Step II). This step helps to address potential issues with indel-induced alignment gaps (e.g., Dwivedi and Gadagkar 2009) and reduce heterogeneity that can occur at the ends of UCE alignments. Finally, for each of the nine datasets that resulted, we performed a third stage of filtering to control for taxon completeness, where we retained loci with 90% (n = 2484), 70% (n = 1932), and 50% (n = 1380) of the total number of taxa (Fig. 1a, Step III). The last step helps to control for the effects of incomplete taxon sampling, i.e., “type I” missing data (*sensu* Hosner et al. 2016). We concatenated each of these datasets using PHYLUCE (Faircloth 2016) prior to phylogenetic analysis.

**Figure 1.**
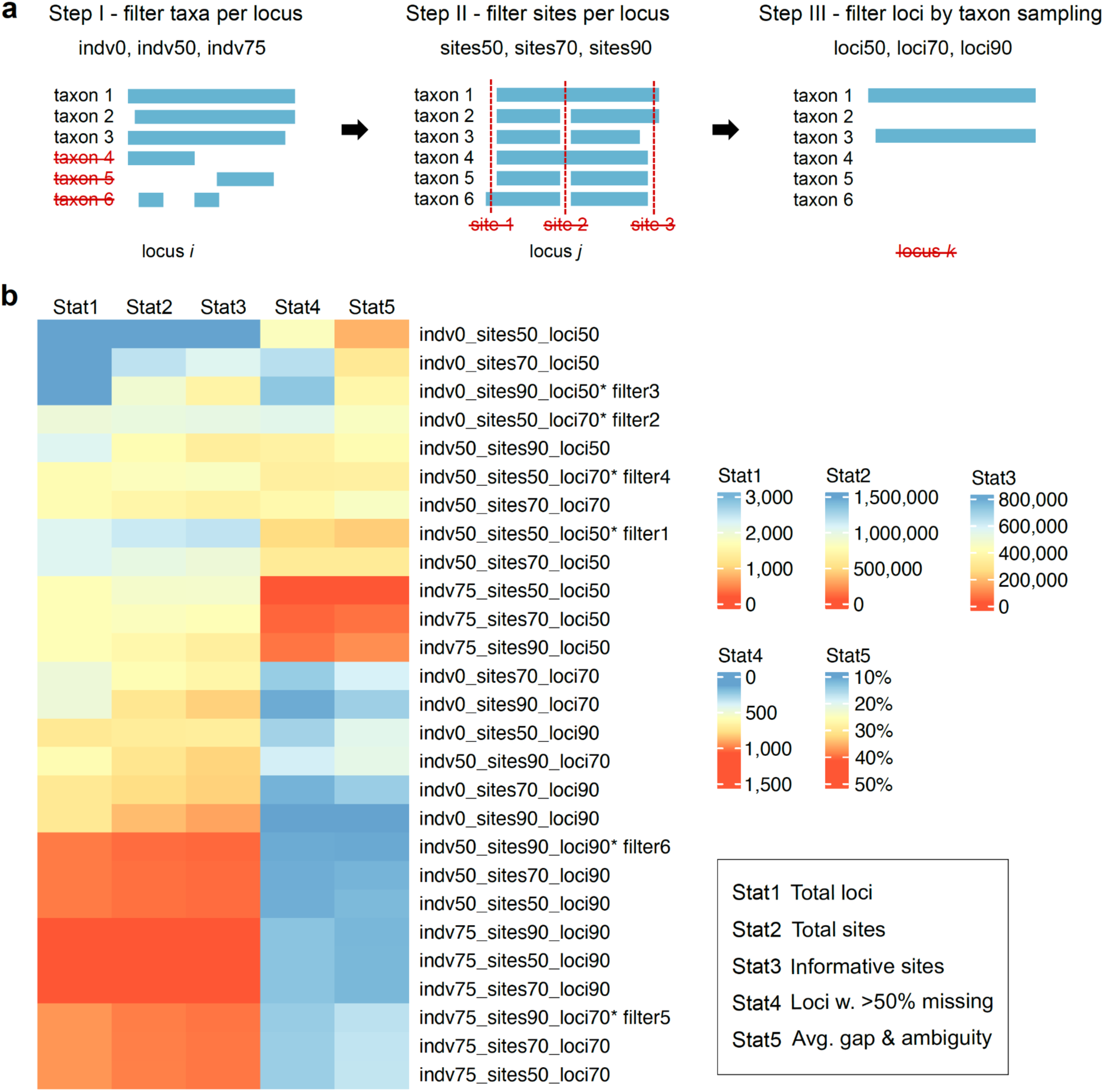
Filtering schemes and information content of different datasets. a). We used a combination of three strategies to filter datasets. Step I. Indv refers to the removal of individual taxa with short sequences for specific loci. indv0 indicates that we did not conduct this filtering step; indv50 and indv75 indicate that sequences shorter than 50% and 75% of the longest sequence for that alignment were removed. Step II. Sites refers to the trimming of sites dominated by gaps and missing data. sites50 indicates that alignment columns where ≥50% of taxa were gaps or missing are removed; sites70 and sites90 removed columns with ≥70% or ≥90% gaps or missing data, respectively. The percentage of taxa with gaps or missing data in a column reflects the number of taxa sampled for the locus of interest. Step III. Loci refers to the removal of poorly sampled loci. loci50, loci70, and loci90 indicate that loci are retained only if they are sampled for ≥50%, ≥70%, and ≥90% (respectively) of taxa in the full data matrix. b). Summary statistics (total number of loci, total number of sites, total number of parsimony informative sites, loci with > 50% data missing, and average proportion of gaps and ambiguities [“-”, “?” and “N”] across all loci) of the sequence alignments in all 27 filtered datasets. For missing data information (Stat4 and Stat5), hotter colors represent more missing data.

For each filtered dataset and the full dataset, we averaged the above individual-based summary statistics across all taxa sampled in that dataset (Supplementary Table S2). To visually inspect if taxa were clustering by study, we performed PCA using FactoMineR v1.34 (Lê et al. 2008) and plotted the first two principal components using ggplot2 v3.3.5.9 (Wickham 2011). We also used IQ-TREE2 (Nguyen et al. 2015) to compute locus-based summary statistics for each filtered dataset, i.e., number of loci, total sites, parsimony informative sites, average gap and ambiguity across all loci, and loci with more than 50% missing data (Supplementary Table S3).

### Initial data exploration

#### Concatenated analyses

We used the message passing interface (MPI) version of RAxML-NG v1.0.1 (Kozlov et al. 2019) to infer a ML phylogeny of the concatenated, full dataset (Table 1, baseline). Because this dataset was large, we ran two concurrent ML analyses that each used 800 compute cores – both used the GTR+R4 site rate substitution model, but one used parsimony to generate starting trees while the other used random starting trees. Because of the compute hours allocated to this project, we were only able to infer seven ML phylogenies using random starting trees and five ML phylogenies using parsimony starting trees. We selected the optimal tree as the one having the lowest log-likelihood across the 12 analyses. We generated support values for the full dataset by performing ML analysis on 10 standard (Felsenstein 1985) bootstrap replicates with the GTR+R4 model. Although the number of bootstrap replicates was limited because of compute hours allocated to this project, we evaluated the 10 bootstrap replicates for convergence using the --bs-converge option. We found that these replicates had converged, and we reconciled the “best” ML tree with the bootstrap replicates using RAxML-NG.

**Table 1.**
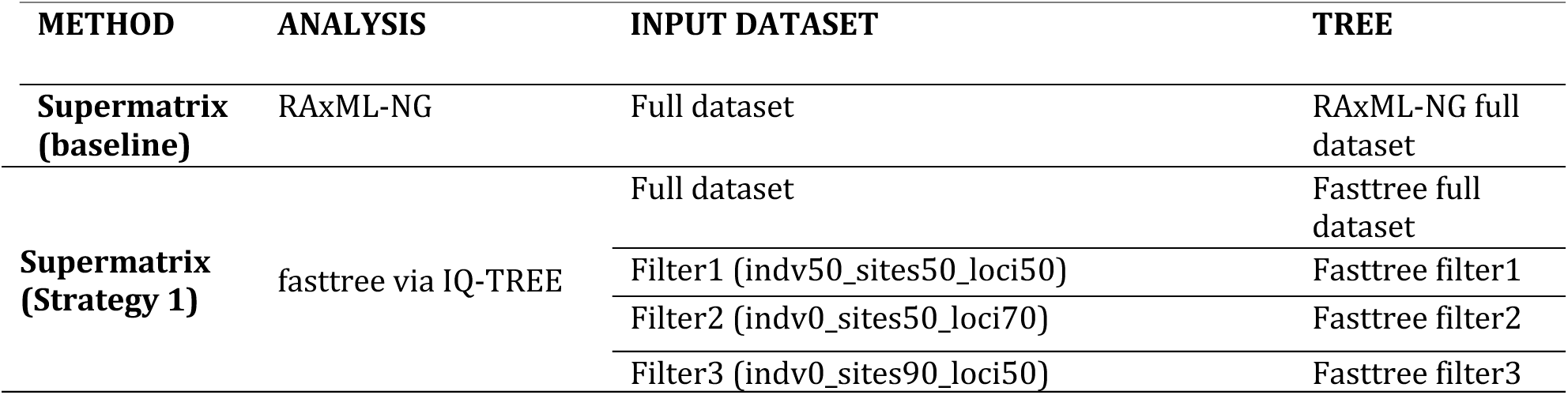
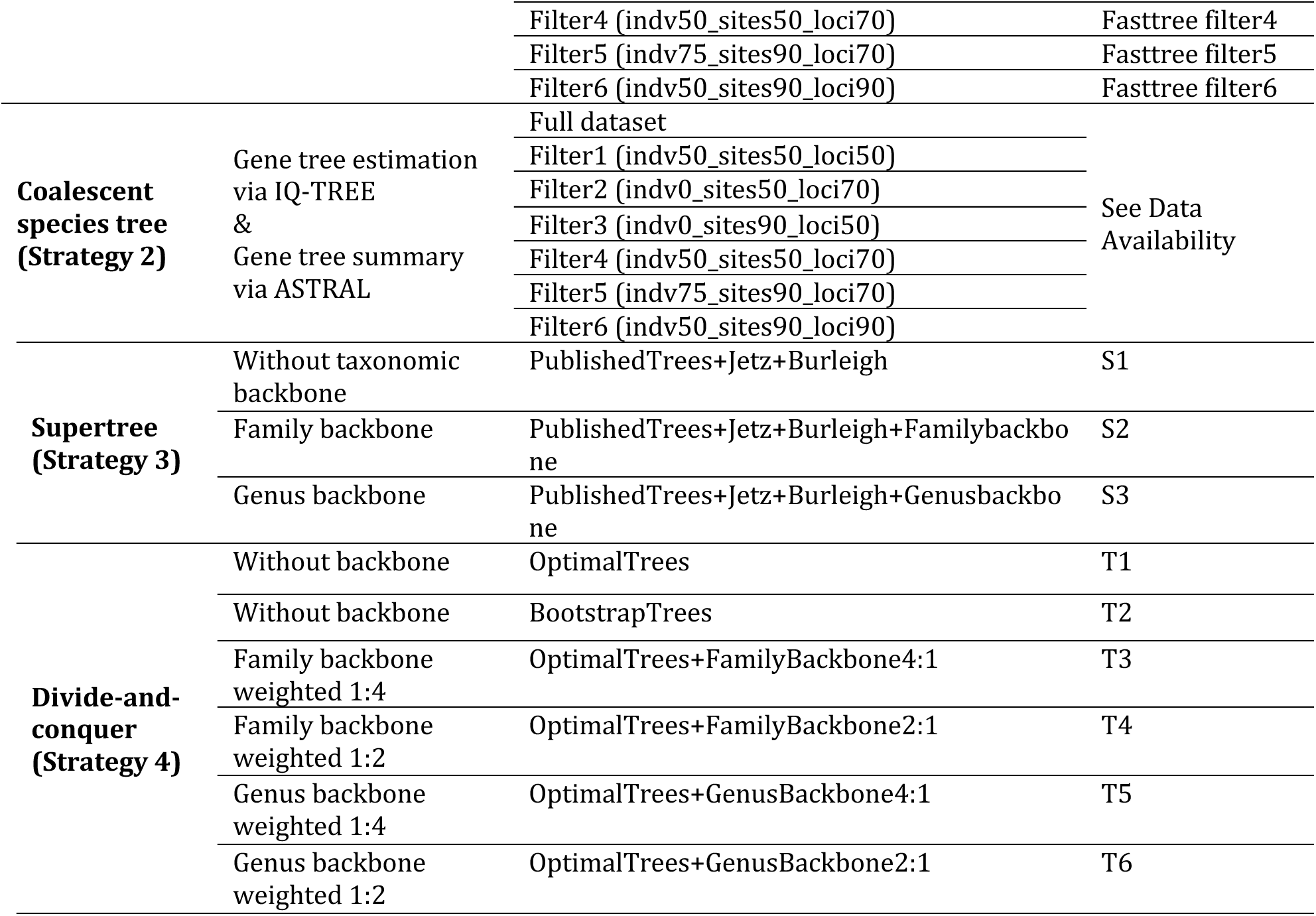
Summary of datasets, analyses, and phylogenetic trees in our initial exploration. The six filtered datasets were numbered based on total number of sites; filter1 with the highest number of sites and filter6 with the lowest number of sites.

To explore a faster method for ML tree estimation, we used the -fast option in IQ-TREE v2.0.5 (Nguyen et al. 2015) with the GTR+G site rate substitution model (Table 1, strategy 1). We initially inferred phylogenies from the concatenated, full dataset along with six filtered datasets that varied in numbers of loci, informative sites, and amounts of missing data. Similar to FastTree (Price et al. 2010), this inference mode estimates two starting trees (under BIONJ (Gascuel 1997) and MP) and then optimizes the trees by nearest neighbor interchange (NNI). This “fasttree” approach uses both parsimony and neighbor-joining starting trees, rapid hill climbing including stochastic nearest neighbor interchanges, and increased tolerance on likelihood values to speed up optimization, which has the potential to reduce accuracy.

Following the inference of trees from concatenated datasets, we performed an initial quality check of the inferred phylogenies by visual assessment of the relationships, and we pruned *Muscipipra vetula* and *Spheniscus mendiculus* from trees using the drop.tip function in ape v5.7-1 (Paradis and Schliep 2019) because these appeared in positions that were unlikely.

### Coalescent species tree estimation

For the full dataset and each of the six filtered datasets, we estimated individual gene trees using IQ-TREE v2.1.3 (Minh et al. 2020) under the GTR+G model, and we combined the ML trees to generate a species tree using ASTRAL v5.7.8 (Zhang et al. 2018) (Table 1, strategy 2).

### Building supertrees using existing phylogenomic trees

Supertree methods (Table 1, strategy 3) infer phylogenies from existing trees, and we identified 53 trees from 46 phylogenomic studies (McCormack, Harvey, et al. 2013; Jarvis et al. 2014; Lamichhaney et al. 2015; Nater et al. 2015; Prum et al. 2015; Bryson et al. 2016; Hosner et al. 2016; Manthey et al. 2016; Ottenburghs et al. 2016; Zarza et al. 2016; Burga et al. 2017; Reddy et al. 2017; Wang et al. 2017; White et al. 2017; Yonezawa et al. 2017; Andersen et al. 2018; Bruxaux et al. 2018; Campillo et al. 2018; Chen et al. 2018; Ferreira et al. 2018; Musher and Cracraft 2018; Younger et al. 2018; Andermann et al. 2019; Andersen et al. 2019; Everson et al. 2019; Jenna M. McCullough et al. 2019; Jenna M McCullough et al. 2019; Oliveros et al. 2019; Sackton et al. 2019; White and Braun 2019; Harvey et al. 2020; Imfeld et al. 2020; Oliveros et al. 2020; Salter et al. 2020; Smith et al. 2020; Vianna et al. 2020; Catanach et al. 2021; Kirchman et al. 2021; Oliveros et al. 2021; McCullough et al. 2022; Vinay et al. 2022; Wang et al. 2022; Smith et al. 2023; Zhao et al. 2023; Braun et al. 2024; for details, see Supplementary Table S4) having taxon samples that overlapped with the species included in the supermatrix datasets. We downloaded tree files representing the phylogenies generated as part of these studies from data repositories. For those that were not publicly available, we requested the tree files from the authors of the study. If we did not receive a tree file from the authors and the tree contained a small number of taxa, we manually created a tree file in NEWICK format to reflect the topology shown in the publication. After obtaining or creating tree files representing all studies, we reconciled the taxon names to match those in the IOC World Bird List v13.1 (Gill et al. 2023) and pruned duplicate tips that represented the same species within a tree using the drop.tip function in ape.

Because the phylogenomic trees we downloaded included few taxa that overlapped among studies, we integrated them using three types of backbone trees: one from Burleigh et al. (2015) that we refer to as the “Burleigh backbone”, a second from Jetz et al. (2012) that we refer to as the “Jetz backbone”, and a third “taxonomic” backbone. To generate the Burleigh backbone, we downloaded 1000 ML bootstrap trees from Dryad (https://doi.org/10.5061/dryad.r6b87) and produced a majority rule consensus of these using SumTrees (Sukumaran and Holder 2010). Similarly, for the Jetz backbone, we downloaded 1000 “Hackett” (Hackett et al. 2008) backbone trees from birdtree.org (1- 1000, accessed October 24, 2022) and produced a majority rule consensus using SumTrees. Then, we manually reconciled taxon names throughout the Burleigh and Jetz backbone trees to match the IOC World Bird List v13.1 taxonomy, and we trimmed duplicated tips from each tree using the drop.tip function of ape. Finally, we created two taxonomic backbone trees (a family-level backbone and a genus-level backbone) to reflect recent taxonomic revisions in the IOC World Bird List v13.1. We began by extracting and combining the taxon names from the Burleigh backbone (6714 tips), the Jetz backbone (9979 tips), and the 53 trees we identified above, which totalled 10,383 taxa after we removed duplicate names using drop.tip function of ape. We created the family-level taxonomic backbone using the IOC World Bird List v13.1 to: group individual taxa by family, cluster taxa from same family into a polytomy, cluster families from the same order into a polytomy, and cluster orders into infraclasses Palaeognathae, Galloanserae, and Neoaves. Finally, we enforced a tree topology to reflect (outgroup,(Palaeognathae,(Galloanserae,Neoaves))), because these relationships among infraclasses are well-established (e.g., Jarvis et al. 2014; Burleigh et al. 2015; Prum et al. 2015; Stiller et al. 2024). We constructed the genus-level taxonomic backbone similarly by using the IOC World Bird List v13.1 to cluster taxa from the same genus into a polytomy, then clustering by family, order, and infraclass and enforcing the same topology among infraclasses.

We used matrix representation with parsimony (MRP) (Baum 1992; Ragan 1992) to generate supertrees following the pipeline described in Kimball et al. (Kimball et al. 2019). Since the supertree method can suffer from source tree incongruence (Bininda-Emonds et al. 2002), we employed a user-guided weighting scheme to address topological conflicts among source trees. Specifically, we assigned different weights to input trees based on the amount of data used to infer them (Supplementary Table S4) by including from one (low weight) to eight (high weight) copies in the supertree matrix. For example, trees that were based on whole-genome sequencing data, such as the Jarvis TENT tree (Jarvis et al. 2014), were given a weight of eight and included in the supertree matrix eight times. We typically weighted UCE trees as four. However, if a study included two UCE trees estimated by different approaches (e.g., methods of tree estimation or filtering strategies) but using completely or largely overlapping data, we assigned each tree a weight of two. We assigned two additional trees (Reddy et al. 2017; Yonezawa et al. 2017) a weight of two because they were based on a large number of “legacy markers” (Kimball et al. 2009) extracted from genome assemblies. Finally, we assigned a weight of one to all backbone trees.

After determining the weighting scheme, we created three supertree matrices: 1) weighted trees with the Burleigh and Jetz backbones; 2) weighted trees with Burleigh, Jetz, and family-level taxonomic backbones; and 3) weighted trees with Burleigh, Jetz, and genus-level taxonomic backbones. Then we used CLANN (Creevey and McInerney 2005) to convert the input tree matrix to a binary (MRP) representation and generated supertrees using PAUP* v4.0 (Swofford 2003). We conducted the searches using the parsimony ratchet (Nixon 1999) as described in Kimball et al. (2019), who used code available from https://github.com/ebraun68/ratchblock to generate PAUP blocks that ran five tree searches with different upweighting scores (15%, 20%, 25%, 30%, and 35%). Each tree search consisted of 100 replicates and produced a strict consensus tree from these replicates after the tree search concluded. For each of the three matrices, we selected the resulting supertree as the one from the five searches that had the best parsimony score. Then we pruned the resulting three supertrees to include only the taxa present in the full (supermatrix) dataset, which resulted in 2751 taxa (seven taxa in our supermatrix were not included in published phylogenies).

### Building supertrees using a divide-and-conquer approach

Because supermatrix methods can be computationally intensive when applied to large datasets, we wanted to test a divide-and-conquer approach that combined supermatrix and supertree methods by dividing the supermatrix into subsets of taxa, inferring trees from each subset using supermatrix methods, then integrating the resulting subset trees with supertree methods (Table 1, strategy 4). To begin the process, we designed three subsetting schemes that differed in the likely number of overlapping taxa shared between them: random subsets, partially stratified subsets, and fully stratified subsets. Then we created 15 random subsets by randomly drawing (with replacement) 150 taxa from the total list of taxa in the full dataset.

To create the partially stratified subsets, we stratified the 2,760 taxa in the full dataset into six major groups (recovered across many studies) based on our taxon sampling (Supplementary Figure S1). Then, we randomly selected 7.5%, 3.1%, 7.8%, 6.8%, 7.4%, and 8.0% of the taxa within each group largely based on its size while avoiding oversampling suboscines, which produced a partially stratified subset of 150 taxa. We repeated this selection process without replacement to create a total of 10 partially stratified subsets.

To create the fully stratified subsets, we stratified all taxa in the full dataset into 25 groups (Supplementary Figure S2) that were based on taxonomy to ensure all taxa were represented at least once across the subsets and were included in trees with congeners (so sister relationships could hopefully be resolved). We set the number of taxa included in each subset under 200 to maximize computational efficiency given our resources (see Supplementary information). Because supertree analyses require overlapping taxa, we then manually selected “linker taxa” from outside each group and included them in the group membership. Initial phylogenetic analyses suggested that using identical linker taxa across fully stratified subsets placed the linker taxa in unexpected positions in the resulting tree. Therefore, we used distinct linker taxa for each subset, which resolved this issue.

We created a total of 50 subsets across all schemes. We extracted subset alignments from the aligned, concatenated, full dataset. Then, we used IQ-TREE v2.1.3 (Minh et al. 2020) to infer the “best” ML phylogenies and generate 1000 ultrafast bootstrap replicates for each subset using the GTR+R4 site rate substitution model.

We followed the same weighted-tree search approach described above to infer a set of supertrees representing all taxa from the 50 “best” ML subtrees. Specifically, we created five supertree matrices using: 1) the 50 best ML subtrees where each tree was given a weight (w) of one (w = 1); 2) the 50 best ML subtrees (w = 4) and the family-level backbone tree (w = 1); 3) the 50 best ML subtrees (w = 4) and the family-level backbone tree (w = 1); 4) the 50 best ML subtrees (w = 2) and the genus-level backbone tree (w = 1); and 5) the 50 best ML subtrees (w = 2) and the genus-level backbone tree (w = 1).

We also built 1000 MRP matrices (each with 50 trees) from the bootstrap replicates by sampling and combining replicates from the subsets in the order they were generated: bootstrap replicate tree one from all 50 subsets combined to form MRP matrix one, bootstrap replicate tree two from all 50 subsets combined to form MRP matrix two, et cetera. Then we performed the tree search process described above for each MRP matrix to produce a set of 1000 phylogenomic supertrees that we summarized to a 50% majority rule consensus using SumTrees. We pruned the six supertrees generated from the steps above to include only the taxa present in the full (supermatrix) dataset.

### Analyzing tree distances

To visually represent differences between the various trees we inferred, we rooted trees on the crocodilian outgroup and used ete3 (Huerta-Cepas et al. 2016) to calculate pairwise normalized Robinson-Foulds distances between the two trees inferred from the full dataset, the six trees inferred from the filtered datasets, the three trees inferred from the supermatrix analyses, and the six trees inferred using the divide-and-conquer approach. We used the write.nexus.dist function in phangorn v2.11.1 (Schliep 2011) to create a NEXUS block of the pairwise Robinson-Foulds distances, and we used PAUP* v4.0 (Swofford 2003) to infer a neighbor-joining (NJ) “tree-of-trees” that we rooted at the midpoint.

### Testing for clade monophyly

Unlike many larger taxonomic groups, the avian phylogeny has been extensively studied, resulting in strongly corroborated clades across multiple analyses (reviewed in Braun et al. (2024)). Sangster et al. (2022) and earlier work (Chen and Field 2020; Queiroz et al. 2020; Sangster and Mayr 2021) highlighted a number of clades near the base of the bird tree that are very likely to be present in the true avian species tree. Modern taxonomies (e.g., IOC (Gill et al. 2023), eBird/Clements (Clements et al. 2023), Howard & Moore (Dickinson and Christidis 2014)) now circumscribe orders, families, and genera in a manner that is likely to be correct (though some families and genera continue to be refined as more information becomes available). Although there are almost certainly some named taxa that do not represent clades in the true bird tree, the majority of named groups are likely to be expected clades. We were interested in comparing how reliably the different tree inference methods resolved these expected clades across the avian phylogeny. These include clades that represent different taxonomic groups recognized by the IOC World Bird List v13.1, as well as 33 high-level clades that have been found in multiple phylogenomic studies (Sangster et al. 2022). We generally assumed that a method was more reliable when it recovered a larger number of these groups as monophyletic. To perform these analyses, we first excluded clades that were only represented by a single species. Then we used the AssessMonophyly function in MonoPhy (Schwery and O’Meara 2016) to calculate how many genera, families, orders, and 33 expected high-level clades were not resolved as monophyletic.

### Summarizing compute time

Because searches over tree space including thousands of taxa are computationally demanding and require significant amounts of energy (Grealey et al. 2022; Kumar 2022), we were interested in summarizing and comparing the compute time required for the tree inferences described above. For this study, analyses were run across several different computing systems: HPC@LSU (RAxML-NG analysis), AMNH HPC (initial fasttree analyses and ASTRAL analyses), and UF HiPerGator (supertree and divide-and-conquer analyses). For each system, we tallied and separated the CPU hours spent for tree searches versus bootstrap replicate searches, if applicable, and we collected the total cluster utilization for each SLURM job. To determine the time required for the RAxML-NG analysis of the full, concatenated dataset, we combined the CPU time for the random and the parsimony starting trees. For the divide-and-conquer analyses, we summed the CPU hours spent for tree search across 50 subsets. Because the supertree component for the divide-and-conquer analyses used very little CPU time compared to the subset concatenation analysis, we added it directly to the total CPU time spent (for the bootstrap trees, time for 1000 runs were added). For the regular supertree analyses, we presented the PAUP tree search time and added time for MRP matrix construction to the total CPU hours spent.

### Tests on two filtered datasets

To improve fasttree search and optimization, we examined the role of the starting tree using two filtered datasets (filter1 and filter3). We chose these filter sets due to their contrasting patterns of expected clade recovery in initial exploration: filter1 performed well deeper in the tree but poorly at the tips (only one expected high-level clade was non-monophyletic, but 41 genera were non-monophyletic), whereas filter3 showed the opposite pattern (eight high-level clades and 37 genera were non-monophyletic). Fasttree searches normally use two starting trees (MP and BIONJ), however, users can supply their own starting tree to bypass the default starting tree estimation process. We generated a total of 24 additional tree searches using two filtered datasets (filter1 and filter3) with different starting trees: 1) MP trees generated by Parsimonator v1.0.2 (https://github.com/stamatak/Parsimonator-1.0.2) using the same filtered dataset as the search; 2) MP trees generated by Parsimonator using different filtered datasets (excluding filter5 and filter6 as these two datasets performed poorly in estimating accurate phylogenies); 3) greedy consensus tree of the MP starting trees of each dataset in previous steps; 4) greedy consensus tree of initial fasttrees (fulldata and filter1–filter4); 5) BIONJ starting tree using the same filtered dataset as the search; 6) BIONJ starting tree using filter2; 7) the ML tree search from the initial analysis; and 8) the RAxML-NG tree. We also reran some of the analyses using alternative site models (GTR+G vs. GTR+R4). We evaluated the log likelihoods of the starting tree and optimal tree and assessed expected clade recovery for the final ML tree in each analysis.

Searches initiated with BIONJ starting trees required a large amount of time, had a much lower likelihood, and resulted in worse expected clade recovery than the initial tree (Supplementary Figure S3 and Supplementary Table S5). Next, we tested MP starting trees generated from four relatively large datasets (filter1-4). Fasttrees built using MP starting trees derived from the same filtered dataset used for the ML search consistently had much better likelihoods than those derived from other filtered datasets. However, in some cases, starting trees based on other filtered datasets produced better expected clade recovery (Supplementary Figure S3). These results suggest a straightforward method to improve the speed and reproducibility of fasttree searches: avoid generating the BIONJ tree and instead conduct multiple searches using MP starting trees generated from the same dataset used for the fasttree search.

### New fasttree method

Based on results from the tests on filter1 and filter3, we found that conducting multiple fasttree searches using MP starting trees generated using the same dataset has the potential to improve the fasttree analysis. Therefore, we used Parsimonator v1.0.2 to estimate four MP starting trees (parsA, parsB, parsC, and parsD) for each of the full and 27 filtered datasets (with *Muscipipra vetula* and *Spheniscus mendiculus* removed from supermatrices). Each MP starting tree was used to run a fasttree analysis in IQ-TREE v2.2.2 (Nguyen et al. 2015) with parsA and parsB using GTR+G model and parsC and parsD using FreeRates model (GTR+R4). There are two filtered datasets that are identical to each other (indv75_sites50_loci90 and indv75_sites70_loci90), therefore only one set of analyses was done for these two datasets. This resulted in a total of 108 new fasttrees, four for each dataset (Table 2). We evaluated their performance in clade recovery and summarized the total CPU time spent.

**Table 2.**
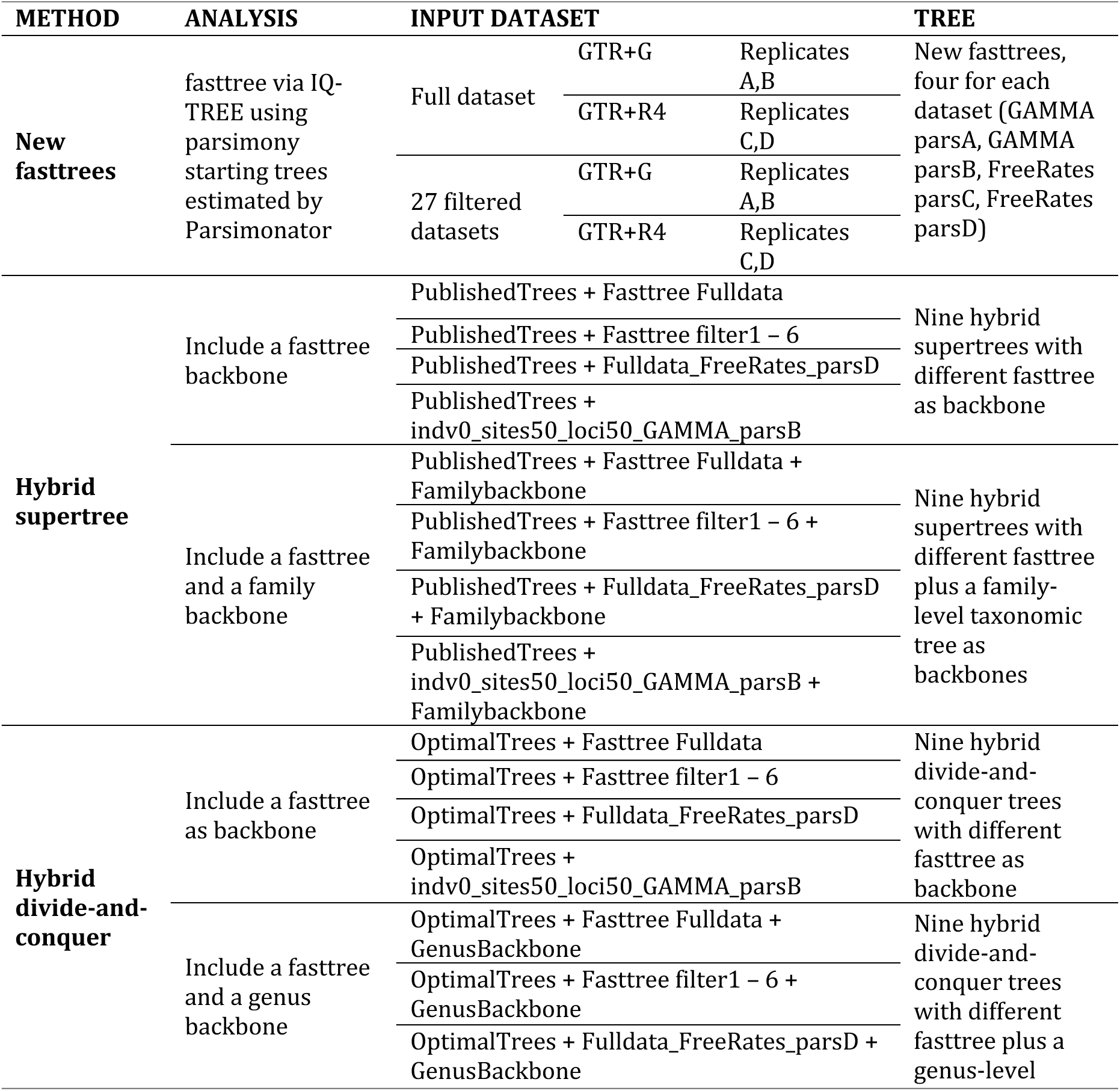
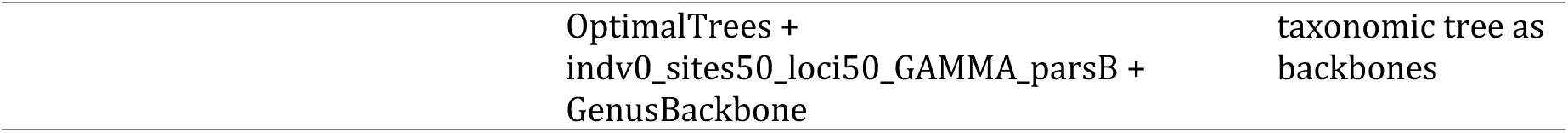
Summary of datasets, analyses, and phylogenetic trees using modified methods.

We used ComplexHeatmap (Gu 2022) in R (R Core Team 2023) to plot 27 filtered datasets using number of loci, total sites, parsimony informative sites, average proportion of gap or ambiguity across all loci, and loci with more than 50% missing data. We also created a version with clusters using z-transformed statistics (Supplementary Figure S4). From each cluster, we selected a representative dataset that performed best in recovering expected clades (Supplementary Table S6). We only presented the best fasttree for these representative datasets in the main text.

### Hybrid approaches

We tested whether fasttrees could improve the supertree and divide-and-conquer methods when used as backbone trees. Unlike the Jetz+Burleigh backbones used initially, our fasttrees included all taxa in the analyses, potentially providing a better backbone to compensate for limited overlap among source trees. Additionally, because our fasttrees were estimated from phylogenomic data, they may offer a more accurate representation of relationships, potentially reducing the need for taxonomic backbones. We referred to these new approaches as the “hybrid supertree approach” and “hybrid divide-and-conquer approach” (Table 2).

We used the two best new fasttrees (based on expected clade recovery) and seven initial fasttrees as the backbone tree in supertree and divide-and-conquer analyses (Table 2). Two sets of analyses, each with nine trees, were estimated for the hybrid supertree approach, with different fasttree as the backbone with or without a family-level taxonomic backbone. Each backbone was given a weight of one, and source trees were given different weights based on the amount of data used to infer them, as described above. For the hybrid divide-and-conquer approach, we also ran two sets of analyses, each with nine trees estimated: 1) using only a fasttree as the backbone, the 50 best ML subtrees and the fasttree backbone were each given a weight of one; and 2) using a fasttree backbone and a genus-level backbone, the 50 best ML subtrees were each given a weight of two and the backbones were given a weight of one. We then followed the same steps as above to build a binary MRP tree matrix in CLANN and generate supertrees using PAUP*. Similarly, we evaluated the performance in expected clade recovery for final output trees. When summarizing the total CPU time spent, we also added in the compute time for generating each MP starting tree and the fasttree. All new fasttrees and hybrid approaches were run on UF HiPerGator.

### Molecular dating

We applied a total of 43 fossil calibrations for node-dating analyses (Supplementary Table S7) following best practices proposed by Parham et al. (2012), and we assigned minimum and maximum possible ages to each calibrated node in our phylogeny (that is, the last common ancestor shared by two given species). Additional information regarding the fossils selected to calibrate divergence time analyses is presented in the Supplementary Information.

Then, due to the size of the resulting trees, we used TreePL (Smith and O’Meara 2012) to estimate divergence times for the (1) RAxML-NG tree inferred from the concatenated, full dataset; (2) two fasttrees using new fasttree methods based on the full dataset and the filtered dataset indiv0_sites50_loci50; 3) two supertrees (one from initial exploration and one from hybrid approach); and 4) two divide-and-conquer trees (one from initial exploration and one from hybrid approach). For the four supertrees and divide-and-conquer trees, we used IQ-TREE2 v.2.2.2 (Nguyen et al. 2015) to optimize the tree branch lengths (--tree-fix) under both GTR+G and GTR+R4 model using the filtered dataset with the smallest amount of missing data (indv0_sites90_loci90). TreePL allows for varying rates across branches but penalizes rate differences over the tree with a rate smoothing parameter, so we identified the optimal rate smoothing parameter through cross-validation that tested 10 values (start = 1e-07; stop = 10000). We also used the “prime” option to identify the best optimization parameters and the “thorough” option to allow the program to iterate until convergence.

For each of the seven resulting chronograms, we extracted divergence times for 12 major groups that have been consistently resolved across studies and that represent both ancient and recently diverged clades as well as both fast- and slow-evolving clades, and we compared these divergence estimates to those in other studies (Claramunt and Cracraft 2015; Prum et al. 2015; Kimball et al. 2019; Kuhl et al. 2021; Brocklehurst and Field 2024; Claramunt et al. 2024; Stiller et al. 2024; Wu et al. 2024a). To extract the divergence time for each of these clades, we defined a species pair that shares the MRCA of the clade and used the fastDist function with phytools (Revell 2012) to compute half of the patristic distance between the two species in each chronogram. Divergences estimated under GTR+G and GTR+R4 models were very similar (see Data Availability), thus only results from GTR+R4 model were plotted. We also computed relative divergence time for these clades by scaling the divergences to Neognathae.

## Results

### Taxon sampling

We created an avian UCE data matrix by combining assembled sequences from 22 sequence capture studies with UCE sequences extracted from 125 genome assemblies available in NCBI (Supplementary Table S1). After standardizing taxonomies and removing duplicate taxa, we generated DNA sequence alignments for 5,121 target captured loci, with an average length of 665 base pairs (bp) and including a total of 2,047,980 parsimony informative sites. The full dataset contained 2,758 tips (including two crocodilian outgroups); members of all 44 extant bird orders and one extinct order (Dinornithiformes); 250 of 253 (98.8%) extant bird families and one extinct family (Emeidae); 1,081 genera; and 2,747 unique species (see Data Availability).

### Dataset characteristics and filtering

Data heterogeneity was evident in descriptive statistics for individual taxa. For instance, taxa showed considerable variation in locus count, sequence length, and individual-based parsimony informative sites both within and between studies (Supplementary Figure S5). PCA of these summary statistics revealed distinct clusters corresponding to their source datasets (Supplementary Figure S6). By combining three levels of filtering criteria, we generated 27 filtered datasets from the full supermatrix dataset to reduce heterogeneity related to taxa with short sequences, sites with gaps and missing data, and poorly sampled loci (Fig. 1a). As anticipated, more stringent filtering schemes substantially increased homogeneity among studies and reduced the amount of missing data. However, these improvements reduced the number of informative sites (Fig. 1b; Supplementary Figure S5).

### Baseline phylogeny

We generated a baseline phylogeny for comparison using RAxML-NG with the concatenated full dataset (Fig. 2; Supplementary Figure S7) and assessed tree quality using the expected clade recovery criterion. The RAxML-NG tree recovered all 33 high-level clades identified by Sangster et al. (2022), all 40 evaluated orders (excluding monotypic or single-sampled orders), all but two of the 138 evaluated families, and all but 38 of the 410 evaluated genera.

**Figure 2.**
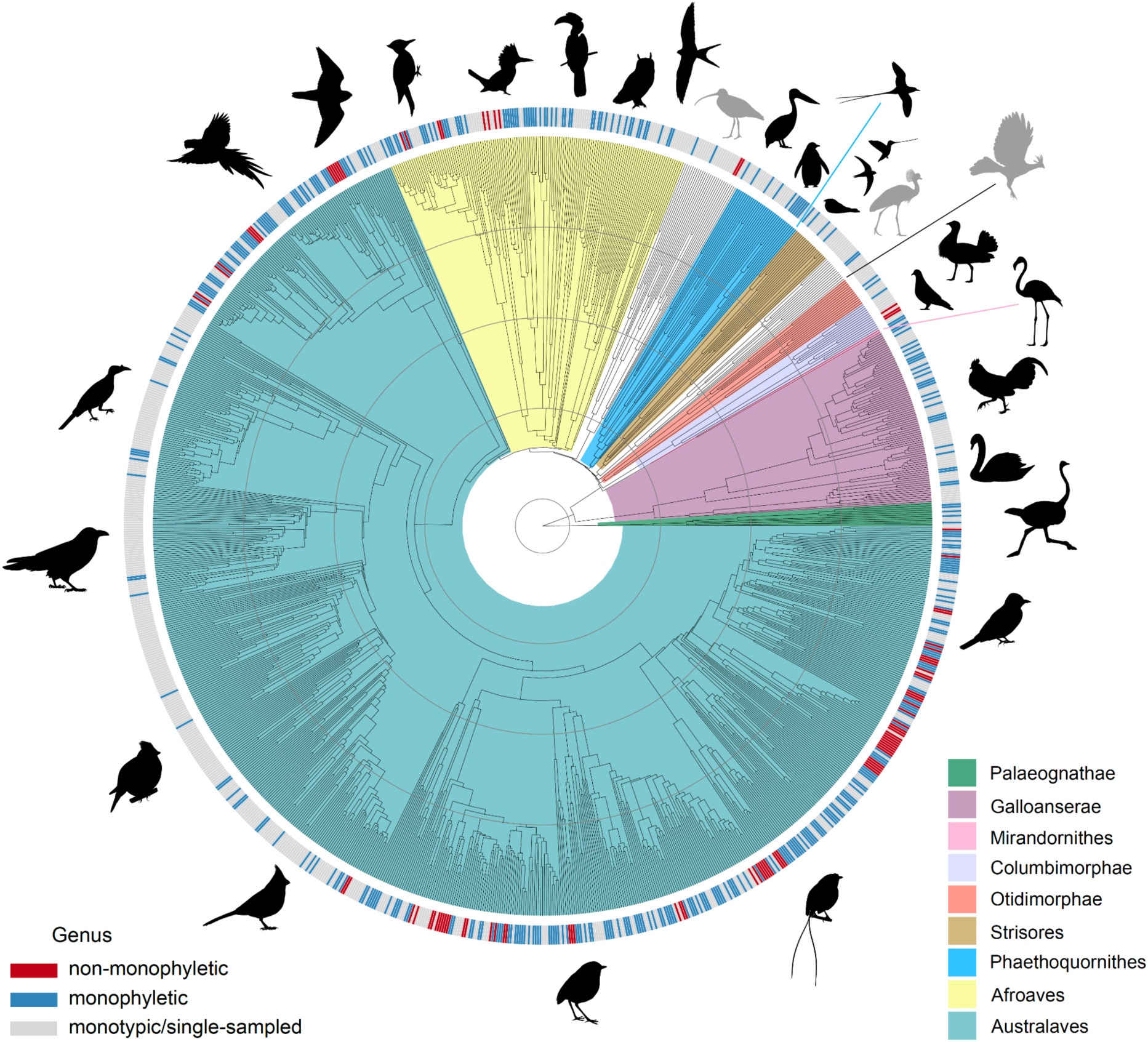
A genus-level RAxML-NG tree with branch lengths converted to divergence time using TreePL. Major bird clades are color-coded, while three lineages (Gruiformes, Charadriiformes and Opisthocomiformes; see Reddy et al. 2017) that were not placed within a strongly corroborated superordinal clade, remain uncolored (silhouettes in gray). Colored bars in the outer ring indicate genera that are monophyletic (blue; n = 372) and non-monophyletic (red; n = 38) in this phylogeny. Monotypic genera (n = 334 with a single species currently recognized in IOC World Bird List v13.1) and genera represented by a single sample in our dataset (n = 337) are gray. The concentric gray circles indicate 20 Ma time intervals. See Supplementary Figure S7 for a version of this tree with tip labels.

Although the RAxML-NG tree appeared to provide an accurate estimate of avian phylogeny based on expected clade recovery, generating this tree required significant computational resources – approximately 49 years of CPU time for the primary search and an additional 37 years of CPU time for a limited number of bootstrap analyses.

### Initial exploration

We explored four alternative approaches (Table 1) that were more computationally efficient than standard ML: (1) implementing a fast ML estimation approach, (2) estimating individual gene trees and combining them into a species tree using ASTRAL, (3) combining source trees into a supertree, and (4) using a divide-and-conquer strategy in which trees were estimated from taxonomic subsets of the supermatrix and then combined into a supertree. The primary goal of these analyses was to determine whether any of these alternative methods could produce trees that were at least as accurate as the RAxML-NG tree while substantially reducing computational requirements.

We explored whether fast ML estimation (Table 1, strategy 1) could produce a tree with similar recovery of expected clades to the RAxML-NG analysis. While the fasttree estimated from the full supermatrix required less computation time, its recovery of expected clades was much lower. The resulting tree did not perform as well as either the RAxML-NG tree or the best trees from other approaches (Fig. 3). Filtering appeared to improve the performance of fasttree analyses, with the best results based on the expected clade recovery criterion observed in trees inferred from the least aggressively filtered datasets (filter1 and filter2). By contrast, the most aggressively filtered datasets (filter5 and filter6) performed poorly in the clade recovery criterion compared to the RAxML-NG tree and the better-performing filter1 and filter2 fasttrees. Moreover, the clade recovery of filter5 and filter6 fasttrees was similar to that of the full data fasttree, suggesting diminishing returns with overly stringent filtering.

**Figure 3.**
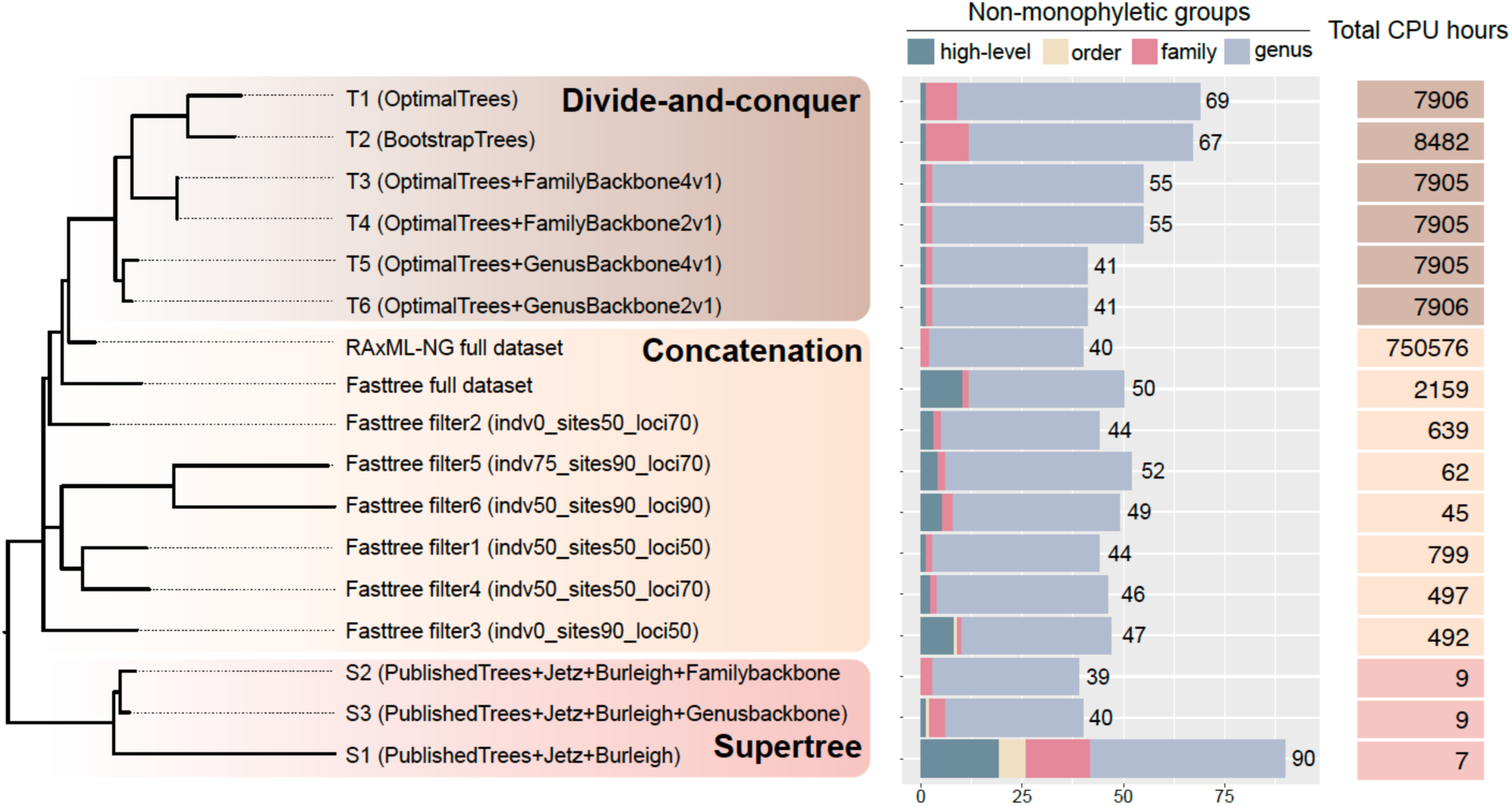
Distribution of trees from initial exploration in tree space built with neighbor-joining using normalized Robinson-Foulds distances (rooted at the midpoint). For each tree, we summarized the number of high-level clades, orders, families, and genera recognized by IOC World Bird List v13.1 that are not monophyletic in the tree; therefore, the higher the number, the more non-monophyletic groups. Non-monophyly may be due to artifacts in phylogenetic inference or taxonomic classification that requires revision. We also summarized the total CPU time used for generating each tree. For the divide-and-conquer approach, we show the tree search CPU time for concatenation analyses, and the time for supertree search was directly added to the total CPU time spent.

The ASTRAL species trees (Table 1, strategy 2) recovered substantially fewer expected clades than either the RAxML-NG tree or the fasttrees, regardless of the filtering procedure (or lack thereof) used to generate the alignments for gene tree estimation. The total number of unresolved groups ranged from 144 to 207 and computation time ranged from 1,026 to 68,663 CPU hours (Supplementary Table S8).

For the supertree analysis (Table 1; strategy 3), the supertree constructed without taxonomic backbones (S1) performed poorly in recovering expected clades (Fig. 3). In contrast, the two supertrees with taxonomic backbones (S2 & S3) performed as well as, or slightly better than the RAxML-NG tree in terms of expected clade recovery while still requiring minimal computation time (Fig. 3).

The divide-and-conquer approach (Table 1, strategy 4) without taxonomic backbones outperformed the supertree without backbones in recovering expected clades (Fig. 3).

However, performance comparable to the RAxML-NG tree was achieved only when a genus backbone was included. Despite requiring the estimation of input trees from the supermatrix, this method was computationally efficient, using approximately 1% of the total compute time required for the full RAxML-NG analysis (Fig. 3).

The two divide-and-conquer trees using the genus backbone (T5 & T6) performed well overall but exhibited polytomies within heavily sampled families, such as Tyrannidae and Thamnophilidae, as well as among some oscine families. Notably, these polytomies were not observed in Oliveros et al. (2019) and Harvey et al. (2020), which were the sources of most of the passerine data. The number of polytomies decreased when the weight of the source trees relative to the genus backbone was reduced (lower in T6 [2:1] versus higher in T5 [4:1]; see Supplementary Information for details on comparing polytomies). However, this adjustment did not affect the recovery of expected clades.

These phylogenetic results (excluding ASTRAL trees) are summarized by a NJ tree based on pairwise normalized Robinson-Foulds distances between alternative topologies (Fig. 3). This analysis indicated that the method of inference (supermatrix, supertree, or divide-and-conquer) strongly influenced topological similarity. Notably, supertree and divide-and-conquer methods formed distinct clusters. For the supertrees, this clustering may reflect biases introduced by relationships within the source trees, which differed from those inferred using other methods. Similarly, the clustering of divide-and-conquer analyses likely stems from the use of the same underlying subset trees (or their bootstrap consensus), which may have contributed unique relationships within the data subsets. By contrast, the fasttrees did not form a single cluster, and branch lengths in the NJ tree indicated greater variation among these analyses compared to the other methods. This increased variation is expected, given that the fasttree datasets differed in content due to filtering.

### New fasttrees

We conducted four searches on the full dataset and each of the 27 filtered datasets. Analysis of expected clade recovery for all new fasttrees (Supplementary Table S6) revealed that one fasttree from the full dataset (using an MP starting tree with the GTR+R4 model in replicate search D, i.e., FreeRates parsD) matched the RAxML-NG tree in both the number and identity of expected clades (Figs. 4 & 5). This best full data fasttree not only closely approximated the RAxML-NG tree in tree space (Fig. 4), but it was also far more computationally efficient (198-fold difference in the number of CPU hours between the two analyses).

**Figure 4.**
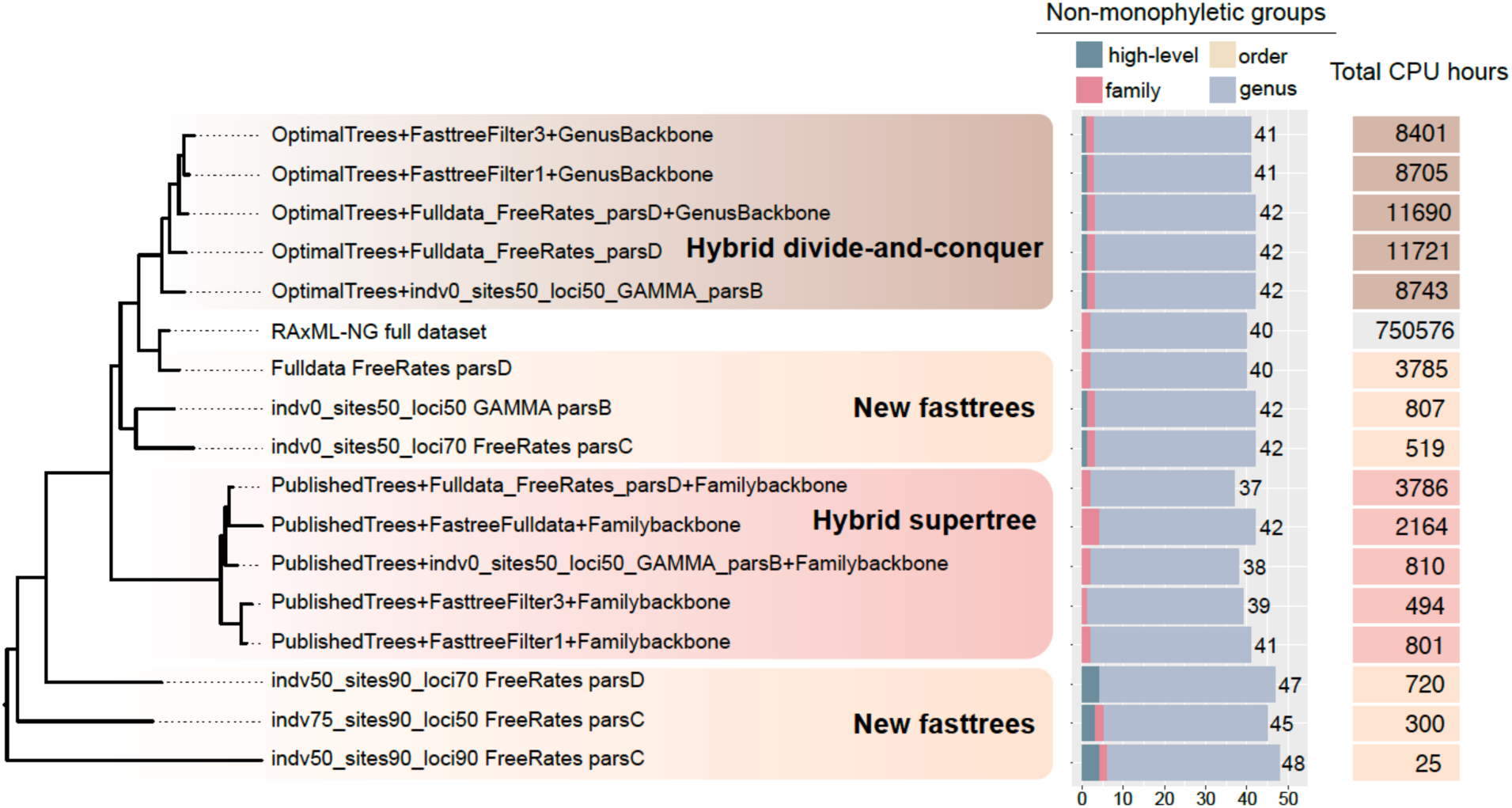
Neighbor-joining tree of trees based on normalized Robinson-Foulds distances (rooted at the midpoint). Here we only present the representative trees for each approach. Full results can be found in Supplementary Tables S7-9. For the two hybrid approaches, we added in the time for running the backbone tree. For the new fasttrees, we summarized the compute time for running four fasttrees analyses for each dataset (using different MP starting trees under GTR+G [parsA & parsB] or GTR+R4 [parsC & parsD] model) and presented the total time used here. For example, for a new fasttree analysis based on the full dataset, four runs each cost 999, 881, 961, and 943 CPU hours; we presented the fasttree parsD here, but the total CPU hours spent was a sum of the four runs (i.e., 3785).

**Figure 5.**
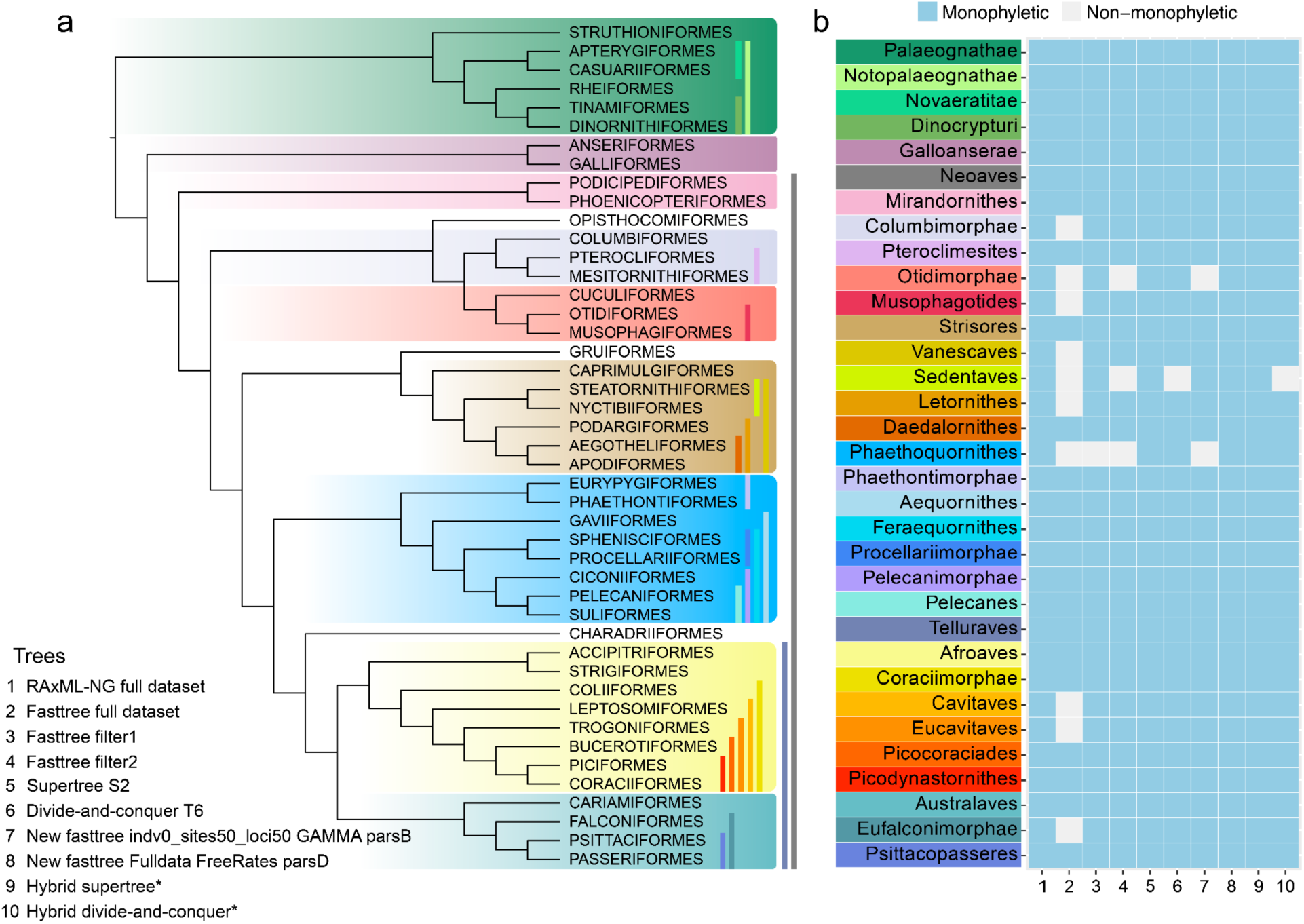
a. Cladogram showing relationships among orders in the best fasttree using the full concatenated dataset and new fasttree approach (tree No. 8). Vertical bars next to order names indicate composition of superordinal clades. b. We compared recovery of superordinal clades in trees estimated from the full dataset (both by RAxML-NG and initial fasttree analysis via IQ-TREE) and trees with the best expected clade recovery using various approaches: two initial fasttrees using filtered datasets, the initial supertree and divide-and-conquer trees, the new fasttrees using new fasttree method (filtered dataset and full dataset), and the trees using hybrid supertree and divide-and-conquer methods (see complete results in Supplementary Table S6).

We compared the performance of filtered datasets to evaluate the effects of different filtering strategies. At the genus level, datasets filtered with indv0 and loci50 (keeping all taxa within specific loci and retaining loci sampled in ≥50% of taxa) achieved the best expected clade recovery. For high-level clades, datasets filtered with sites50 (removing alignment columns where ≥50% of taxa were gaps or missing) performed best. In contrast, more aggressive filtering approaches, such as loci90 (retaining loci sampled in ≥90% of taxa) and indv75 (keeping taxa with ≥75% of sequence completeness), consistently resulted in poorer clade recovery. As expected, filtering reduced the number of sites and CPU time was positively correlated with the size of the supermatrix across all fasttree analyses (R^2^ = 0.8; Supplementary Figure S8). While we observed no consistent pattern in clade recovery between trees estimated with FreeRates and GAMMA models, GAMMA models generally required less computation time.

### Hybrid supertrees and hybrid divide-and-conquer trees

Using a fasttree backbone in the hybrid supertree approach led to poor clade recovery, with some iterations performing worse than our initial analyses using the Jetz+Burleigh backbones (Fig. 4 and Supplementary Table S9). However, as in the initial analyses, adding a taxonomic backbone greatly improved performance, with several hybrid supertree analyses recovering more expected clades than the RAxML-NG tree. Despite these improvements, a better backbone did not eliminate the novel relationships introduced in the supertree analyses. Hybrid supertrees still produced topologies that were the most divergent from those inferred by RAxML-NG, our best new fasttrees, or our best hybrid divide-and-conquer trees (Fig. 4).

The hybrid divide-and-conquer trees were similar to the RAxML-NG tree and the new fasttree based on the full dataset in tree space (Fig. 4). However, even when using a fasttree with strong expected taxa recovery (e.g., the fasttree fulldata parsD), these trees recovered fewer expected clades than the RAxML-NG analysis. While the inclusion of a taxonomic backbone provided some improvement, none of the hybrid divide-and-conquer trees outperformed the best hybrid supertrees (Fig. 4). Additionally, some polytomies observed in the initial analyses persisted, even with the inclusion of both the fasttree and a taxonomic backbone.

### Divergence time estimation

To produce our final time-calibrated macrophylogenies, we incorporated 43 fossil calibrations (Supplementary Table S7; Supplementary information). Despite being estimated using different methods and datasets, the divergence time estimates for key nodes were generally similar across our own seven trees (Fig. 7). Recent studies also show broadly similar relative divergence times (to Neognathae) for comparable groups.

## Discussion

### Baseline phylogeny and expected clade recovery

The RAxML-NG tree provided an excellent estimate of the bird phylogeny, and most cases of non-monophyly at lower taxonomic levels matched results from recently published phylogenomic studies (e.g., Harvey et al. 2020; Smith et al. 2023). Some instances of non-monophyly likely reflected artifacts, such as limited taxon sampling or insufficient sequence data, particularly from historical museum specimens, while others appear to reflect the true phylogenetic relationships of genera or families for which formal taxonomic revision is pending (e.g., *Tyranneutes* nested in *Neopelma* (Leite et al. 2021), *Antilophia* in *Chiroxiphia* (Zhao et al. 2023), and Tityridae divided into Tityridae *sensu stricto*, Onychorhynchidae, and Oxyruncidae (Oliveros et al. 2019)). However, the high computational demand, though expected as the challenges of large tree searches under the likelihood criterion have long been recognized (reviewed by Yang and Rannala 2012), restricted our ability to fully assess uncertainty in the data matrix using bootstrap replicates. To address this limitation, we evaluated alternative analytical approaches that might offer similar or even greater accuracy while requiring fewer computational resources.

### Fasttree approaches

Our initial exploration of computationally efficient methods highlighted that supertree and divide-and-conquer approaches provided the best expected clade recovery, though both required taxonomic backbones to achieve this favorable performance (Fig. 3). Fasttrees, while not as accurate as the RAxML-NG tree or the best-performing supertrees and divide-and-conquer trees, still demonstrated relatively good recovery of expected clades. A notable finding from the initial fasttree analyses was that the fasttree based on the full dataset exhibited poorer clade recovery than most of the filtered datasets. This result suggests that the heterogeneity of the full dataset may interfere with fasttree searches, unlike the RAxML-NG analysis, which appeared more robust to this heterogeneity.

We improved the fasttree search and optimization process by using a single MP starting tree and conducting replicate searches. This modification resulted in a best full data fasttree that achieved phylogenetic accuracy comparable to the RAxML-NG tree, yet with a substantially reduced computational burden (Fig. 4). Although two trees differed in the arrangement of Otidimorphae, Columbimorphae, and Opisthocomiformes (Supplementary Figure S9), the relationships among high-level clades at the base of Neoaves remain a particularly challenging phylogenetic problem (reviewed by Braun et al. 2019), with no consensus achieved to date (cf. Stiller et al. 2024; Wu et al. 2024a).

In contrast to our attempts to improve search efficiency, dataset filtering approaches yielded mixed results. Unlike our initial analyses, filtering to remove missing data did not enhance new fasttree performance in recovering expected clades, likely because filtered datasets also had fewer parsimony informative sites (Fig. 6). We found that site filtering had a greater impact on high-level clade recovery, whereas locus and individual filtering more strongly influenced resolution of expected genera. The most effective filtering strategy likely depends on the taxonomic level of interest. Testing various filtering strategies and models is now highly feasible due to the significantly reduced computation time of the modified fasttree approach. This efficiency also makes it possible to incorporate multiple replicates to account for stochasticity in tree searches. As phylogenomic datasets continue to grow in size, further advancements in computational efficiency for tree estimation will be essential to maintaining the utility of these methods.

**Figure 6.**
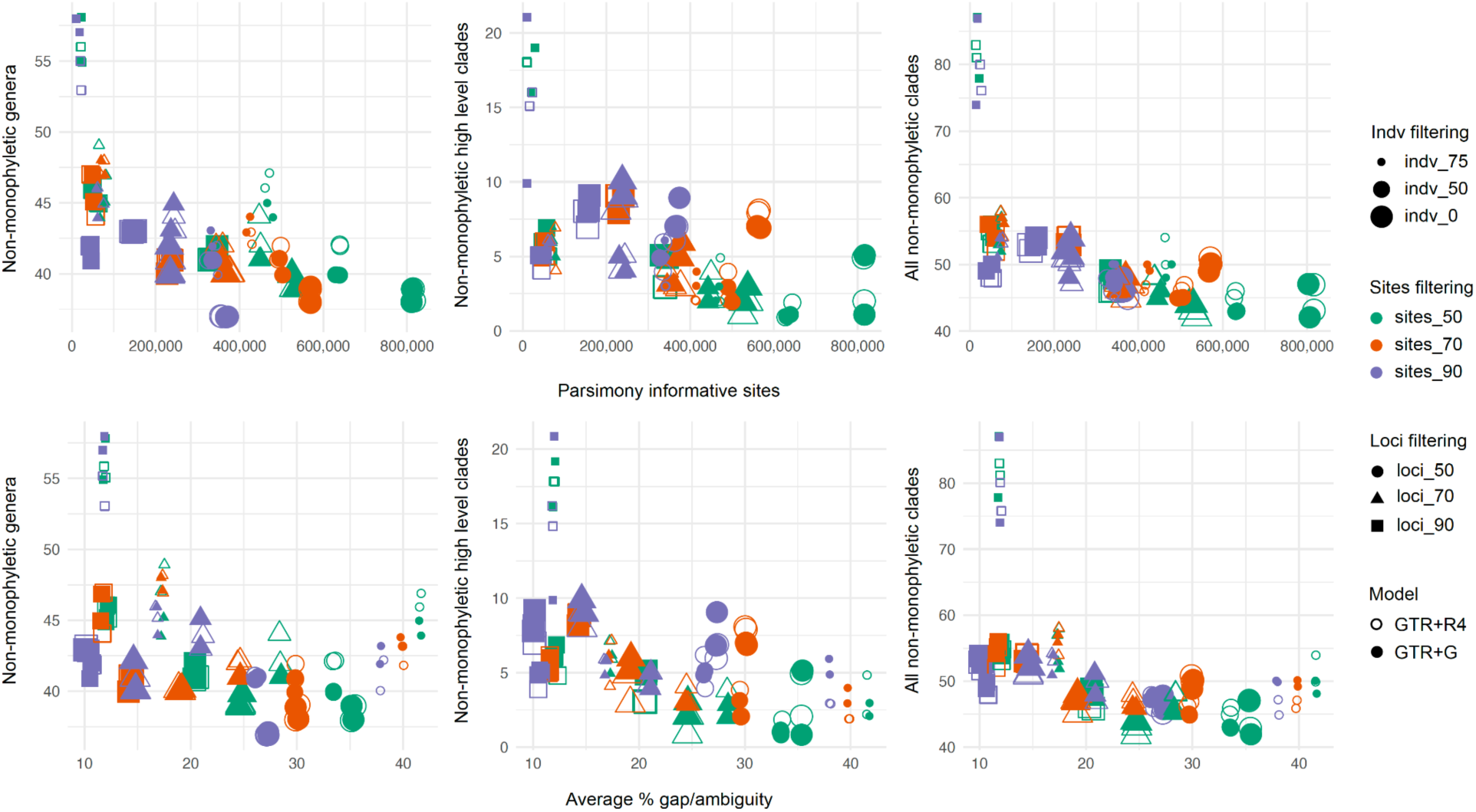
A comparison between the number of unresolved expected clades (genera, high-level clades, and all expected clades combined) and parsimony informative sites (above), as well as average proportion of gaps and ambiguities (“-”, “?” or “N”) across all locus alignments for the dataset (below). For each filtered dataset, four fasttrees with different parsimony starting trees were evaluated. A slight jitter was used when two shapes were completely overlapping each other so that both shapes would be visible.

### Species tree method on heterogeneous datasets

Results from the ASTRAL species tree methods are consistent with previous studies, which have shown that poor sequence recovery and missing data can bias gene tree summary methods (Liu et al. 2010; Springer and Gatesy 2014; Hosner et al. 2016; Xi et al. 2016; Zhao et al. 2025). One contributing factor is the distribution of informative sites in UCE alignments, which are disproportionately located near the ends of the alignments (Faircloth et al. 2012). These regions may be underrepresented when sequence recovery is poor, particularly in lower-quality samples such as those derived from historical museum specimens. Consequently, taxa with poor sequence recovery may be misplaced in estimated gene trees or excluded from certain gene trees altogether, leading to inaccuracies in the ASTRAL tree. Improving ASTRAL trees would entail excluding lower-quality samples and result in a tree with many fewer tips. Overall, ASTRAL was not an accurate method for estimating macrophylogeny with this type of heterogeneous UCE data, even when using the more homogenous filtered subsets. Additionally, it was less computationally efficient than many of the other methods tested (Supplementary Table S8).

### Supertree and divide-and-conquer approaches

Despite apparent computational efficiency, the supertrees contained novel nodes that contradicted all input trees, potentially due to issues of hidden support (e.g., Gatesy et al. 2004; Wilkinson et al. 2005). While signals from the input phylogenomic trees should dominate the supertree topology due to their higher weights relative to the backbones, novel relationships likely arose from topological incompatibilities or asymmetric taxon sampling in the published phylogenomic trees used as input. These issues appeared to be intrinsic to the structures of the input trees (see examples in Supplementary Information). Consequently, hybrid supertrees still produced topologies that were the most divergent from other trees (Fig. 4). This outcome may be explained by the reliance of supertree methods on input trees generated using different analytical approaches by different investigators, as we combined trees from 46 distinct phylogenomic studies. Although the computation time required for supertree analyses was minimal (Fig. 3), this does not include the time needed to locate and code the source trees for analysis.

Overall, while we were able to produce supertrees that provided reasonably accurate representations of the Avian Tree of Life, the methods were not straightforward. Consistent with previous studies, we found that incorporating backbones was critical for improving taxonomic overlap (Redelings and Holder 2017; Kimball et al. 2019; McTavish et al. 2024). An alternative or complementary approach involves pruning problematic taxa from the source trees (Bininda-Emonds et al. 2002) or upweighting more accurate source trees (Bininda-Emonds and Sanderson 2001). While these strategies can improve phylogenetic accuracy, they require prior knowledge and subjective decisions about phylogenetic relationships, which may not always be feasible or unbiased.

Compared to typical supertree approaches, the divide-and-conquer method has advantages, as the individual trees integrated using supertree techniques are generated from sequence data under consistent programs, parameter settings, and computing platforms. While this approach has potential, its performance is dependent on the inclusion of a taxonomic backbone, and further improvements might be achievable with analyses of additional data subsets. All divide-and-conquer trees included unresolved nodes which were particularly evident in species-rich clades where limited overlap in taxon sampling across subsets may have contributed to the increased number of polytomies. This suggests that the 50 subsets used for the divide-and-conquer analyses were insufficient and that additional subsets may be required to improve resolution.

Although the source trees differed between the supertree and divide-and-conquer analyses, both used the same approach to estimate the final tree and faced similar limitations. In both cases, the best results were achieved using taxonomic backbones. While standardized taxonomic backbones are available for well-studied groups like birds, their absence in many other taxonomic groups limits the broader applicability of these methods. Another challenge lies in determining an appropriate weighting scheme for matrix representation with parsimony (MRP; Baum 1992; Ragan 1992). Identifying optimal weights for backbone and source trees, especially when different types of source trees are involved, is difficult. However, the computational efficiency of supertree analyses allows for testing alternative weighting schemes (e.g., Moore et al. 2006; Baker et al. 2009; Nyakatura and Bininda-Emonds 2012) to evaluate their impact on resolution–provided robust criteria, such as expected clade recovery, are available for comparison. Finally, neither method inherently supports branch length estimation. Various approaches can assign branch lengths to supertrees, with or without molecular data (e.g., Purvis 1995; Bininda-Emonds et al. 1999; Torices 2010; Kimball et al. 2019). In our study, branch length-optimized supertrees and divide-and-conquer trees yielded divergence time estimates that were highly similar to those from the concatenated trees, suggesting this limitation may not be critical for most studies.

While traditional supertree and divide-and-conquer methods face similar challenges, they differ in how they address data limitations and computational demands. In traditional supertree approaches, source trees are often limited in number and overlap, necessitating the inclusion of backbones to effectively link taxa. Additionally, genomic data for taxa not included in published phylogenies may need to be incorporated, presenting another challenge. By contrast, the divide-and-conquer method allows for the creation of new subsets that can improve linkage and enhance phylogenetic estimation, albeit at the cost of increased compute time. Moreover, this approach establishes a direct link between sequence data and supertree estimation, addressing the data-dissociation problem inherent in traditional supertree methods (e.g., Moore et al. 2006). Given these advantages, the divide-and-conquer may offer a reasonable alternative for large tree inference, particularly when sufficient computational resources are available.

### Divergence time estimation

The timing of avian phylogeny has been a topic of substantial debate. Some studies support an upper Cretaceous ancient origin for most high-level clades in Neoaves (Pacheco et al. 2011; Mitchell et al. 2015; Wu et al. 2024a; Wu et al. 2024b), while others suggest these lineages originated much closer to the Cretaceous-Paleogene (K-Pg) mass extinction event (∼66 Ma) (Jarvis et al. 2014; Claramunt and Cracraft 2015; Prum et al. 2015; Kimball et al. 2019; Brocklehurst and Field 2024; Claramunt et al. 2024; Stiller et al. 2024). Despite these differences, all studies agree that crown birds originated in the mid- to late-Cretaceous, consistent with crown bird fossils predating the K-Pg boundary (e.g., Field et al. 2020).

Despite variation in tree topologies and branch lengths due to differences in data completeness, divergence time estimates were largely consistent across our methods (Fig. 7). This consistency held regardless of whether branch lengths were estimated during the tree search (RAxML-NG and fasttrees) or added later for methods that do not estimate meaningful branch lengths (supertree and divide-and-conquer analyses). These findings suggest that for downstream comparative analyses requiring time-calibrated trees, the choice of tree estimation method may have minimal impact, provided the method reliably recovers topological relationships. These results, supported by our calibrations, corroborate the hypothesis that the rapid diversification of modern birds occurred near the K-Pg event. Taking these factors into account, we present the first “macrophylogenomic tree” for birds, a resource that can be leveraged in future comparative research.

**Figure 7.**
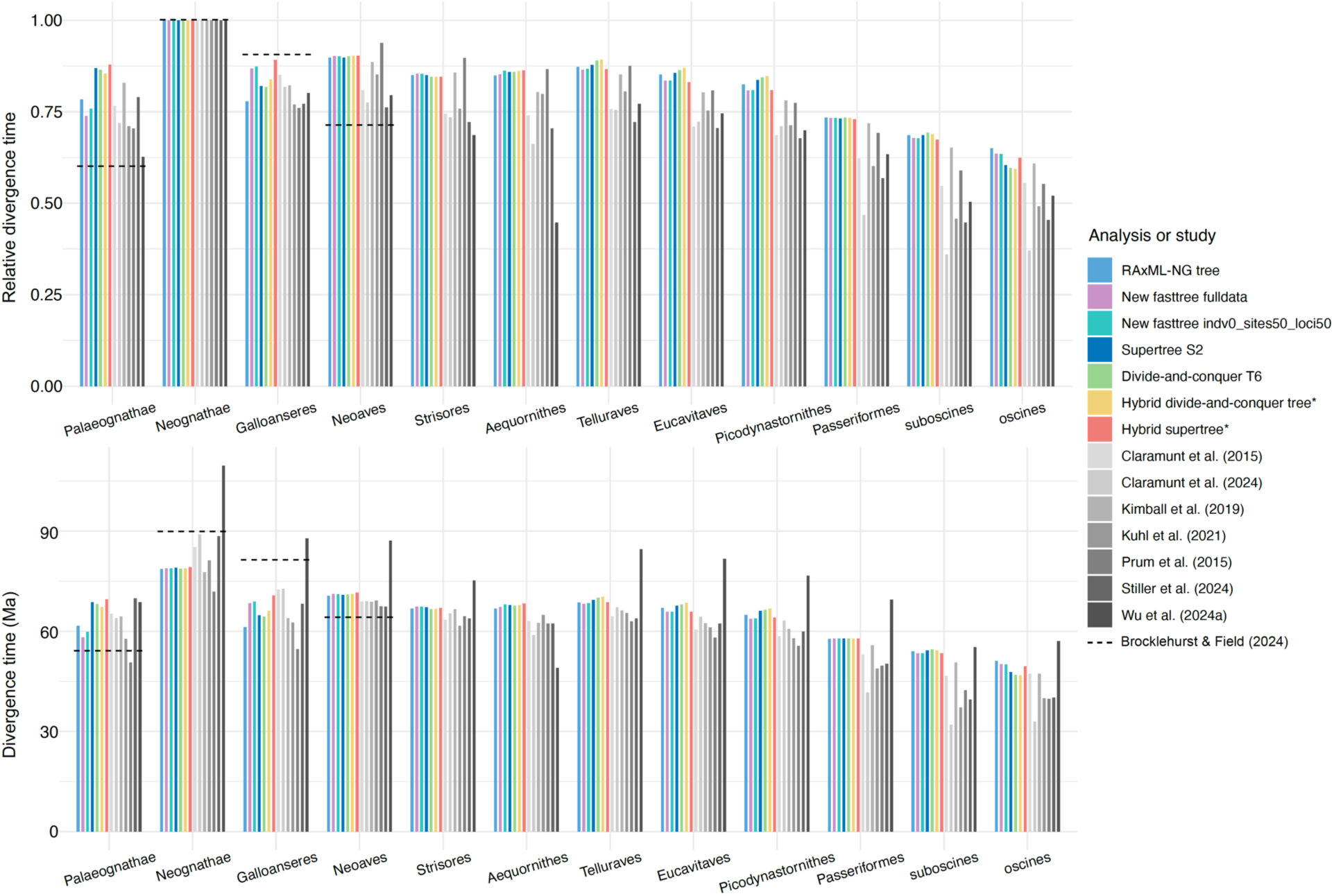
Crown ages for 12 major avian clades with relative divergence time (to Neognathae) on the top panel and absolute divergence time on the bottom panel. Divergence times were extracted from the time-calibrated RAxML-NG tree and the six other time-calibrated trees from various approaches in our study as well as time-calibrated trees from previous studies. Branch lengths of the supertrees and divide-and-conquer trees were optimized using the same filtered dataset (see methods).

## Conclusions

Overall, our analyses demonstrate that accurate macrophylogenies can be estimated using computationally efficient methods. This was achieved with a heterogeneous dataset assembled from many independent studies, reflecting the likely approach for estimating most large-scale phylogenies across the Tree of Life. While assembling such datasets introduces heterogeneity, our results demonstrate that filtering may not always be necessary. In fact, filtering can lead to lower accuracy, as observed in our study, where fewer expected clades were recovered from filtered datasets compared to the full dataset.

Although we successfully estimated trees using several approaches that appeared accurate based on expected clade criterion, both traditional supertree and divide-and-conquer methods required additional information, such as taxonomic backbones, to achieve results comparable to our best ML estimates. By contrast, our new fasttree approach using the full dataset provided a strong alternative to RAxML-NG, delivering similar topological accuracy and branch length estimates with a substantially reduced computational burden. Running multiple replicate analyses to test different MP starting trees and models is also computationally efficient, and simple criteria, such as likelihood values, can be used to assess the resulting trees for those taxonomic groups that lack sufficient study to define expected clades. Thus, the fasttree approach can be broadly applicable to any taxonomic group. By demonstrating the feasibility of computationally efficient methods, this study offers a roadmap for constructing large-scale phylogenies across the Tree of Life.

## Acknowledgements

We thank Bui Quang Minh for insights on the fasttree approach using IQ-TREE and Siavash Mirarab and Chao Zhang for advice on running large ASTRAL trees. We thank Jeremy Kirchman, Zongji Wang, and Qi Zhou for sharing tree files. We also thank J. Klicka, M. Ahmad, W. Tsai Nakashima, and E.M. Smith for support. Portions of this research were conducted with high-performance computing resources provided by Louisiana State University (http://www.hpc.lsu.edu). This work was supported by National Science Foundation grant DEB-1655624 (BCF and RTB), DEB-2217442 (BCF and RTB - supplement), DEB-1655736 (BTS), DEB-1655683 (ELB and RTK) and Villum Fonden 25925 (PAH). Part of this work was funded by UKRI grant MR/X015130/1. For the purpose of open access, the authors have applied a Creative Commons Attribution (CC BY) license to any Author Accepted Manuscript version arising.

## Disclaimer

Any use of trade, firm, or product names is for descriptive purposes only and does not imply endorsement by the US government.

## Notes

### Competing Interest Statement

The authors have declared no competing interest.

## Reference

Andermann T, Fernandes AM, Olsson U, Töpel M, Pfeil B, Oxelman B, Aleixo A, Faircloth BC, Antonelli A. 2019. Allele phasing greatly improves the phylogenetic utility of ultraconserved elements. Syst. Biol. 68:32–46.

Andersen MJ, McCullough JM, Friedman NR, Peterson AT, Moyle RG, Joseph L, Nyári ÁS. 2019. Ultraconserved elements resolve genus-level relationships in a major Australasian bird radiation (Aves: Meliphagidae). Emu - Austral Ornithology 119:218– 232.

Andersen MJ, McCullough JM, Mauck WM, Smith BT, Moyle RG. 2018. A phylogeny of kingfishers reveals an Indomalayan origin and elevated rates of diversification on oceanic islands. J. Biogeogr. 45:269–281.

Bader DA, Roshan U, Stamatakis A. 2006. Computational grand challenges in assembling the tree of life: problems and solutions. In: Computational biology and bioinformatics. Vol. 68. Advances in Computers. Elsevier. p. 127–176.

Baker WJ, Savolainen V, Asmussen-Lange CB, Chase MW, Dransfield J, Forest F, Harley MM, Uhl NW, Wilkinson M. 2009. Complete generic-level phylogenetic analyses of palms (Arecaceae) with comparisons of supertree and supermatrix approaches. Syst. Biol. 58:240–256.

Baum BR. 1992. Combining trees as a way of combining data sets for phylogenetic inference, and the desirability of combining gene trees. Taxon 41:3–10.

Bininda-Emonds OR, Gittleman JL, Purvis A. 1999. Building large trees by combining phylogenetic information: a complete phylogeny of the extant Carnivora (Mammalia). Biol. Rev. Camb. Philos. Soc. 74:143–175.

Bininda-Emonds OR, Sanderson MJ. 2001. Assessment of the accuracy of matrix representation with parsimony analysis supertree construction. Syst. Biol. 50:565–579.

Bininda-Emonds ORP, Gittleman JL, Steel MA. 2002. THE (SUPER)TREE OF LIFE: Procedures, Problems, and Prospects. Annu. Rev. Ecol. Syst. 33:265–289.

Bininda-Emonds ORP. 2004. The evolution of supertrees. Trends Ecol. Evol. 19:315–322.

Borowiec ML. 2016. AMAS: a fast tool for alignment manipulation and computing of summary statistics. PeerJ 4:e1660.

Braun EL, Cracraft J, Houde P. 2019. Resolving the Avian Tree of Life from Top to Bottom: The Promise and Potential Boundaries of the Phylogenomic Era. In: Kraus RHS, editor. Avian Genomics in Ecology and Evolution: From the Lab into the Wild. Cham: Springer International Publishing. p. 151–210.

Braun EL, Oliveros CH, White Carreiro ND, Zhao M, Glenn TC, Brumfield RT, Braun MJ, Kimball RT, Faircloth BC. 2024. Testing the mettle of METAL: A comparison of phylogenomic methods using a challenging but well-resolved phylogeny. BioRxiv.

Brocklehurst N, Field DJ. 2024. Tip dating and Bayes factors provide insight into the divergences of crown bird clades across the end-Cretaceous mass extinction. Proc. Biol. Sci. 291:20232618.

Bruxaux J, Gabrielli M, Ashari H, Prŷs-Jones R, Joseph L, Milá B, Besnard G, Thébaud C. 2018. Recovering the evolutionary history of crowned pigeons (Columbidae: Goura): Implications for the biogeography and conservation of New Guinean lowland birds. Mol. Phylogenet. Evol. 120:248–258.

Bryson RW, Faircloth BC, Tsai WLE, McCormack JE, Klicka J. 2016. Target enrichment of thousands of ultraconserved elements sheds new light on early relationships within New World sparrows (Aves: Passerellidae). Auk 133:451–458.

Burga A, Wang W, Ben-David E, Wolf PC, Ramey AM, Verdugo C, Lyons K, Parker PG, Kruglyak L. 2017. A genetic signature of the evolution of loss of flight in the Galapagos cormorant. Science 356.

Burleigh JG, Kimball RT, Braun EL. 2015. Building the avian tree of life using a large-scale, sparse supermatrix. Mol. Phylogenet. Evol. 84:53–63.

Campillo LC, Oliveros CH, Sheldon FH, Moyle RG. 2018. Genomic data resolve gene tree discordance in spiderhunters (Nectariniidae, Arachnothera). Mol. Phylogenet. Evol. 120:151–157.

Capella-Gutiérrez S, Silla-Martínez JM, Gabaldón T. 2009. trimAl: A tool for automated alignment trimming in large-scale phylogenetic analyses. Bioinformatics 25:1972– 1973.

Catanach TA, Halley MR, Allen JM, Johnson JA, Thorstrom R, Palhano S, Poor Thunder C, Gallardo JC, Weckstein JD. 2021. Systematics and conservation of an endemic radiation of Accipiter hawks in the Caribbean islands. Ornithology 138.

Charif D, Lobry JR. 2007. SeqinR 1.0-2: A Contributed Package to the R Project for Statistical Computing Devoted to Biological Sequences Retrieval and Analysis. In: Bastolla U, Porto M, Roman HE, Vendruscolo M, editors. Structural approaches to sequence evolution. Biological and medical physics, biomedical engineering. Berlin, Heidelberg: Springer Berlin Heidelberg. p. 207–232.

Chen A, Field DJ. 2020. Phylogenetic definitions for Caprimulgimorphae (Aves) and major constituent clades under the International Code of Phylogenetic Nomenclature. Senckenberg Gesellschaft für Naturforschung.

Chen D, Braun EL, Forthman M, Kimball RT, Zhang Z. 2018. A simple strategy for recovering ultraconserved elements, exons, and introns from low coverage shotgun sequencing of museum specimens: Placement of the partridge genus Tropicoperdix within the galliformes. Mol. Phylogenet. Evol. 129:304–314.

Claramunt S, Braun EL, Cracraft J, Fjeldså J, Ho SYW, Houde P, Nguyen JMT, Stiller J. 2024. Calibrating the genomic clock of modern birds using fossils. Proc Natl Acad Sci USA 121:e2405887121.

Claramunt S, Cracraft J. 2015. A new time tree reveals Earth history’s imprint on the evolution of modern birds. Sci. Adv. 1:e1501005.

Clements JF, Rasmussen PC, Schulenberg TS, Iliff MJ, Fredericks TA, Gerbracht JA, Lepage D, Spencer A, Billerman SM, Sullivan BL, et al. 2023. The eBird/Clements checklist of Birds of the World: v2023. Available from: https://www.birds.cornell.edu/clementschecklist/download/

Cotton JA, Wilkinson M. 2009. Supertrees join the mainstream of phylogenetics. Trends Ecol. Evol. 24:1–3.

Cracraft J, Donoghue MJ. 2004. Assembling the tree of life. Oxford University Press

Creevey CJ, McInerney JO. 2005. Clann: investigating phylogenetic information through supertree analyses. Bioinformatics 21:390–392.

Danecek P, Auton A, Abecasis G, Albers CA, Banks E, DePristo MA, Handsaker RE, Lunter G, Marth GT, Sherry ST, et al. 2011. The variant call format and VCFtools. Bioinformatics 27:2156–2158.

Delsuc F, Brinkmann H, Philippe H. 2005. Phylogenomics and the reconstruction of the tree of life. Nat. Rev. Genet. 6:361–375.

Dickinson EC, Christidis L eds. 2014. The Howard and Moore Complete Checklist of the Birds of the World Fourth Edition. Eastbourne: Aves Press

Driskell AC, Ané C, Burleigh JG, McMahon MM, O’meara BC, Sanderson MJ. 2004. Prospects for building the tree of life from large sequence databases. Science 306:1172–1174.

Dwivedi B, Gadagkar SR. 2009. Phylogenetic inference under varying proportions of indel- induced alignment gaps. BMC Evol. Biol. 9:211.

Everson KM, McLaughlin JF, Cato IA, Evans MM, Gastaldi AR, Mills KK, Shink KG, Wilbur SM, Winker K. 2019. Speciation, gene flow, and seasonal migration in Catharus thrushes (Aves:Turdidae). Mol. Phylogenet. Evol. 139:106564.

Faircloth BC, McCormack JE, Crawford NG, Harvey MG, Brumfield RT, Glenn TC. 2012. Ultraconserved elements anchor thousands of genetic markers spanning multiple evolutionary timescales. Syst. Biol. 61:717–726.

Faircloth BC. 2016. PHYLUCE is a software package for the analysis of conserved genomic loci. Bioinformatics 32:786–788.

Felsenstein J. 1985. Confidence limits on phylogenies: an approach using the bootstrap. Evolution 39:783–791.

Ferreira M, Fernandes AM, Aleixo A, Antonelli A, Olsson U, Bates JM, Cracraft J, Ribas CC. 2018. Evidence for mtDNA capture in the jacamar Galbula leucogastra/chalcothorax species-complex and insights on the evolution of white-sand ecosystems in the Amazon basin. Mol. Phylogenet. Evol. 129:149–157.

Field DJ, Benito J, Chen A, Jagt JWM, Ksepka DT. 2020. Late Cretaceous neornithine from Europe illuminates the origins of crown birds. Nature 579:397–401.

Gascuel O. 1997. BIONJ: an improved version of the NJ algorithm based on a simple model of sequence data. Mol. Biol. Evol. 14:685–695.

Gatesy J, Baker RH, Hayashi C. 2004. Inconsistencies in arguments for the supertree approach: supermatrices versus supertrees of Crocodylia. Syst. Biol. 53:342–355.

Gill F, Donsker D, Rasmussen P. 2023. IOC World Bird List (v 13.1). Available from: 10.14344/IOC.ML.13.1

Goloboff PA, Catalano SA, Marcos Mirande J, Szumik CA, Salvador Arias J, Källersjö M, Farris JS. 2009. Phylogenetic analysis of 73 060 taxa corroborates major eukaryotic groups. Cladistics 25:211–230.

Grealey J, Lannelongue L, Saw W-Y, Marten J, Méric G, Ruiz-Carmona S, Inouye M. 2022. The carbon footprint of bioinformatics. Mol. Biol. Evol. 39.

Gu Z. 2022. Complex heatmap visualization. iMeta 1:e43.

Hackett SJ, Kimball RT, Reddy S, Bowie RCK, Braun EL, Braun MJ, Chojnowski JL, Cox WA, Han K-L, Harshman J, et al. 2008. A phylogenomic study of birds reveals their evolutionary history. Science 320:1763–1768.

Harvey MG, Bravo GA, Claramunt S, Cuervo AM, Derryberry GE, Battilana J, Seeholzer GF, McKay JS, O’Meara BC, Faircloth BC, et al. 2020. The evolution of a tropical biodiversity hotspot. Science 370:1343–1348.

Hinchliff CE, Smith SA, Allman JF, Burleigh JG, Chaudhary R, Coghill LM, Crandall KA, Deng J, Drew BT, Gazis R, et al. 2015. Synthesis of phylogeny and taxonomy into a comprehensive tree of life. Proc Natl Acad Sci USA 112:12764–12769.

Hosner PA, Faircloth BC, Glenn TC, Braun EL, Kimball RT. 2016. Avoiding missing data biases in phylogenomic inference: an empirical study in the landfowl (aves: galliformes). Mol. Biol. Evol. 33:1110–1125.

Huerta-Cepas J, Serra F, Bork P. 2016. ETE 3: reconstruction, analysis, and visualization of phylogenomic data. Mol. Biol. Evol. 33:1635–1638.

Imfeld TS, Barker FK, Brumfield RT. 2020. Mitochondrial genomes and thousands of ultraconserved elements resolve the taxonomy and historical biogeography of the Euphonia and Chlorophonia finches (Passeriformes: Fringillidae). Auk 137.

Jarvis ED, Mirarab S, Aberer AJ, Li B, Houde P, Li C, Ho SYW, Faircloth BC, Nabholz B, Howard JT, et al. 2014. Whole-genome analyses resolve early branches in the tree of life of modern birds. Science 346:1320–1331.

Jetz W, Thomas GH, Joy JB, Hartmann K, Mooers AO. 2012. The global diversity of birds in space and time. Nature 491:444–448.

Kapli P, Yang Z, Telford MJ. 2020. Phylogenetic tree building in the genomic age. Nat. Rev. Genet. 21:428–444.

Katoh K, Standley DM. 2013. MAFFT multiple sequence alignment software version 7: improvements in performance and usability. Mol. Biol. Evol. 30:772–780.

Kimball RT, Braun EL, Barker FK, Bowie RCK, Braun MJ, Chojnowski JL, Hackett SJ, Han K-L, Harshman J, Heimer-Torres V, et al. 2009. A well-tested set of primers to amplify regions spread across the avian genome. Mol. Phylogenet. Evol. 50:654–660.

Kimball RT, Oliveros CH, Wang N, White ND, Barker FK, Field DJ, Ksepka DT, Chesser RT, Moyle RG, Braun MJ, et al. 2019. A phylogenomic supertree of birds. Diversity (Basel) 11:109.

Kirchman JJ, Rotzel McInerney N, Giarla TC, Olson SL, Slikas E, Fleischer RC. 2021. Phylogeny based on ultra-conserved elements clarifies the evolution of rails and allies (Ralloidea) and is the basis for a revised classification. Ornithology.

Koonin EV. 2005. Orthologs, paralogs, and evolutionary genomics. Annu. Rev. Genet. 39:309–338.

Kozlov AM, Darriba D, Flouri T, Morel B, Stamatakis A. 2019. RAxML-NG: a fast, scalable and user-friendly tool for maximum likelihood phylogenetic inference. Bioinformatics 35:4453–4455.

Kuhl H, Frankl-Vilches C, Bakker A, Mayr G, Nikolaus G, Boerno ST, Klages S, Timmermann B, Gahr M. 2021. An Unbiased Molecular Approach Using 3’-UTRs Resolves the Avian Family-Level Tree of Life. Mol. Biol. Evol. 38:108–127.

Kumar S. 2022. Embracing green computing in molecular phylogenetics. Mol. Biol. Evol. 39.

Lamichhaney S, Berglund J, Almén MS, Maqbool K, Grabherr M, Martinez-Barrio A, Promerová M, Rubin C-J, Wang C, Zamani N, et al. 2015. Evolution of Darwin’s finches and their beaks revealed by genome sequencing. Nature 518:371–375.

Lê S, Josse J, Husson F. 2008. FactoMineR : An R package for multivariate analysis. J. Stat. Softw. 25.

Leite RN, Kimball RT, Braun EL, Derryberry EP, Hosner PA, Derryberry GE, Anciães M, McKay JS, Aleixo A, Ribas CC, et al. 2021. Phylogenomics of manakins (Aves: Pipridae) using alternative locus filtering strategies based on informativeness. Mol. Phylogenet. Evol. 155:107013.

Liu FG, Miyamoto MM, Freire NP, Ong PQ, Tennant MR, Young TS, Gugel KF. 2001. Molecular and morphological supertrees for eutherian (placental) mammals. Science 291:1786–1789.

Liu L, Yu L, Edwards SV. 2010. A maximum pseudo-likelihood approach for estimating species trees under the coalescent model. BMC Evol. Biol. 10:302.

Manthey JD, Campillo LC, Burns KJ, Moyle RG. 2016. Comparison of Target-Capture and Restriction-Site Associated DNA Sequencing for Phylogenomics: A Test in Cardinalid Tanagers (Aves, Genus: Piranga). Syst. Biol. 65:640–650.

McCormack JE, Harvey MG, Faircloth BC, Crawford NG, Glenn TC, Brumfield RT. 2013. A phylogeny of birds based on over 1,500 loci collected by target enrichment and high- throughput sequencing. PLoS ONE 8:e54848.

McCormack JE, Hird SM, Zellmer AJ, Carstens BC, Brumfield RT. 2013. Applications of next- generation sequencing to phylogeography and phylogenetics. Mol. Phylogenet. Evol. 66:526–538.

McCormack JE, Tsai WLE, Faircloth BC. 2016. Sequence capture of ultraconserved elements from bird museum specimens. Mol. Ecol. Resour 16:1189–1203.

McCullough JM., Joseph L, Moyle RG, Andersen MJ. 2019. Ultraconserved elements put the final nail in the coffin of traditional use of the genus Meliphaga (Aves: Meliphagidae). Zool. Scr. 48:411–418.

McCullough JM, Moyle RG, Smith BT, Andersen MJ. 2019. A Laurasian origin for a pantropical bird radiation is supported by genomic and fossil data (Aves: Coraciiformes). Proc. Biol. Sci. 286:20190122.

McCullough JM, Oliveros CH, Benz BW, Zenil-Ferguson R, Cracraft J, Moyle RG, Andersen MJ. 2022. Wallacean and Melanesian Islands Promote Higher Rates of Diversification within the Global Passerine Radiation Corvides. Syst. Biol. 71:1423–1439.

McTavish EJ, Gerbracht JA, Holder MT, Iliff MJ, Lepage D, Rasmussen P, Redelings B, Sanchez Reyes LL, Miller ET. 2024. A complete and dynamic tree of birds. BioRxiv.

Minh BQ, Schmidt HA, Chernomor O, Schrempf D, Woodhams MD, von Haeseler A, Lanfear R. 2020. IQ-TREE 2: New models and efficient methods for phylogenetic inference in the genomic era. Mol. Biol. Evol. 37:1530–1534.

Mitchell KJ, Cooper A, Phillips MJ. 2015. Comment on “Whole-genome analyses resolve early branches in the tree of life of modern birds”. Science 349:1460.

Molloy EK, Warnow T. 2018. To include or not to include: the impact of gene filtering on species tree estimation methods. Syst. Biol. 67:285–303.

Moore B, Smith S, Donoghue M. 2006. Increasing Data Transparency and Estimating Phylogenetic Uncertainty in Supertrees: Approaches Using Nonparametric Bootstrapping. Syst. Biol. 55:662–676.

Moyle RG, Oliveros CH, Andersen MJ, Hosner PA, Benz BW, Manthey JD, Travers SL, Brown RM, Faircloth BC. 2016. Tectonic collision and uplift of Wallacea triggered the global songbird radiation. Nat. Commun. 7:12709.

Musher LJ, Cracraft J. 2018. Phylogenomics and species delimitation of a complex radiation of Neotropical suboscine birds (Pachyramphus). Mol. Phylogenet. Evol. 118:204–221.

Nater A, Burri R, Kawakami T, Smeds L, Ellegren H. 2015. Resolving Evolutionary Relationships in Closely Related Species with Whole-Genome Sequencing Data. Syst. Biol. 64:1000–1017.

Nguyen L-T, Schmidt HA, von Haeseler A, Minh BQ. 2015. IQ-TREE: a fast and effective stochastic algorithm for estimating maximum-likelihood phylogenies. Mol. Biol. Evol. 32:268–274.

Nixon K. 1999. The parsimony ratchet, a new method for rapid parsimony analysis. Cladistics 15:407–414.

Nyakatura K, Bininda-Emonds ORP. 2012. Updating the evolutionary history of Carnivora (Mammalia): a new species-level supertree complete with divergence time estimates. BMC Biol. 10:12.

Ogden TH, Rosenberg MS. 2006. Multiple sequence alignment accuracy and phylogenetic inference. Syst. Biol. 55:314–328.

Oliveros CH, Andersen MJ, Hosner PA, Mauck WM, Sheldon FH, Cracraft J, Moyle RG. 2020. Rapid Laurasian diversification of a pantropical bird family during the Oligocene– Miocene transition. Ibis 162:137–152.

Oliveros CH, Andersen MJ, Moyle RG. 2021. A phylogeny of white-eyes based on ultraconserved elements. Mol. Phylogenet. Evol. 164:107273.

Oliveros CH, Field DJ, Ksepka DT, Barker FK, Aleixo A, Andersen MJ, Alström P, Benz BW, Braun EL, Braun MJ, et al. 2019. Earth history and the passerine superradiation. Proc Natl Acad Sci USA 116:7916–7925.

Ottenburghs J, Megens H-J, Kraus RHS, Madsen O, van Hooft P, van Wieren SE, Crooijmans RPMA, Ydenberg RC, Groenen MAM, Prins HHT. 2016. A tree of geese: A phylogenomic perspective on the evolutionary history of True Geese. Mol. Phylogenet. Evol. 101:303– 313.

Pacheco MA, Battistuzzi FU, Lentino M, Aguilar RF, Kumar S, Escalante AA. 2011. Evolution of modern birds revealed by mitogenomics: timing the radiation and origin of major orders. Mol. Biol. Evol. 28:1927–1942.

Page AJ, Taylor B, Delaney AJ, Soares J, Seemann T, Keane JA, Harris SR. 2016. SNP-sites: rapid efficient extraction of SNPs from multi-FASTA alignments. Microb. Genom. 2:e000056.

Paradis E, Schliep K. 2019. ape 5.0: an environment for modern phylogenetics and evolutionary analyses in R. Bioinformatics 35:526–528.

Parham JF, Donoghue PCJ, Bell CJ, Calway TD, Head JJ, Holroyd PA, Inoue JG, Irmis RB, Joyce WG, Ksepka DT, et al. 2012. Best practices for justifying fossil calibrations. Syst. Biol. 61:346–359.

Philippe H, Brinkmann H, Lavrov DV, Littlewood DTJ, Manuel M, Wörheide G, Baurain D. 2011. Resolving difficult phylogenetic questions: why more sequences are not enough. PLoS Biol. 9:e1000602.

Philippe H, Snell EA, Bapteste E, Lopez P, Holland PWH, Casane D. 2004. Phylogenomics of eukaryotes: impact of missing data on large alignments. Mol. Biol. Evol. 21:1740–1752.

Price MN, Dehal PS, Arkin AP. 2010. FastTree 2 — approximately maximum-likelihood trees for large alignments. PLoS ONE 5:e9490.

Prum RO, Berv JS, Dornburg A, Field DJ, Townsend JP, Lemmon EM, Lemmon AR. 2015. A comprehensive phylogeny of birds (Aves) using targeted next-generation DNA sequencing. Nature 526:569–573.

Purvis A. 1995. A composite estimate of primate phylogeny. Philos. Trans. R. Soc. Lond. B Biol. Sci. 348:405–421.

Queiroz K de, Cantino PD, Gauthier JA. 2020. Phylonyms: A companion to the phylocode. (de Queiroz K, Cantino P, Gauthier J, editors.). Boca Raton : CRC Press, [2019]: CRC Press

Ragan MA. 1992. Phylogenetic inference based on matrix representation of trees. Mol. Phylogenet. Evol. 1:53–58.

Reddy S, Kimball RT, Pandey A, Hosner PA, Braun MJ, Hackett SJ, Han K-L, Harshman J, Huddleston CJ, Kingston S, et al. 2017. Why Do Phylogenomic Data Sets Yield Conflicting Trees? Data Type Influences the Avian Tree of Life more than Taxon Sampling. Syst. Biol. 66:857–879.

Redelings BD, Holder MT. 2017. A supertree pipeline for summarizing phylogenetic and taxonomic information for millions of species. PeerJ 5:e3058.

Revell LJ. 2012. phytools: An R package for phylogenetic comparative biology (and other things). Methods in Ecology and Evolution 3:217–223.

Robinson DF, Foulds LR. 1981. Comparison of phylogenetic trees. Math. Biosci. 53:131–147.

R Core Team. 2023. R: A language and environment for statistical computing. Vienna, Austria: R Foundation for Statistical Computing

Sackton TB, Grayson P, Cloutier A, Hu Z, Liu JS, Wheeler NE, Gardner PP, Clarke JA, Baker AJ, Clamp M, et al. 2019. Convergent regulatory evolution and loss of flight in paleognathous birds. Science 364:74–78.

Salter JF, Oliveros CH, Hosner PA, Manthey JD, Robbins MB, Moyle RG, Brumfield RT, Faircloth BC. 2020. Extensive paraphyly in the typical owl family (Strigidae). Auk 137.

Sanderson MJ, McMahon MM, Steel M. 2010. Phylogenomics with incomplete taxon coverage: the limits to inference. BMC Evol. Biol. 10:155.

Sanderson MJ, Purvis A, Henze C. 1998. Phylogenetic supertrees: Assembling the trees of life. Trends Ecol. Evol. 13:105–109.

Sangster G, Braun EL, Johansson US, Kimball RT, Mayr G, Suh A. 2022. Phylogenetic definitions for 25 higher-level clade names of birds. Avian Res.:100027.

Sangster G, Mayr G. 2021. Feraequornithes: a name for the clade formed by Procellariiformes, Sphenisciformes, Ciconiiformes, Suliformes and Pelecaniformes (Aves). Vertebr. Zool. 71:49–53.

Schliep KP. 2011. phangorn: phylogenetic analysis in R. Bioinformatics 27:592–593.

Schwery O, O’Meara BC. 2016. MonoPhy : a simple R package to find and visualize monophyly issues. PeerJ Computer Science 2:e56.

Smith BT, Harvey MG, Faircloth BC, Glenn TC, Brumfield RT. 2014. Target capture and massively parallel sequencing of ultraconserved elements for comparative studies at shallow evolutionary time scales. Syst. Biol. 63:83–95.

Smith BT, Mauck WM, Benz BW, Andersen MJ. 2020. Uneven Missing Data Skew Phylogenomic Relationships within the Lories and Lorikeets. Genome Biol. Evol. 12:1131–1147.

Smith BT, Merwin J, Provost KL, Thom G, Brumfield RT, Ferreira M, Mauck WM, Moyle RG, Wright TF, Joseph L. 2023. Phylogenomic Analysis of the Parrots of the World Distinguishes Artifactual from Biological Sources of Gene Tree Discordance. Syst. Biol. 72:228–241.

Smith SA, O’Meara BC. 2012. treePL: Divergence time estimation using penalized likelihood for large phylogenies. Bioinformatics 28:2689–2690.

Springer MS, Gatesy J. 2014. Land plant origins and coalescence confusion. Trends Plant Sci. 19:267–269.

Stiller J, Feng S, Chowdhury A-A, Rivas-González I, Duchêne DA, Fang Q, Deng Y, Kozlov A, Stamatakis A, Claramunt S, et al. 2024. Complexity of avian evolution revealed by family-level genomes. Nature 629:851–860.

Sukumaran J, Holder MT. 2010. DendroPy: a Python library for phylogenetic computing. Bioinformatics 26:1569–1571.

Sun K, Meiklejohn KA, Faircloth BC, Glenn TC, Braun EL, Kimball RT. 2014. The evolution of peafowl and other taxa with ocelli (eyespots): a phylogenomic approach. Proc. Biol. Sci. 281.

Swofford DL. 2003. PAUP* Phylogenetic Analysis Using Parsimony (*and Other Methods). Version 4. http://paup.csit.fsu.edu/.

Title PO, Rabosky DL. 2017. Do Macrophylogenies Yield Stable Macroevolutionary Inferences? An Example from Squamate Reptiles. Syst. Biol. 66:843–856.

Torices R. 2010. Adding time-calibrated branch lengths to the Asteraceae supertree. J. Syst. Evol. 48:271–278.

Vianna JA, Fernandes FAN, Frugone MJ, Figueiró HV, Pertierra LR, Noll D, Bi K, Wang-Claypool CY, Lowther A, Parker P, et al. 2020. Genome-wide analyses reveal drivers of penguin diversification. Proc Natl Acad Sci USA 117:22303–22310.

Vinay KL, Natesh M, Mehta P, Jayapal R, Mukherjee S, Robin VV. 2022. Re-assessing the phylogenetic status and evolutionary relationship of Forest Owlet [Athene blewitti (Hume 1873)] using genomic data. Ibis.

Wang N, Hosner PA, Liang B, Braun EL, Kimball RT. 2017. Historical relationships of three enigmatic phasianid genera (Aves: Galliformes) inferred using phylogenomic and mitogenomic data. Mol. Phylogenet. Evol. 109:217–225.

Wang Z, Zhang J, Xu X, Witt C, Deng Y, Chen G, Meng G, Feng S, Xu L, Szekely T, et al. 2022. Phylogeny and sex chromosome evolution of Palaeognathae. J. Genet. Genomics 49:109–119.

White ND, Braun MJ. 2019. Extracting phylogenetic signal from phylogenomic data: Higher-level relationships of the nightbirds (Strisores). Mol. Phylogenet. Evol. 141:106611.

White ND, Mitter C, Braun MJ. 2017. Ultraconserved elements resolve the phylogeny of potoos (Aves: Nyctibiidae). J. Avian Biol. 48:872–880.

Wickham H. 2011. ggplot2. WIREs Comp Stat 3:180–185.

Wilkinson M, Pisani D, Cotton JA, Corfe I. 2005. Measuring support and finding unsupported relationships in supertrees. Syst. Biol. 54:823–831.

Winker K, Glenn TC, Faircloth BC. 2018. Ultraconserved elements (UCEs) illuminate the population genomics of a recent, high-latitude avian speciation event. PeerJ 6:e5735.

Wu S, Rheindt FE, Zhang J, Wang J, Zhang L, Quan C, Li Z, Wang M, Wu F, Qu Y, et al. 2024a. Genomes, fossils, and the concurrent rise of modern birds and flowering plants in the Late Cretaceous. Proc Natl Acad Sci USA 121:e2319696121.

Wu S, Rheindt FE, Zhang J, Wang J, Zhang L, Quan C, Li Z, Wang M, Wu F, Qu Y, et al. 2024b. Reply to Claramunt et al.: Robustness of the Cretaceous radiation of crown aves. Proc Natl Acad Sci USA 121:e2412448121.

Xi Z, Liu L, Davis CC. 2016. The impact of missing data on species tree estimation. Mol. Biol. Evol. 33:838–860.

Yang Z, Rannala B. 2012. Molecular phylogenetics: principles and practice. Nat. Rev. Genet. 13:303–314.

Yonezawa T, Segawa T, Mori H, Campos PF, Hongoh Y, Endo H, Akiyoshi A, Kohno N, Nishida S, Wu J, et al. 2017. Phylogenomics and morphology of extinct paleognaths reveal the origin and evolution of the ratites. Curr. Biol. 27:68–77.

Younger JL, Strozier L, Maddox JD, Nyári ÁS, Bonfitto MT, Raherilalao MJ, Goodman SM, Reddy S. 2018. Hidden diversity of forest birds in Madagascar revealed using integrative taxonomy. Mol. Phylogenet. Evol. 124:16–26.

Zarza E, Faircloth BC, Tsai WLE, Bryson RW, Klicka J, McCormack JE. 2016. Hidden histories of gene flow in highland birds revealed with genomic markers. Mol. Ecol. 25:5144– 5157.

Zhang C, Rabiee M, Sayyari E, Mirarab S. 2018. ASTRAL-III: Polynomial time species tree reconstruction from partially resolved gene trees. BMC Bioinformatics 19:153.

Zhang G, Li B, Li C, Gilbert MTP, Jarvis ED, Wang J, Avian Genome Consortium. 2014. Comparative genomic data of the Avian Phylogenomics Project. Gigascience 3:26.

Zhao M, Kurtis SM, White ND, Moncrieff AE, Leite RN, Brumfield RT, Braun EL, Kimball RT. 2023. Exploring conflicts in whole genome phylogenetics: A case study within manakins (aves: pipridae). Syst. Biol. 72:161–178.

Zhao M., Oswald JA., Allen JM, Owens HL, Hosner PA, Guralnick RP, Braun EL and Kimball RT. 2025. A phylogenomic tree of wood-warblers (Aves: Parulidae): Dealing with good, bad, and ugly samples. Mol. Phylogenet. Evol. 202:108235.

