## Supplementary for "Efficient Inference of Macrophylogenies: Insights from the Avian Tree of Life"

#### Supplementary Figures

**Figure S1**. Schematic representation showing the selection of taxa for partially stratified subsets used in the divide-and-conquer approach. The full data set was stratified into six major groups (Supplementary Data S5). Then, 30, 40, 30, 25, 12 and 13 taxa were randomly selected from each major group to produce a partially stratified subset of 150 taxa.

**Figure S2**. Schematic representation showing the selection of taxa for fully stratified subsets used in the divide-and-conquer approach. Taxa in the full data set (1 to 2600; see Supplementary Data S5) were arranged by taxonomic order and stratified into 25 subsets based on taxonomy to ensure all taxa were represented at least once across the subsets, and were included in trees with congeners. Each vertical blue line represents a taxon present; those that cluster together form an ingroup whereas the more scattered lines are "linker taxa".

**Figure S3**. "Trees-of-trees" for trees generated using IQ-TREE with the -fast option and different starting trees using (A) filter set 1 (indv_50_sites_50_loci_50) data and (B) filter set 3 (indv_0_sites_90_loci_50) data. These trees were generated by neighbor joining of Robinson-Foulds distances among the fasttrees. Apart from the RAxML-NG tree, which is included for comparison, all trees are named based on the starting tree. The pink highlight indicates cases where the starting trees were generated using the same filter set that was used for the fasttree search; clade recovery values listed in the pink box are. We indicate the optimal trees based on their likelihood given the GTR+G and GTR+R4 models (all optimal trees based on likelihood are in the pink box); the FreeRates (+R4) model had a higher likelihood than the GAMMA (+G) model. The numbers in parentheses after each tree are the total numbers of non-monophyletic expected clades. Fasttrees with the best expected clade recovery (i.e., the smallest number of non-monophylies) are underlined. Complete results can be found in Data Availability.

**Figure S4**. Summary statistics of the UCE alignments in all 27 filtered datasets. Each summary statistic (column) was z-transformed. For missing data information (Loci w. <50% missing and Avg. non-gap/ambiguity), the lighter the colors, the more missing data. For each cluster of datasets, we selected one representative (marked with an asterisk) based on the overall performance of the new fasttrees in resolving credible clades.

**Figure S5**. Heatmap based on normalized summary statistics calculated per individual for the full data sets showing the variation across distinct datasets. The colors on the left bar correspond to distinct data sets in the legend.

**Figure S6**. Principal Component Analysis (PCA) showing clustering patterns among samples based on summary statistics calculated per individual for the full and six filtered datasets.

**Figure S7**. A genus-level RAxML-NG tree with major bird groups color-coded. Tip labels in red represent the non-monophyletic genera.

**Figure S8**. CPU hours spent for new fasttree analysis vs. total number of sites in the dataset for the full dataset and 27 filtered datasets.

**Figure S9**. Differences in rearrangements of major superordinal clades within Neoaves across supermatrix, supertree and divide-and-conquer analyses.

### Figure S1

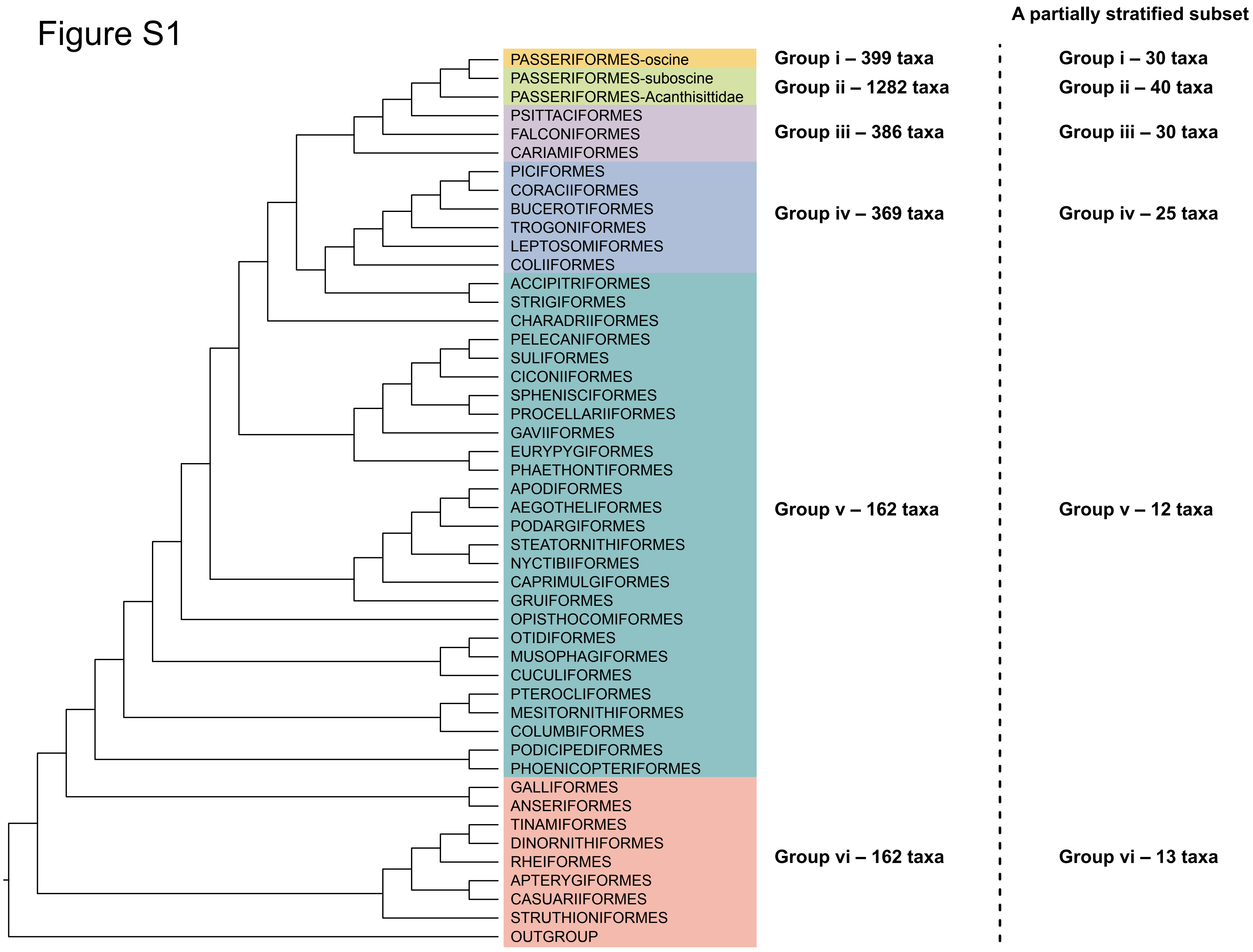

A partially stratified subset
PASSERIFORMES-oscine
PASSERIFORMES-suboscine
PASSERIFORMES-Acanthisittidae
PSITTACIFORMES
FALCONIFORMES
CARIAMIFORMES
PICIFORMES
CORACIIFORMES
BUCEROTIFORMES
TROGONIFORMES
LEPTOSOMIFORMES
COLIIFORMES
ACCIPITRIFORMES
STRIGIFORMES
CHARADRIIFORMES
PELECANIFORMES
SULIFORMES
CICONIIFORMES
SPHENISCIFORMES
PROCELLARIIFORMES
GAVIIFORMES
EURYPYGIFORMES
PHAETHONTIFORMES
APODIFORMES
AEGOTHELIFORMES
PODARGIFORMES
STEATORNITHIFORMES
NYCTIBIIFORMES
CAPRIMULGIFORMES
GRUIFORMES
OPISTHOCOMIFORMES
OTIDIFORMES
MUSOPHAGIFORMES
CUCULIFORMES
PTEROCLIFORMES
MESITORNITHIFORMES
COLUMBIFORMES
PODICIPEDIFORMES
PHOENICOPTERIFORMES
GALLIFORMES
ANSERIFORMES
TINAMIFORMES
DINORNITHIFORMES
RHEIFORMES
APTERYGIFORMES
CASUARIIFORMES
STRUTHIONIFORMES
OUTGROUP
Group i – 399 taxa
Group ii – 1282 taxa
Group iii – 386 taxa
Group iv – 369 taxa
Group v – 162 taxa
Group vi – 162 taxa
Group i – 30 taxa
Group ii – 40 taxa
Group iii – 30 taxa
Group iv – 25 taxa
Group v – 12 taxa
Group vi – 13 taxa

### Figure S2

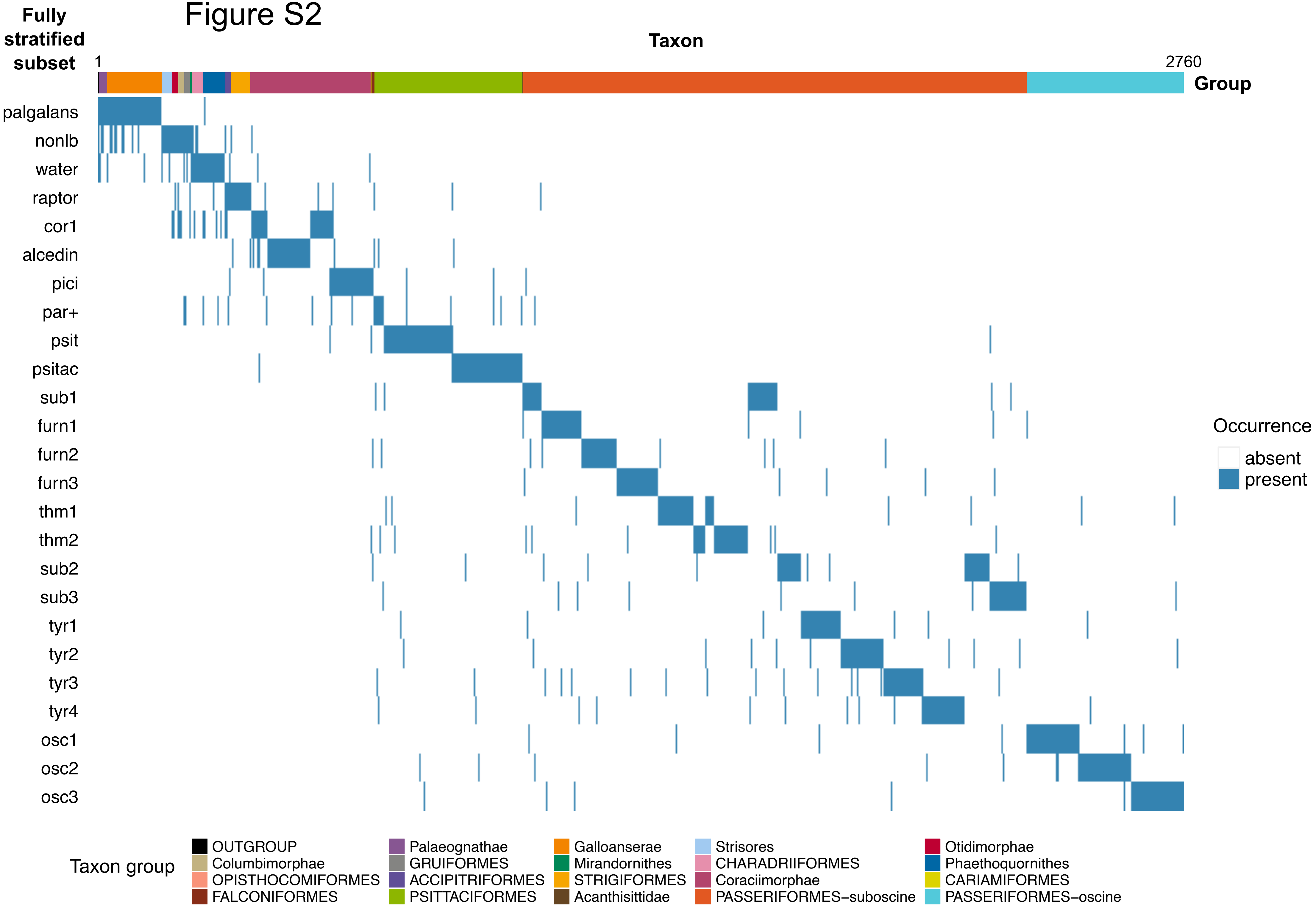

Fully stratified subset
Taxon
1
2760
Group
palgalans
nonlb
water
raptor
cor1
alcedin
pici
par+
psit
psitac
sub1
furn1
furn2
furn3
thm1
thm2
sub2
sub3
tyr1
tyr2
tyr3
tyr4
osc1
osc2
osc3
Occurrence
absent
present
Taxon group
OUTGROUP
Palaeognathae
Galloanserae
Strisores
Otidimorphae
Columbimorphae
GRUIFORMES
Mirandornithes
CHARADRIIFORMES
Phaethoquornithes
OPISTHOCOMIFORMES
ACCIPITRIFORMES
STRIGIFORMES
Coraciimorphae
CARIAMIFORMES
FALCONIFORMES
PSITTACIFORMES
Acanthisittidae
PASSERIFORMES−suboscine
PASSERIFORMES−oscine

Figure S3

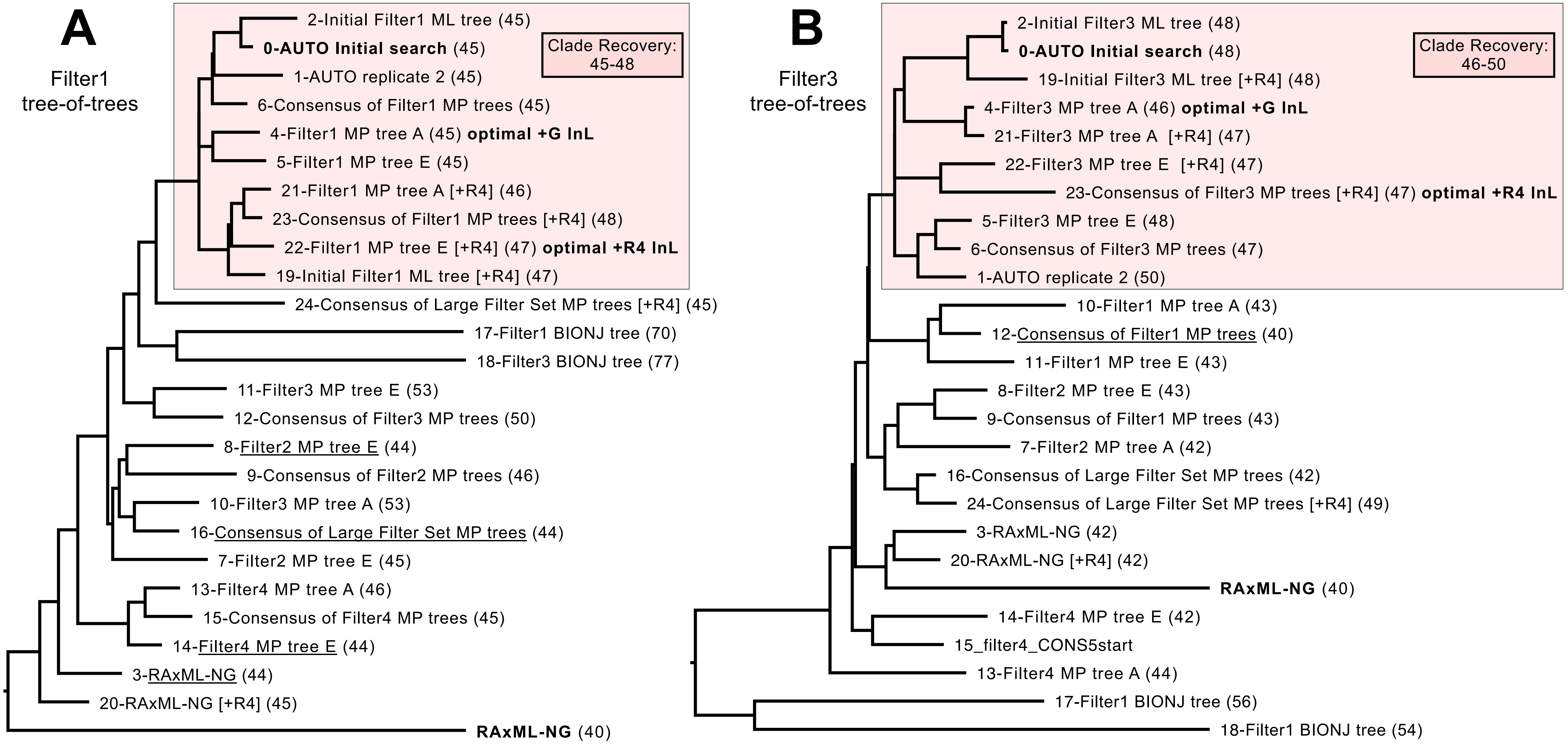
A
Filter1
tree-of-trees
Clade Recovery:
45-48
2-Initial Filter1 ML tree (45)
0-AUTO Initial search (45)
1-AUTO replicate 2 (45)
6-Consensus of Filter1 MP trees (45)
4-Filter1 MP tree A (45) optimal +G lnL
5-Filter1 MP tree E (45)
21-Filter1 MP tree A [+R4] (46)
23-Consensus of Filter1 MP trees [+R4] (48)
22-Filter1 MP tree E [+R4] (47) optimal +R4 lnL
19-Initial Filter1 ML tree [+R4] (47)
24-Consensus of Large Filter Set MP trees [+R4] (45)
17-Filter1 BIONJ tree (70)
18-Filter3 BIONJ tree (77)
11-Filter3 MP tree E (53)
12-Consensus of Filter3 MP trees (50)
8-Filter2 MP tree E (44)
9-Consensus of Filter2 MP trees (46)
10-Filter3 MP tree A (53)
16-Consensus of Large Filter Set MP trees (44)
7-Filter2 MP tree E (45)
13-Filter4 MP tree A (46)
15-Consensus of Filter4 MP trees (45)
14-Filter4 MP tree E (44)
3-RAxML-NG (44)
20-RAxML-NG [+R4] (45)
RAxML-NG (40)
B
Filter3
tree-of-trees
Clade Recovery:
46-50
2-Initial Filter3 ML tree (48)
0-AUTO Initial search (48)
19-Initial Filter3 ML tree [+R4] (48)
4-Filter3 MP tree A (46) optimal +G lnL
21-Filter3 MP tree A [+R4] (47)
22-Filter3 MP tree E [+R4] (47)
23-Consensus of Filter3 MP trees [+R4] (47) optimal +R4 lnL
5-Filter3 MP tree E (48)
6-Consensus of Filter3 MP trees (47)
1-AUTO replicate 2 (50)
10-Filter1 MP tree A (43)
12-Consensus of Filter1 MP trees (40)
11-Filter1 MP tree E (43)
8-Filter2 MP tree E (43)
9-Consensus of Filter1 MP trees (43)
7-Filter2 MP tree A (42)
16-Consensus of Large Filter Set MP trees (42)
24-Consensus of Large Filter Set MP trees [+R4] (49)
3-RAxML-NG (42)
20-RAxML-NG [+R4] (42)
RAxML-NG (40)
14-Filter4 MP tree E (42)
15_filter4_CONS5start
13-Filter4 MP tree A (44)
17-Filter1 BIONJ tree (56)
18-Filter1 BIONJ tree (54)

### Figure S4

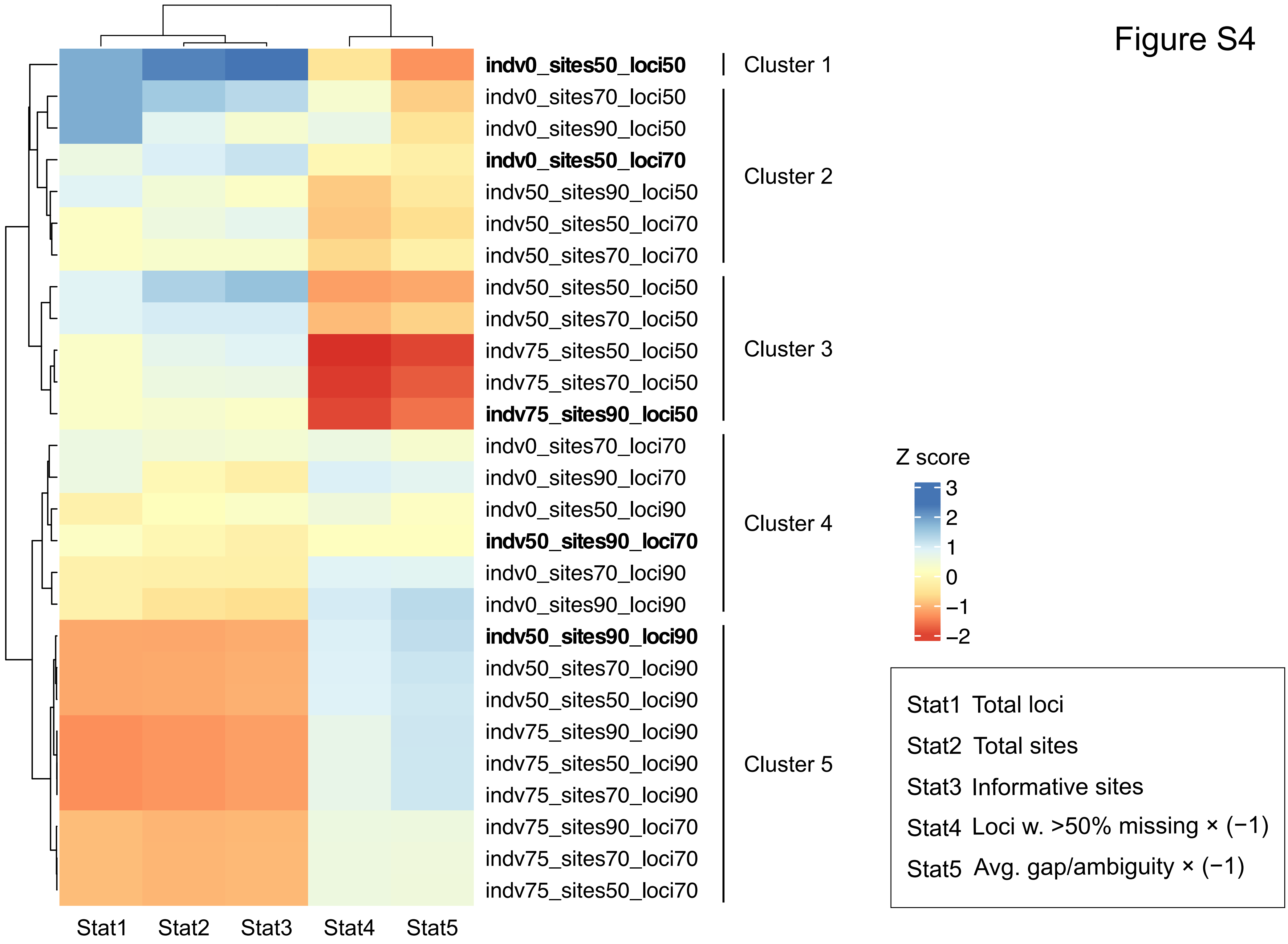
indv0_sites50_loci50
indv0_sites70_loci50
indv0_sites90_loci50
indv0_sites50_loci70
indv50_sites90_loci50
indv50_sites50_loci70
indv50_sites70_loci70
indv50_sites50_loci50
indv50_sites70_loci50
indv75_sites50_loci50
indv75_sites70_loci50
indv75_sites90_loci50
indv0_sites70_loci70
indv0_sites90_loci70
indv0_sites50_loci90
indv50_sites90_loci70
indv0_sites70_loci90
indv0_sites90_loci90
indv50_sites90_loci90
indv50_sites70_loci90
indv50_sites50_loci90
indv75_sites90_loci90
indv75_sites50_loci90
indv75_sites70_loci90
indv75_sites90_loci70
indv75_sites70_loci70
indv75_sites50_loci70
Cluster 1
Cluster 2
Cluster 3
Cluster 4
Cluster 5
Stat1
Stat2
Stat3
Stat4
Stat5
Z score
3
2
1
0
−1
−2
Stat1 Total loci
Stat2 Total sites
Stat3 Informative sites
Stat4 Loci w. >50% missing × (−1)
Stat5 Avg. gap/ambiguity × (−1)

### Figure S5

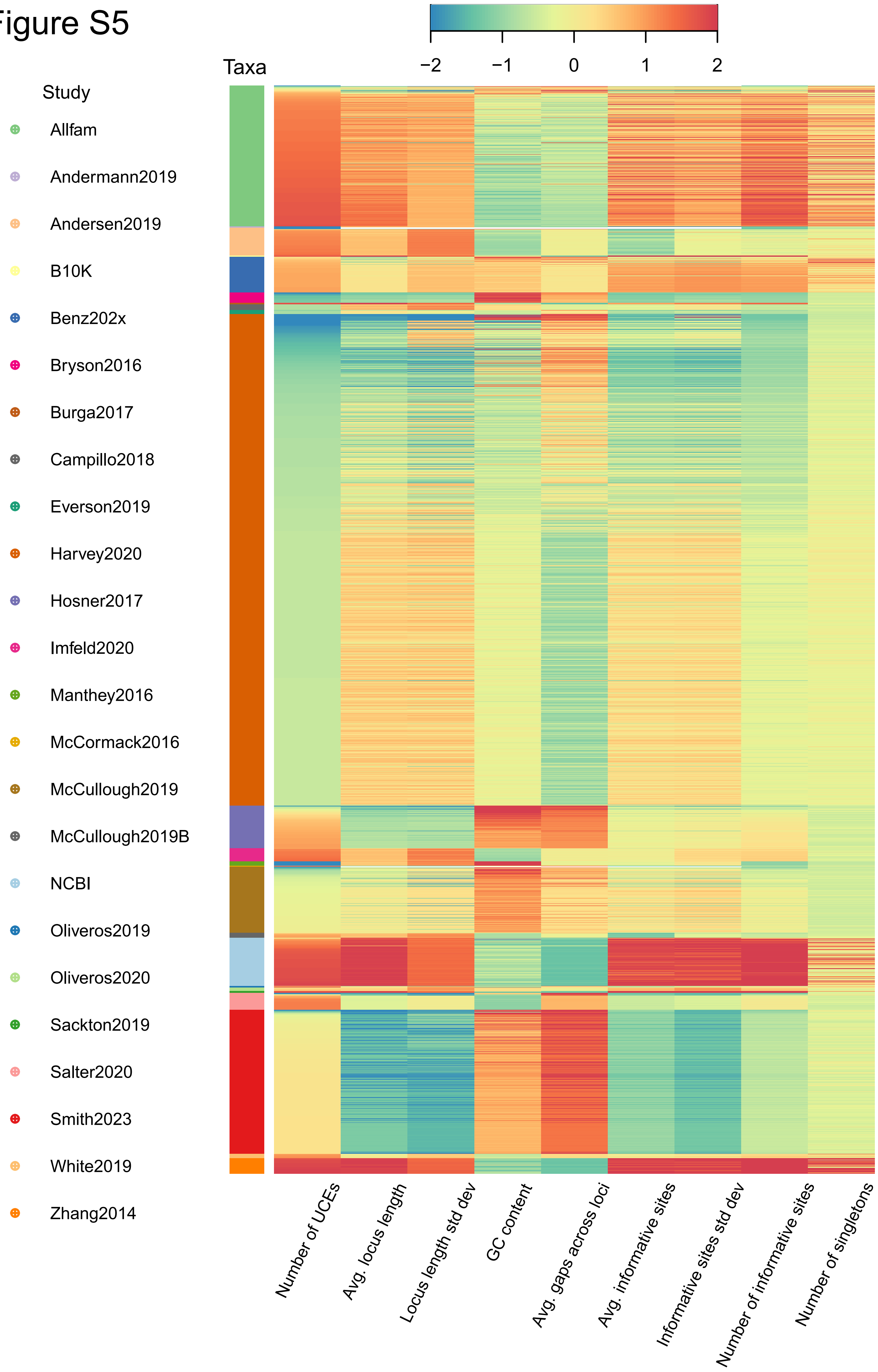

Study
Allfam
Andermann2019
Andersen2019
B10K
Benz202x
Bryson2016
Burga2017
Campillo2018
Everson2019
Harvey2020
Hosner2017
Imfeld2020
Manthey2016
McCormack2016
McCullough2019
McCullough2019B
NCBI
Oliveros2019
Oliveros2020
Sackton2019
Salter2020
Smith2023
White2019
Zhang2014
Taxa
−2
−1
0
1
2
Number of UCEs
Avg. locus length
Locus length std dev
GC content
Avg. gaps across loci
Avg. informative sites
Informative sites std dev
Number of informative sites
Number of singletons

### Figure S6

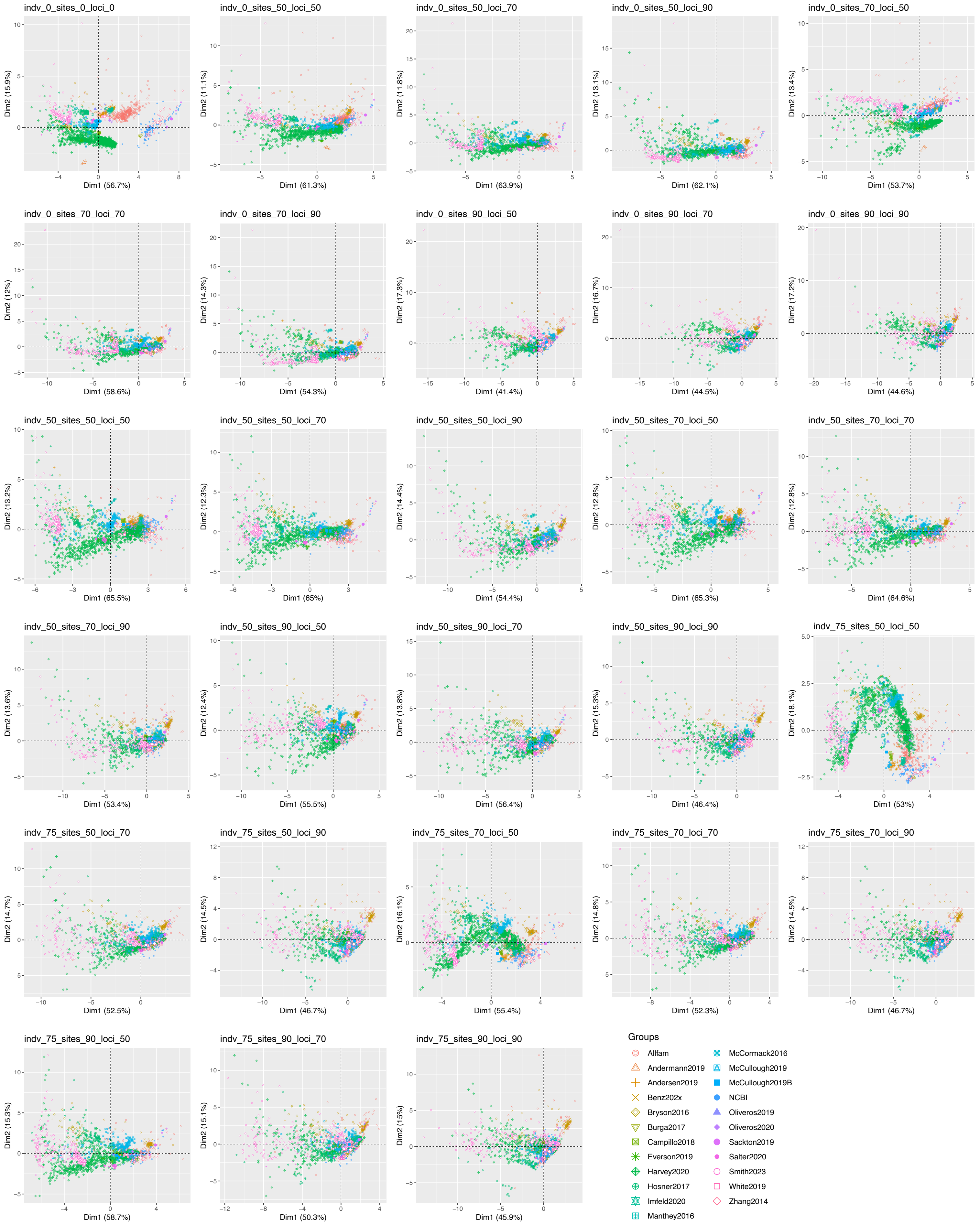
indv_0_sites_0_loci_0
Dim1 (56.7%)
Dim2 (15.9%)
indv_0_sites_50_loci_50
Dim1 (61.3%)
Dim2 (11.1%)
indv_0_sites_50_loci_70
Dim1 (63.9%)
Dim2 (11.8%)
indv_0_sites_50_loci_90
Dim1 (62.1%)
Dim2 (13.1%)
indv_0_sites_70_loci_50
Dim1 (53.7%)
Dim2 (13.4%)
indv_0_sites_70_loci_70
Dim1 (58.6%)
Dim2 (12%)
indv_0_sites_70_loci_90
Dim1 (54.3%)
Dim2 (14.3%)
indv_0_sites_90_loci_50
Dim1 (41.4%)
Dim2 (17.3%)
indv_0_sites_90_loci_70
Dim1 (44.5%)
Dim2 (16.7%)
indv_0_sites_90_loci_90
Dim1 (44.6%)
Dim2 (17.2%)
indv_50_sites_50_loci_50
Dim1 (65.5%)
Dim2 (13.2%)
indv_50_sites_50_loci_70
Dim1 (65%)
Dim2 (12.3%)
indv_50_sites_50_loci_90
Dim1 (54.4%)
Dim2 (14.4%)
indv_50_sites_70_loci_50
Dim1 (65.3%)
Dim2 (12.8%)
indv_50_sites_70_loci_70
Dim1 (64.6%)
Dim2 (12.8%)
indv_50_sites_70_loci_90
Dim1 (53.4%)
Dim2 (13.6%)
indv_50_sites_90_loci_50
Dim1 (55.5%)
Dim2 (12.4%)
indv_50_sites_90_loci_70
Dim1 (56.4%)
Dim2 (13.8%)
indv_50_sites_90_loci_90
Dim1 (46.4%)
Dim2 (15.3%)
indv_75_sites_50_loci_50
Dim1 (53%)
Dim2 (18.1%)
indv_75_sites_50_loci_70
Dim1 (52.5%)
Dim2 (14.7%)
indv_75_sites_50_loci_90
Dim1 (46.7%)
Dim2 (14.5%)
indv_75_sites_70_loci_50
Dim1 (55.4%)
Dim2 (16.1%)
indv_75_sites_70_loci_70
Dim1 (52.3%)
Dim2 (14.8%)
indv_75_sites_70_loci_90
Dim1 (46.7%)
Dim2 (14.5%)
indv_75_sites_90_loci_50
Dim1 (58.7%)
Dim2 (15.3%)
indv_75_sites_90_loci_70
Dim1 (50.3%)
Dim2 (15.1%)
indv_75_sites_90_loci_90
Dim1 (45.9%)
Dim2 (15%)
Groups
Allfam
Andermann2019
Andersen2019
Benz202x
Bryson2016
Burga2017
Campillo2018
Everson2019
Harvey2020
Hosner2017
Imfeld2020
Manthey2016
McCormack2016
McCullough2019
McCullough2019B
NCBI
Oliveros2019
Oliveros2020
Sackton2019
Salter2020
Smith2023
White2019
Zhang2014

### Figure S7

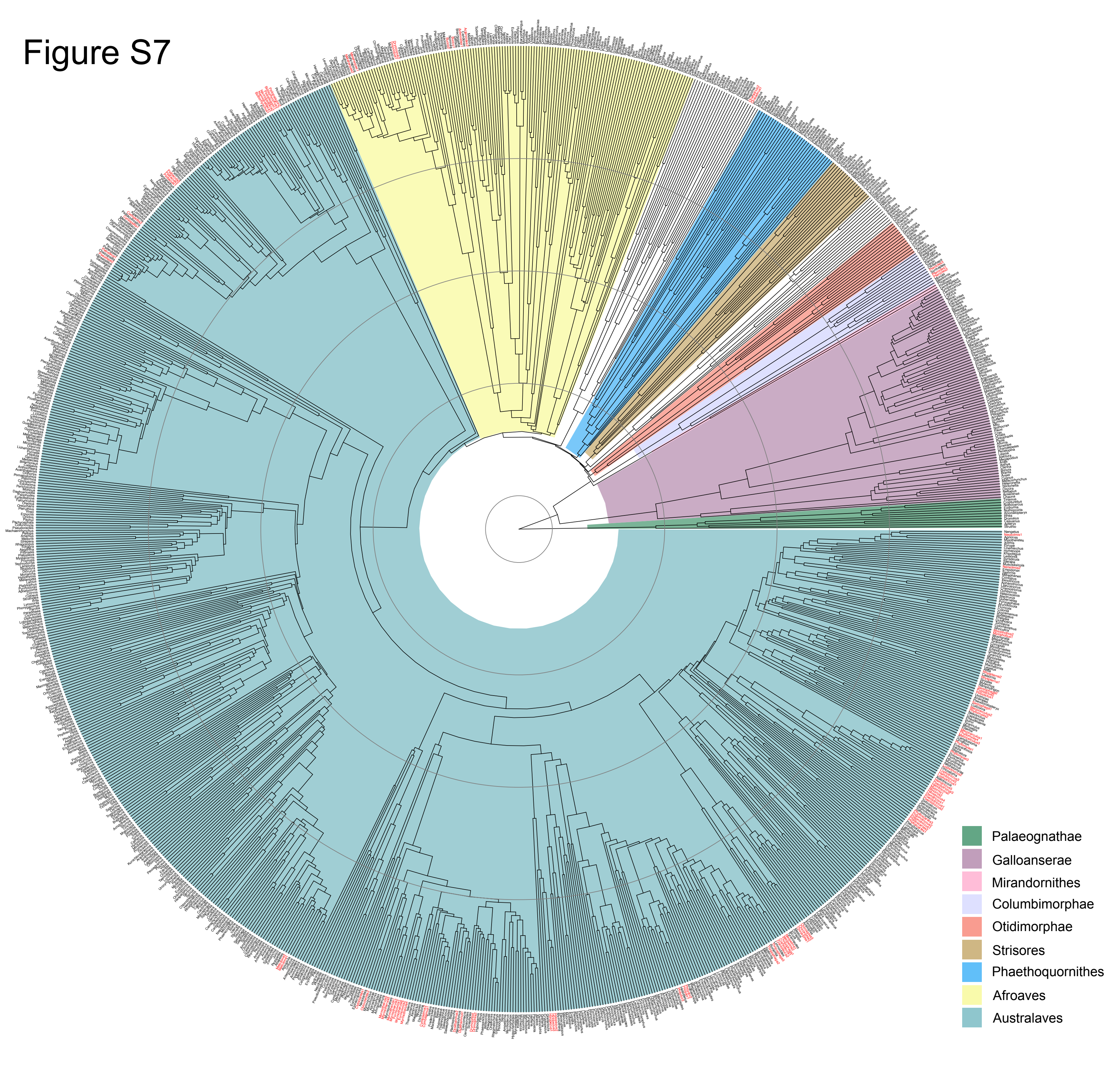
Palaeognathae
Galloanserae
Mirandornithes
Columbimorphae
Otidimorphae
Strisores
Phaethoquornithes
Afroaves
Australaves

Figure S8

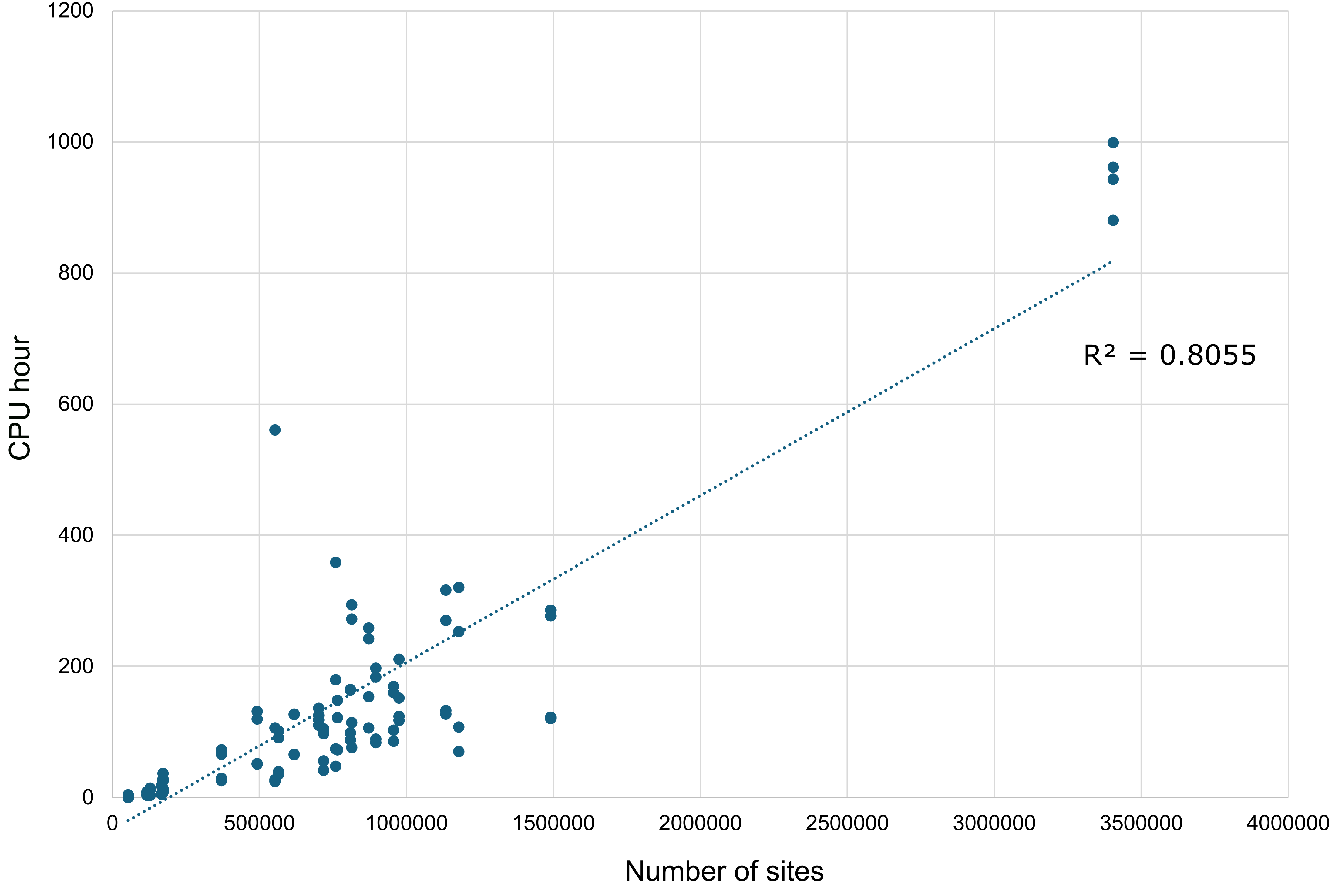
1200
1000
800
600
400
200
0
CPU hour
0
500000
1000000
1500000
2000000
2500000
3000000
3500000
4000000
Number of sites
R² = 0.8055

Figure S9

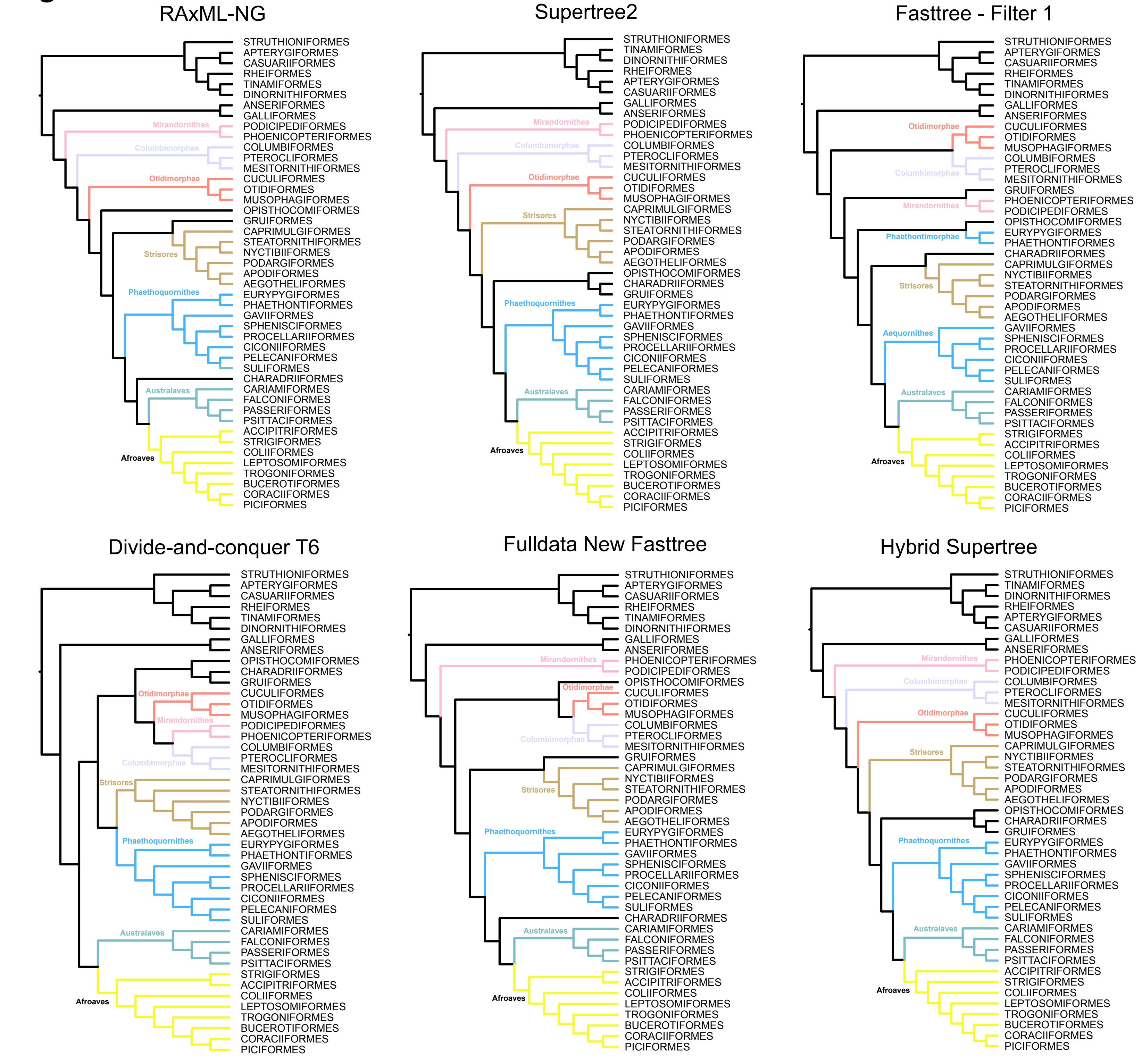

RAxML-NG
STRUTHIONIFORMES
APTERYGIFORMES
CASUARIIFORMES
RHEIFORMES
TINAMIFORMES
DINORNITHIFORMES
ANSERIFORMES
GALLIFORMES
Mirandornithes
PODICIPEDIFORMES
PHOENICOPTERIFORMES
Columbimorphae
COLUMBIFORMES
PTEROCLIFORMES
MESITORNITHIFORMES
Otidimorphae
CUCULIFORMES
OTIDIFORMES
MUSOPHAGIFORMES
OPISTHOCOMIFORMES
GRUIFORMES
CAPRIMULGIFORMES
STEATORNITHIFORMES
Strisores
NYCTIBIIFORMES
PODARGIFORMES
APODIFORMES
AEGOTHELIFORMES
Phaethoquornithes
EURYPYGIFORMES
PHAETHONTIFORMES
GAVIIFORMES
SPHENISCIFORMES
PROCELLARIIFORMES
CICONIIFORMES
PELECANIFORMES
SULIFORMES
CHARADRIIFORMES
Australaves
CARIAMIFORMES
FALCONIFORMES
PASSERIFORMES
PSITTACIFORMES
ACCIPITRIFORMES
STRIGIFORMES
Afroaves
COLIIFORMES
LEPTOSOMIFORMES
TROGONIFORMES
BUCEROTIFORMES
CORACIIFORMES
PICIFORMES
Supertree2
STRUTHIONIFORMES
TINAMIFORMES
DINORNITHIFORMES
RHEIFORMES
APTERYGIFORMES
CASUARIIFORMES
GALLIFORMES
ANSERIFORMES
Mirandornithes
PODICIPEDIFORMES
PHOENICOPTERIFORMES
Columbimorphae
COLUMBIFORMES
PTEROCLIFORMES
MESITORNITHIFORMES
Otidimorphae
CUCULIFORMES
OTIDIFORMES
MUSOPHAGIFORMES
Strisores
CAPRIMULGIFORMES
NYCTIBIIFORMES
STEATORNITHIFORMES
PODARGIFORMES
APODIFORMES
AEGOTHELIFORMES
OPISTHOCOMIFORMES
CHARADRIIFORMES
GRUIFORMES
Phaethoquornithes
EURYPYGIFORMES
PHAETHONTIFORMES
GAVIIFORMES
SPHENISCIFORMES
PROCELLARIIFORMES
CICONIIFORMES
PELECANIFORMES
SULIFORMES
Australaves
CARIAMIFORMES
FALCONIFORMES
PASSERIFORMES
PSITTACIFORMES
ACCIPITRIFORMES
STRIGIFORMES
Afroaves
COLIIFORMES
LEPTOSOMIFORMES
TROGONIFORMES
BUCEROTIFORMES
CORACIIFORMES
PICIFORMES
Fasttree - Filter 1
STRUTHIONIFORMES
APTERYGIFORMES
CASUARIIFORMES
RHEIFORMES
TINAMIFORMES
DINORNITHIFORMES
GALLIFORMES
ANSERIFORMES
Otidimorphae
CUCULIFORMES
OTIDIFORMES
MUSOPHAGIFORMES
COLUMBIFORMES
Columbimorphae
PTEROCLIFORMES
MESITORNITHIFORMES
GRUIFORMES
Mirandornithes
PHOENICOPTERIFORMES
PODICIPEDIFORMES
OPISTHOCOMIFORMES
Phaethontimorphae
EURYPYGIFORMES
PHAETHONTIFORMES
CHARADRIIFORMES
CAPRIMULGIFORMES
NYCTIBIIFORMES
Strisores
STEATORNITHIFORMES
PODARGIFORMES
APODIFORMES
AEGOTHELIFORMES
Aequornithes
GAVIIFORMES
SPHENISCIFORMES
PROCELLARIIFORMES
CICONIIFORMES
PELECANIFORMES
SULIFORMES
Australaves
CARIAMIFORMES
FALCONIFORMES
PASSERIFORMES
PSITTACIFORMES
STRIGIFORMES
ACCIPITRIFORMES
Afroaves
COLIIFORMES
LEPTOSOMIFORMES
TROGONIFORMES
BUCEROTIFORMES
CORACIIFORMES
PICIFORMES
Divide-and-conquer T6
STRUTHIONIFORMES
APTERYGIFORMES
CASUARIIFORMES
RHEIFORMES
TINAMIFORMES
DINORNITHIFORMES
GALLIFORMES
ANSERIFORMES
OPISTHOCOMIFORMES
CHARADRIIFORMES
GRUIFORMES
Otidimorphae
CUCULIFORMES
OTIDIFORMES
MUSOPHAGIFORMES
Mirandornithes
PODICIPEDIFORMES
PHOENICOPTERIFORMES
COLUMBIFORMES
PTEROCLIFORMES
Columbimorphae
MESITORNITHIFORMES
Strisores
CAPRIMULGIFORMES
STEATORNITHIFORMES
NYCTIBIIFORMES
PODARGIFORMES
APODIFORMES
AEGOTHELIFORMES
Phaethoquornithes
EURYPYGIFORMES
PHAETHONTIFORMES
GAVIIFORMES
SPHENISCIFORMES
PROCELLARIIFORMES
CICONIIFORMES
PELECANIFORMES
SULIFORMES
Australaves
CARIAMIFORMES
FALCONIFORMES
PASSERIFORMES
PSITTACIFORMES
STRIGIFORMES
ACCIPITRIFORMES
Afroaves
COLIIFORMES
LEPTOSOMIFORMES
TROGONIFORMES
BUCEROTIFORMES
CORACIIFORMES
PICIFORMES
Fulldata New Fasttree
STRUTHIONIFORMES
APTERYGIFORMES
CASUARIIFORMES
RHEIFORMES
TINAMIFORMES
DINORNITHIFORMES
GALLIFORMES
ANSERIFORMES
Mirandornithes
PHOENICOPTERIFORMES
PODICIPEDIFORMES
OPISTHOCOMIFORMES
Otidimorphae
CUCULIFORMES
OTIDIFORMES
MUSOPHAGIFORMES
COLUMBIFORMES
Columbimorphae
PTEROCLIFORMES
MESITORNITHIFORMES
GRUIFORMES
CAPRIMULGIFORMES
NYCTIBIIFORMES
Strisores
STEATORNITHIFORMES
PODARGIFORMES
APODIFORMES
AEGOTHELIFORMES
Phaethoquornithes
EURYPYGIFORMES
PHAETHONTIFORMES
GAVIIFORMES
SPHENISCIFORMES
PROCELLARIIFORMES
CICONIIFORMES
PELECANIFORMES
SULIFORMES
CHARADRIIFORMES
Australaves
CARIAMIFORMES
FALCONIFORMES
PASSERIFORMES
PSITTACIFORMES
STRIGIFORMES
ACCIPITRIFORMES
Afroaves
COLIIFORMES
LEPTOSOMIFORMES
TROGONIFORMES
BUCEROTIFORMES
CORACIIFORMES
PICIFORMES
Hybrid Supertree
STRUTHIONIFORMES
TINAMIFORMES
DINORNITHIFORMES
RHEIFORMES
APTERYGIFORMES
CASUARIIFORMES
GALLIFORMES
ANSERIFORMES
Mirandornithes
PHOENICOPTERIFORMES
PODICIPEDIFORMES
Columbimorphae
COLUMBIFORMES
PTEROCLIFORMES
MESITORNITHIFORMES
Otidimorphae
CUCULIFORMES
OTIDIFORMES
MUSOPHAGIFORMES
CAPRIMULGIFORMES
Strisores
NYCTIBIIFORMES
STEATORNITHIFORMES
PODARGIFORMES
APODIFORMES
AEGOTHELIFORMES
OPISTHOCOMIFORMES
CHARADRIIFORMES
GRUIFORMES
Phaethoquornithes
EURYPYGIFORMES
PHAETHONTIFORMES
GAVIIFORMES
SPHENISCIFORMES
PROCELLARIIFORMES
CICONIIFORMES
PELECANIFORMES
SULIFORMES
Australaves
CARIAMIFORMES
FALCONIFORMES
PASSERIFORMES
PSITTACIFORMES
ACCIPITRIFORMES
STRIGIFORMES
Afroaves
COLIIFORMES
LEPTOSOMIFORMES
TROGONIFORMES
BUCEROTIFORMES
CORACIIFORMES
PICIFORMES

**Table S1**. Source of phylogenomic data. References are available in the main text.

| Source | Study | Focal group | Taxa used |
|---|---|---|---|
| Allfam | (Braun et al., 2024) | Aves | 357 |
| Andermann2019 | (Andermann et al., 2019) | Trochilidae | 3 |
| Andersen2019 | (Andersen et al., 2019) | Meliphagidae | 68 |
| Benz202x | Benz et al. unpublished | Picidae | 91 |
| Bryson2016 | (Bryson et al., 2016) | Passerellidae | 26 |
| Burga2017 | (Burga et al., 2017) | Phalacrocorax | 4 |
| Campillo2018 | (Campillo et al., 2018) | Arachnothera | 14 |
| Everson2019 | (Everson et al., 2019) | Turdidae | 11 |
| Harvey2020 | (Harvey et al., 2020) | Suboscine | 1248 |
| Hosner2017 | (Hosner et al., 2016) | Phasianidae | 108 |
| Imfeld2020 | (Imfeld et al., 2020) | Fringillidae | 34 |
| Manthey2016 | (Manthey et al., 2016) | Piranga | 11 |
| McCormack2016 | (McCormack et al., 2016) | Aphelocoma | 2 |
| McCullough2019 | (McCullough et al., 2019) | Meliphagidae | 166 |
| McCullough2019B | (McCullough et al., 2019B) | Coraciiformes | 13 |
| NCBI | genomes | Aves | 125 |
| Oliveros2019 | (Oliveros et al., 2019) | Passerines | 4 |
| Oliveros2020 | (Oliveros et al., 2020) | Trogonidae | 9 |
| Sackton2019 | (Sackton et al., 2019) | Palaeognathae | 5 |
| Salter2020 | (Salter et al., 2020) | Strigidae | 43 |
| Smith2023 | (Smith et al., 2023) | Psittaciformes | 366 |
| White2019 | (White and Braun, 2019) | Strisores | 11 |
| Zhang2014 | (Zhang et al., 2014) | Aves | 40 |

**Table S2**. Summary statistics used in PCA for 28 datasets (full dataset and 27 filtered datasets). The full dataset and six filtered datasets (filter1 – filter6) were used in the initial fasttree analysis. Avg UCE, average number of UCEs per taxon; Avg length, UCE length averaged across all loci for each taxon and then averaged across all taxa, and its standard deviation (Length SD); Avg GC, GC content averaged across all loci for each taxon and then averaged across all taxa, and its standard deviation (GC SD); Avg gaps, number of missing data in gaps averaged across all loci for each taxon and then averaged across all taxa, and its standard deviation (Gaps SD); Avg IPI, the total number of individual-based parsimony informative sites per taxon (number of alternative alleles that a taxon carries that were present in two or more taxa in the dataset, assuming the outgroup as a reference) averaged across all taxa in the dataset, and its standard deviation (IPI SD); Total IPI, the total number of individual-based parsimony informative sites for all taxa in the dataset; Avg singletons, the total number of singletons in a dataset averaged across all taxa.

| Dataset | Used in initial analysis | Avg UCE | Avg length | Length SD | Avg GC | GC SD | Avg gaps | Gaps SD | Avg IPI | IPI SD | Total IPI | Avg singletons |
|---|---|---|---|---|---|---|---|---|---|---|---|---|
| **indv0_sites0_loci0** | Full dataset | 2946 | 440.5 | 138.4 | 0.39 | 0.05 | 156.7 | 125.6 | 43.4 | 38.4 | 138210.8 | 1310.0 |
| **indv0_sites50_loci50** | | 2340 | 405.9 | 114.9 | 0.4 | 0.05 | 71.6 | 69.6 | 36.0 | 32.1 | 86188.6 | 687.6 |
| **indv0_sites50_loci70** | Filter2 | 1734 | 408.0 | 115.6 | 0.4 | 0.05 | 73.2 | 67.6 | 36.2 | 31.8 | 64438.8 | 487.7 |
| **indv0_sites50_loci90** | | 1166 | 416.5 | 111.6 | 0.4 | 0.05 | 70.7 | 63.1 | 36.6 | 32.2 | 43300.7 | 313.3 |
| **indv0_sites70_loci50** | | 2340 | 349.0 | 79.9 | 0.4 | 0.05 | 30.0 | 36.1 | 25.1 | 21.7 | 59458.0 | 528.1 |
| **indv0_sites70_loci70** | | 1734 | 352.3 | 81.9 | 0.4 | 0.05 | 32.2 | 35.7 | 25.5 | 22.1 | 45033.3 | 374.6 |
| **indv0_sites70_loci90** | | 1166 | 357.3 | 74.2 | 0.4 | 0.05 | 30.2 | 31.1 | 25.4 | 21.7 | 29909.4 | 238.7 |
| **indv0_sites90_loci50** | Filter3 | 2340 | 276.2 | 50.8 | 0.4 | 0.05 | 9.0 | 14.2 | 14.3 | 13.1 | 33790.5 | 368.1 |
| **indv0_sites90_loci70** | | 1733 | 276.0 | 50.9 | 0.4 | 0.05 | 9.0 | 14.0 | 14.1 | 12.9 | 24525.2 | 253.4 |
| **indv0_sites90_loci90** | | 1166 | 283.3 | 45.1 | 0.41 | 0.05 | 9.0 | 13.0 | 14.0 | 12.8 | 16416.7 | 165.9 |
| **indv50_sites50_loci50** | Filter1 | 1679 | 427.0 | 125.0 | 0.4 | 0.05 | 60.4 | 64.9 | 38.3 | 34.2 | 69858.1 | 549.0 |
| **indv50_sites50_loci70** | Filter4 | 1321 | 420.4 | 129.9 | 0.4 | 0.05 | 55.2 | 61.0 | 37.9 | 35.0 | 53638.5 | 388.1 |
| **indv50_sites50_loci90** | | 361 | 306.4 | 73.1 | 0.4 | 0.04 | 13.8 | 22.3 | 16.4 | 16.1 | 5999.4 | 58.8 |
| **indv50_sites70_loci50** | | 1679 | 391.4 | 101.9 | 0.4 | 0.05 | 31.4 | 38.3 | 30.8 | 27.6 | 55304.4 | 459.6 |
| **indv50_sites70_loci70** | | 1321 | 386.5 | 105.6 | 0.4 | 0.05 | 29.0 | 35.8 | 30.7 | 28.1 | 42934.6 | 327.8 |
| **indv50_sites70_loci90** | | 361 | 303.7 | 67.8 | 0.4 | 0.04 | 12.1 | 18.1 | 15.8 | 14.7 | 5761.8 | 57.8 |
| **indv50_sites90_loci50** | | 1679 | 325.2 | 69.7 | 0.4 | 0.05 | 9.1 | 15.1 | 19.4 | 18.0 | 33822.8 | 330.3 |

| | | | | | | | | | | | | |
|---|---|---|---|---|---|---|---|---|---|---|---|---|
| **indv50_sites90_loci70** | | 1321 | 322.8 | 71.1 | 0.4 | 0.05 | 8.8 | 14.3 | 19.6 | 18.2 | 26714.6 | 239.0 |
| **indv50_sites90_loci90** | Filter6 | 361 | 284.9 | 52.9 | 0.4 | 0.05 | 8.0 | 11.7 | 13.0 | 11.9 | 4746.6 | 52.4 |
| **indv75_sites50_loci50** | | 1104 | 420.7 | 144.0 | 0.39 | 0.05 | 25.8 | 35.1 | 33.8 | 32.0 | 43037.6 | 380.4 |
| **indv75_sites50_loci70** | | 483 | 289.4 | 73.2 | 0.39 | 0.05 | 6.5 | 12.2 | 16.1 | 16.3 | 7900.5 | 76.1 |
| **indv75_sites50_loci90** | | 163 | 280.8 | 57.1 | 0.4 | 0.04 | 5.0 | 8.1 | 11.9 | 11.2 | 1975.6 | 22.3 |
| **indv75_sites70_loci50** | | 1104 | 406.9 | 132.7 | 0.39 | 0.05 | 15.8 | 23.7 | 30.6 | 29.0 | 38448.1 | 352.9 |
| **indv75_sites70_loci70** | | 483 | 289.0 | 72.2 | 0.39 | 0.05 | 6.3 | 11.3 | 15.9 | 15.9 | 7820.4 | 75.9 |
| **indv75_sites70_loci90** | | 163 | 280.8 | 57.1 | 0.4 | 0.04 | 5.0 | 8.1 | 11.9 | 11.2 | 1975.6 | 22.3 |
| **indv75_sites90_loci50** | | 1104 | 375.3 | 111.2 | 0.39 | 0.05 | 6.8 | 11.9 | 24.0 | 22.5 | 29412.3 | 300.6 |
| **indv75_sites90_loci70** | Filter5 | 483 | 283.6 | 68.2 | 0.39 | 0.05 | 5.1 | 9.0 | 14.7 | 14.1 | 7210.3 | 73.4 |
| **indv75_sites90_loci90** | | 163 | 279.7 | 56.4 | 0.4 | 0.04 | 4.8 | 7.8 | 11.6 | 10.7 | 1916.4 | 22.1 |

**Table S3**. Summary statistics for all datasets using locus-based summary statistics.

| Dataset | Filtering scheme | | | Loci number | Total sites | Informative sites | Avg. gap % | #taxa missing50% |
|---|---|---|---|---|---|---|---|---|
| | Indv | Sites | Loci | | | | | |
| **i0s50l50** | 0 | 50 | 50 | 3140 | 1490832 | 810880 | 35.3229 | 575 |
| **i0s50l70** | 0 | 50 | 70 | 1988 | 956470 | 529425 | 24.6981 | 442 |
| **i0s50l90** | 0 | 50 | 90 | 1269 | 617523 | 336777 | 20.5238 | 273 |
| **i0s70l50** | 0 | 70 | 50 | 3140 | 1177750 | 561991 | 29.9888 | 321 |
| **i0s70l70** | 0 | 70 | 70 | 1988 | 764867 | 373482 | 19.0502 | 247 |
| **i0s70l90** | 0 | 70 | 90 | 1269 | 491069 | 234289 | 14.3653 | 167 |
| **i0s90l50** | 0 | 90 | 50 | 3140 | 895617 | 365026 | 27.2780 | 227 |
| **i0s90l70** | 0 | 90 | 70 | 1987 | 564595 | 231975 | 14.6610 | 146 |
| **i0s90l90** | 0 | 90 | 90 | 1269 | 370371 | 149909 | 10.1489 | 125 |
| **i50s50l50** | 50 | 50 | 50 | 2222 | 1133804 | 634424 | 33.4680 | 796 |
| **i50s50l70** | 50 | 50 | 70 | 1634 | 808882 | 445609 | 28.3897 | 687 |
| **i50s50l90** | 50 | 50 | 90 | 397 | 127221 | 54172 | 12.1369 | 154 |
| **i50s70l50** | 50 | 70 | 50 | 2222 | 974352 | 499797 | 29.6783 | 717 |
| **i50s70l70** | 50 | 70 | 70 | 1634 | 700861 | 353731 | 24.5368 | 633 |
| **i50s70l90** | 50 | 70 | 90 | 397 | 125435 | 52633 | 11.6569 | 152 |
| **i50s90l50** | 50 | 90 | 50 | 2222 | 758682 | 335019 | 26.2277 | 675 |
| **i50s90l70** | 50 | 90 | 70 | 1634 | 552210 | 240359 | 21.0098 | 382 |
| **i50s90l90** | 50 | 90 | 90 | 397 | 116204 | 46386 | 10.6428 | 145 |
| **i75s50l50** | 75 | 50 | 50 | 1681 | 871630 | 470904 | 41.6242 | 1079 |
| **i75s50l70** | 75 | 50 | 70 | 575 | 171367 | 72580 | 17.3646 | 255 |
| **i75s50l90** | 75 | 50 | 90 | 184 | 52839 | 19327 | 11.9425 | 218 |
| **i75s70l50** | 75 | 70 | 50 | 1681 | 813133 | 420938 | 39.8546 | 1053 |
| **i75s70l70** | 75 | 70 | 70 | 575 | 170911 | 72144 | 17.2674 | 255 |
| **i75s70l90** | identical to i75s50l90 | | | | | | | |
| **i75s90l50** | 75 | 90 | 50 | 1681 | 718041 | 341080 | 37.9833 | 1020 |
| **i75s90l70** | 75 | 90 | 70 | 575 | 166784 | 68908 | 16.8756 | 249 |
| **i75s90l90** | 75 | 90 | 90 | 184 | 52617 | 19132 | 11.8960 | 218 |
| **Full dataset** | 0 | 0 | 0 | 5121 | 3403758 | 2047668 | na | na |

**Table S4**. Phylogenomic trees used in the supertree analysis.

| Tree name | Data type | Weight in MPR | Taxon group | Taxa number before trim | Reference |
|---|---|---|---|---|---|
| AllFam | UCE | 4 | All birds | 396 | (Braun et al., 2024) |
| Andermann2018_Rooted | UCE | 4 | Trochilidae | 3 | (Andermann et al., 2019) |
| Andersen_2019_Meliphagidae_raxml | UCE | 2 | Meliphagidae | 59 | (Andersen et al., 2019) |
| Andersen_2019_Meliphagidae_svdq | | 2 | | | |
| Anderson2018_RAxML_3249 | UCE | 4 | Alcedinidae | 21 | (Andersen et al., 2018) |
| Benz_unpublished_Picidae | UCE | 4 | Picidae | 95 | Benz et al. unpublished |
| Bruxaux2017 | UCE | 4 | Goura | 6 | (Bruxaux et al., 2018) |
| Bryson2016_RAxML | UCE | 4 | Passerellidae | 31 | (Bryson et al., 2016) |
| Burga2017 | WGS | 8 | Phalacrocorax | 7 | (Burga et al., 2017) |
| Campillo2017_ASTRAL | UCE | 4 | Arachnothera | 17 | (Campillo et al., 2018) |
| Catanach_2021_Accipiter | UCE | 4 | Accipiter | 5 | (Catanach et al., 2021) |
| Chen2018 | UCE | 4 | Phasianidae | 27 | (Chen et al., 2018) |
| Everson_2019_Turdidae_DRAW | UCE | 4 | Turdidae | 11 | (Everson et al., 2019) |
| Ferreira2018 | UCE | 4 | Piciformes | 3 | (Ferreira et al., 2018) |
| Harvey_2020_Suboscine | UCE | 4 | Suboscine | 1287 | (Harvey et al., 2020) |
| Hosner2017 | UCE | 4 | Phasianidae | 114 | (Hosner et al., 2016) |
| Imfeld_2020ml_Fringillidae | UCE | 2 | Fringillidae | 36 | (Imfeld et al., 2020) |
| Imfeld_2020msc_Fringillidae | | 2 | | | |
| JarvisTENT_rooted | WGS | 8 | Neornithes | 48 | (Jarvis et al., 2014) |
| JarvisUCE | UCE | 4 | | | |
| Kirchman2021ml_Ralloidea | UCE | 2 | Ralloidea | 65 | (Kirchman et al., 2021) |
| Kirchman2021msc_Ralloidea | | 2 | | | |
| Lamichhaney2015 | WGS | 8 | Geospiza | 20 | (Lamichhaney et al., 2015) |
| Manthy2016_ASTRALpruned | UCE | 4 | Piranga | 12 | (Manthey et al., 2016) |
| McCormack2013_1541uce | UCE | 2 | Neoaves | 33 | (McCormack et al., 2016) |
| McCormack2013_416uce | | 2 | | | |

| | | | | | |
|---|---|---|---|---|---|
| McCullough_2019_Coraciiformes | UCE | 4 | Coraciiformes | 192 | (McCullough et al., 2019a) |
| McCullough_2019astral_Meliphagidae | UCE | 2 | Meliphagidae | 16 | (McCullough et al., 2019b) |
| McCullough_2019ml_Meliphagidae | | 2 | | | |
| McCullough_2022_corvides | UCE | 4 | Corvides | 727 | (McCullough et al., 2022) |
| Musher2018_Fig5_ASTRAL | UCE | 4 | Pachyramphus | 18 | (Musher and Cracraft, 2018) |
| Nater2015 | UCE | 4 | Ficedula | 6 | (Nater et al., 2015) |
| Oliveros_2019_Trogonidae_DRAW | UCE | 4 | Trogonidae | 18 | (Oliveros et al., 2020) |
| Oliveros_2020_Passerines | UCE | 4 | Passerines | 221 | (Oliveros et al., 2019) |
| Oliveros_2021_White-eyes | UCE | 4 | White-eyes | 71 | (Oliveros et al., 2021) |
| Ottenburghs2016 | WGS | 8 | Anatidae | 19 | (Ottenburghs et al., 2016) |
| Prum_Alldata_RY | AHE | 3 | Neornithes | 200 | (Prum et al., 2015) |
| Reddy2017_Fig3 | LEGACY | 2 | Neornithes | 235 | (Reddy et al., 2017) |
| Sackton2018 | WGS | 8 | Palaeognathae | 15 | (Sackton et al., 2019) |
| Salter_2020ml_Strigidae | UCE | 2 | Strigidae | 48 | (Salter et al., 2020) |
| Salter_2020msc_Strigidae | | 2 | | | |
| Smith_2022_Psittaciformes | UCE | 4 | Psittaciformes | 385 | (Smith et al., 2023) |
| Smith2018_BiorXiv | UCE | 4 | Psittaculidae | 54 | (Smith et al., 2020) |
| Vianna_2020_Penguins | UCE | 4 | Penguins | 19 | (Vianna et al., 2020) |
| Vinay_2022_Strigidae | UCE | 4 | Strigidae | 51 | (Vinay et al., 2022) |
| Wang2017_partRAxMLpruned | UCE | 4 | Phasianidae | 20 | (Wang et al., 2017) |
| Wang2022_Palaeognathae | UCE | 4 | Palaeognathae | 16 | (Wang et al., 2022) |
| White2017_GARLI_75trim | UCE | 4 | Nyctibiidae | 12 | (White et al., 2017) |
| White_2019_Strisores | UCE | 4 | Strisores | 23 | (White and Braun, 2019) |
| Yonezawa2017_IOC_Pruned | LEGACY | 2 | Palaeognathae | 25 | (Yonezawa et al., 2017) |
| Younger2018_Rooted | UCE | 4 | Newtonia | 4 | (Younger et al., 2018) |
| Zarza2016_Rooted | UCE | 4 | Aphelocoma | 3 | (Zarza et al., 2016) |
| Zhao_2022_Pipridae | UCE+introns | 6 | Pipridae | 19 | (Zhao et al., 2023) |
| Burleigh et al BigBird | LEGACY | 1 | All birds | 6697 | (Burleigh et al., 2015) |
| Jetz | LEGACY + taxonomy | 1 | All birds | 9979 | (Jetz et al., 2012) |

**Table S5**. Log likelihoods and credible clade recovery of the initial fasttree analyses.

| Filter | Dataset | sites | PI sites | MP lnL | BIONJ lnL | MP-BIONJ lnL |
|---|---|---|---|---|---|---|
| | Full | 3403758 | 2047980 | -160487114 | -165308430 | 4821316.36 |
| 1 | Indv50_sites50_loci50 | 1133804 | 634509 | -62680545 | -65125877 | 2445332.44 |
| 2 | Indv0_sites50_loci70 | 956470 | 529511 | -53200569 | -55169666 | 1969097.21 |
| 3 | Indv0_sites90_loci50 | 895617 | 365126 | -27170600 | -27700991 | 530390.474 |
| 4 | Indv50_sites50_loci70 | 808882 | 445674 | -45197017 | -46987653 | 1790635.89 |
| 5 | Indv75_sites90_loci70 | 166784 | 68923 | -5370575.2 | -5505755.1 | 135179.915 |
| 6 | Indv50_sites90_loci90 | 116204 | 46395 | -3532340.6 | -3623823 | 91482.404 |

| Filter | Dataset | Optimal lnL | Opt-Start lnL | MP search | Param (MP) | MLdist |
|---|---|---|---|---|---|---|
| | Full | -160472895 | 14218.66 | 6466.063 | 138796.681 | 4651448.53 |
| 1 | Indv50_sites50_loci50 | -62675001 | 5543.794 | 1768.67 | 19767.175 | 976617.111 |
| 2 | Indv0_sites50_loci70 | -53195848 | 4721.011 | 1556.255 | 13646.366 | 960605.84 |
| 3 | Indv0_sites90_loci50 | -27167864 | 2735.599 | 987.054 | 9867.174 | 512446.15 |
| 4 | Indv50_sites50_loci70 | -45193006 | 4010.83 | 1261.921 | 11227.571 | 715865.01 |
| 5 | Indv75_sites90_loci70 | -5369723 | 852.202 | 218.43 | 922.282 | 131329.254 |
| 6 | Indv50_sites90_loci90 | -3531574.1 | 766.538 | 137.615 | 625.773 | 108417.459 |

| Filter | Dataset | BIONJ | BIONJ/MP | ML treesearch | Total CPU time | total time (h:m:s) |
|---|---|---|---|---|---|---|
| | Full | 54.514 | 32.0213079 | 1773911.89 | 7771185.08 | 2158h:39m:45s |
| 1 | Indv50_sites50_loci50 | 106.634 | 45.3533978 | 487579.415 | 2876026.29 | 798h:53m:46s |
| 2 | Indv0_sites50_loci70 | 108.034 | 63.1939633 | 347897.003 | 2299717.76 | 638h:48m:37s |
| 3 | Indv0_sites90_loci50 | 56.254 | 47.2168453 | 211235.092 | 1771074.01 | 491h:57m:54s |
| 4 | Indv50_sites50_loci70 | 111.268 | 57.326293 | 335672.793 | 1788646.82 | 496h:50m:46s |
| 5 | Indv75_sites90_loci70 | 130.827 | 115.243884 | 24257.007 | 223807.34 | 62h:10m:7s |
| 6 | Indv50_sites90_loci90 | 103.588 | 142.15713 | 12534.583 | 162515.019 | 45h:8m:35s |

| Filter | Dataset | Non-monophyletic group | | | | |
|---|---|---|---|---|---|---|
| | | Order | Family | Genus | High-level | sum |
| | Full | 0 | 2 | 38 | 10 | 50 |
| 1 | Indv50_sites50_loci50 | 0 | 2 | 41 | 1 | 44 |
| 2 | Indv0_sites50_loci70 | 0 | 2 | 39 | 3 | 44 |
| 3 | Indv0_sites90_loci50 | 1 | 1 | 37 | 8 | 47 |
| 4 | Indv50_sites50_loci70 | 0 | 2 | 42 | 2 | 46 |
| 5 | Indv75_sites90_loci70 | 0 | 2 | 46 | 4 | 52 |
| 6 | Indv50_sites90_loci90 | 0 | 3 | 41 | 5 | 49 |

**Table S6**. Summary of credible clade recovery of all new fasttrees.

| dataset | Non-monophyletic group | | | | | model | MP starting tree |
|---|---|---|---|---|---|---|---|
| | order | family | genus | High-level | sum | | |
| fulldata | 0 | 2 | 39 | 2 | 43 | GTR+G | parsA |
| fulldata | 0 | 2 | 40 | 1 | 43 | GTR+G | parsB |
| fulldata | 0 | 2 | 38 | 3 | 43 | GTR+R4 | parsC |
| fulldata | 0 | 2 | 38 | 0 | 40 | GTR+R4 | parsD |
| i0s50l50 | 1 | 3 | 38 | 5 | 47 | GTR+G | parsA |
| i0s50l50 | 0 | 2 | 39 | 1 | 42 | GTR+G | parsB |
| i0s50l50 | 1 | 3 | 38 | 5 | 47 | GTR+R4 | parsC |
| i0s50l50 | 0 | 2 | 39 | 2 | 43 | GTR+R4 | parsD |
| i0s50l70 | 0 | 2 | 40 | 2 | 44 | GTR+G | parsA |
| i0s50l70 | 0 | 2 | 39 | 3 | 44 | GTR+G | parsB |
| i0s50l70 | 0 | 2 | 39 | 1 | 42 | GTR+R4 | parsC |
| i0s50l70 | 0 | 2 | 39 | 2 | 43 | GTR+R4 | parsD |
| i0s50l90 | 0 | 2 | 41 | 5 | 48 | GTR+G | parsA |
| i0s50l90 | 0 | 2 | 42 | 5 | 49 | GTR+G | parsB |
| i0s50l90 | 0 | 2 | 41 | 3 | 46 | GTR+R4 | parsC |
| i0s50l90 | 0 | 2 | 41 | 3 | 46 | GTR+R4 | parsD |
| i0s70l50 | 1 | 3 | 39 | 7 | 50 | GTR+G | parsA |
| i0s70l50 | 1 | 3 | 38 | 7 | 49 | GTR+G | parsB |
| i0s70l50 | 1 | 3 | 38 | 8 | 50 | GTR+R4 | parsC |
| i0s70l50 | 1 | 3 | 39 | 8 | 51 | GTR+R4 | parsD |
| i0s70l70 | 0 | 2 | 40 | 5 | 47 | GTR+G | parsA |
| i0s70l70 | 0 | 2 | 40 | 6 | 48 | GTR+G | parsB |
| i0s70l70 | 0 | 2 | 40 | 3 | 45 | GTR+R4 | parsC |
| i0s70l70 | 0 | 2 | 40 | 5 | 47 | GTR+R4 | parsD |
| i0s70l90 | 1 | 3 | 40 | 9 | 53 | GTR+G | parsA |
| i0s70l90 | 1 | 3 | 41 | 8 | 53 | GTR+G | parsB |
| i0s70l90 | 1 | 3 | 41 | 9 | 54 | GTR+R4 | parsC |
| i0s70l90 | 1 | 3 | 41 | 9 | 54 | GTR+R4 | parsD |
| i0s90l50 | 1 | 1 | 37 | 7 | 46 | GTR+G | parsA |
| i0s90l50 | 1 | 1 | 37 | 9 | 48 | GTR+G | parsB |
| i0s90l50 | 1 | 1 | 37 | 6 | 45 | GTR+R4 | parsC |
| i0s90l50 | 1 | 1 | 37 | 7 | 46 | GTR+R4 | parsD |
| i0s90l70 | 1 | 1 | 42 | 10 | 54 | GTR+G | parsA |
| i0s90l70 | 1 | 2 | 40 | 9 | 52 | GTR+G | parsB |
| i0s90l70 | 1 | 1 | 41 | 8 | 51 | GTR+R4 | parsC |

| | | | | | | | |
|---|---|---|---|---|---|---|---|
| i0s90l70 | 1 | 1 | 40 | 9 | 51 | GTR+R4 | parsD |
| i0s90l90 | 1 | 1 | 43 | 8 | 53 | GTR+G | parsA |
| i0s90l90 | 1 | 1 | 43 | 9 | 54 | GTR+G | parsB |
| i0s90l90 | 1 | 1 | 43 | 8 | 53 | GTR+R4 | parsC |
| i0s90l90 | 1 | 1 | 43 | 7 | 52 | GTR+R4 | parsD |
| i50s50l50 | 0 | 2 | 40 | 1 | 43 | GTR+G | parsA |
| i50s50l50 | 0 | 2 | 40 | 1 | 43 | GTR+G | parsB |
| i50s50l50 | 0 | 2 | 42 | 1 | 45 | GTR+R4 | parsC |
| i50s50l50 | 0 | 2 | 42 | 2 | 46 | GTR+R4 | parsD |
| i50s50l70 | 0 | 2 | 41 | 3 | 46 | GTR+G | parsA |
| i50s50l70 | 0 | 2 | 41 | 2 | 45 | GTR+G | parsB |
| i50s50l70 | 0 | 2 | 44 | 2 | 48 | GTR+R4 | parsC |
| i50s50l70 | 0 | 2 | 42 | 4 | 48 | GTR+R4 | parsD |
| i50s50l90 | 0 | 3 | 46 | 7 | 56 | GTR+G | parsA |
| i50s50l90 | 0 | 3 | 45 | 6 | 54 | GTR+G | parsB |
| i50s50l90 | 0 | 3 | 46 | 6 | 55 | GTR+R4 | parsC |
| i50s50l90 | 0 | 3 | 45 | 5 | 53 | GTR+R4 | parsD |
| i50s70l50 | 0 | 2 | 40 | 3 | 45 | GTR+G | parsA |
| i50s70l50 | 0 | 2 | 41 | 2 | 45 | GTR+G | parsB |
| i50s70l50 | 0 | 2 | 41 | 4 | 47 | GTR+R4 | parsC |
| i50s70l50 | 0 | 2 | 42 | 2 | 46 | GTR+R4 | parsD |
| i50s70l70 | 0 | 2 | 41 | 3 | 46 | GTR+G | parsA |
| i50s70l70 | 0 | 2 | 41 | 3 | 46 | GTR+G | parsB |
| i50s70l70 | 0 | 2 | 42 | 3 | 47 | GTR+R4 | parsC |
| i50s70l70 | 0 | 2 | 42 | 4 | 48 | GTR+R4 | parsD |
| i50s70l90 | 0 | 3 | 45 | 6 | 54 | GTR+G | parsA |
| i50s70l90 | 0 | 4 | 47 | 5 | 56 | GTR+G | parsB |
| i50s70l90 | 0 | 2 | 44 | 6 | 52 | GTR+R4 | parsC |
| i50s70l90 | 0 | 3 | 47 | 5 | 55 | GTR+R4 | parsD |
| i50s90l50 | 0 | 2 | 41 | 5 | 48 | GTR+G | parsA |
| i50s90l50 | 0 | 1 | 41 | 5 | 47 | GTR+G | parsB |
| i50s90l50 | 0 | 1 | 41 | 4 | 46 | GTR+R4 | parsC |
| i50s90l50 | 0 | 1 | 41 | 6 | 48 | GTR+R4 | parsD |
| i50s90l70 | 0 | 1 | 43 | 4 | 48 | GTR+G | parsA |
| i50s90l70 | 0 | 1 | 45 | 5 | 51 | GTR+G | parsB |
| i50s90l70 | 0 | 1 | 44 | 5 | 50 | GTR+R4 | parsC |
| i50s90l70 | 0 | 0 | 43 | 4 | 47 | GTR+R4 | parsD |
| i50s90l90 | 0 | 3 | 41 | 5 | 49 | GTR+G | parsA |
| i50s90l90 | 0 | 3 | 42 | 5 | 50 | GTR+G | parsB |

| | | | | | | | |
|---|---|---|---|---|---|---|---|
| i50s90l90 | 0 | 2 | 42 | 4 | 48 | GTR+R4 | parsC |
| i50s90l90 | 0 | 1 | 42 | 5 | 48 | GTR+R4 | parsD |
| i75s50l50 | 0 | 2 | 45 | 3 | 50 | GTR+G | parsA |
| i75s50l50 | 0 | 2 | 44 | 2 | 48 | GTR+G | parsB |
| i75s50l50 | 0 | 2 | 46 | 2 | 50 | GTR+R4 | parsC |
| i75s50l50 | 0 | 2 | 47 | 5 | 54 | GTR+R4 | parsD |
| i75s50l70 | 0 | 2 | 45 | 5 | 52 | GTR+G | parsA |
| i75s50l70 | 0 | 3 | 44 | 6 | 53 | GTR+G | parsB |
| i75s50l70 | 0 | 3 | 47 | 7 | 57 | GTR+R4 | parsC |
| i75s50l70 | 0 | 3 | 49 | 6 | 58 | GTR+R4 | parsD |
| i75s50l90 | 3 | 4 | 55 | 15 | 77 | GTR+G | parsA |
| i75s50l90 | 4 | 6 | 58 | 18 | 86 | GTR+G | parsB |
| i75s50l90 | 3 | 6 | 56 | 18 | 83 | GTR+R4 | parsC |
| i75s50l90 | 3 | 5 | 55 | 18 | 81 | GTR+R4 | parsD |
| i75s70l50 | 0 | 3 | 43 | 3 | 49 | GTR+G | parsA |
| i75s70l50 | 0 | 2 | 44 | 4 | 50 | GTR+G | parsB |
| i75s70l50 | 0 | 2 | 42 | 2 | 46 | GTR+R4 | parsC |
| i75s70l50 | 0 | 2 | 43 | 2 | 47 | GTR+R4 | parsD |
| i75s70l70 | 0 | 3 | 47 | 6 | 56 | GTR+G | parsA |
| i75s70l70 | 0 | 3 | 48 | 6 | 57 | GTR+G | parsB |
| i75s70l70 | 0 | 3 | 47 | 4 | 54 | GTR+R4 | parsC |
| i75s70l70 | 0 | 3 | 48 | 7 | 58 | GTR+R4 | parsD |
| i75s90l50 | 0 | 2 | 42 | 6 | 50 | GTR+G | parsA |
| i75s90l50 | 0 | 2 | 43 | 5 | 50 | GTR+G | parsB |
| i75s90l50 | 0 | 2 | 40 | 3 | 45 | GTR+R4 | parsC |
| i75s90l50 | 0 | 2 | 42 | 3 | 47 | GTR+R4 | parsD |
| i75s90l70 | 0 | 2 | 46 | 6 | 54 | GTR+G | parsA |
| i75s90l70 | 0 | 2 | 44 | 5 | 51 | GTR+G | parsB |
| i75s90l70 | 0 | 2 | 46 | 6 | 54 | GTR+R4 | parsC |
| i75s90l70 | 0 | 2 | 45 | 6 | 53 | GTR+R4 | parsD |
| i75s90l90 | 4 | 5 | 57 | 21 | 87 | GTR+G | parsA |
| i75s90l90 | 2 | 4 | 58 | 10 | 74 | GTR+G | parsB |
| i75s90l90 | 3 | 5 | 53 | 14 | 75 | GTR+R4 | parsC |
| i75s90l90 | 4 | 5 | 55 | 16 | 80 | GTR+R4 | parsD |

**Table S7**. Fossil calibrations used in the divergence time estimation. Full information regarding the fossils selected and references are presented in the Supplementary Information.

| **MRCA** | **Tip 1** | **Tip 2** | **Minimum Age Constraint** | **Maximum Age Constraint** |
|---|---|---|---|---|
| **root** | alligator_mississippiensis | dromaius_novaehollandiae | 246.7 | 260 |
| Aves | Nettapus_auritus_GCA_011076525 | Crypturellus_cinnamomeus_OUT0029 | 66.55 | 86 |
| Stem Casuariiformes | dromaius_novaehollandiae | casuarius_casuarius | 24.459 | 58.7 |
| Galloanserae | Nettapus_auritus_GCA_011076525 | Meleagris_ocellata_FLMNH45111 | 61.1 | 86 |
| Anseres | Nettapus_auritus_GCA_011076525 | anseranas_semipalmata | 33.85 | 66.55 |
| Stem Phasianidae | Meleagris_gallopavo_NCBI_genome | Oreortyx_pictus_RTK474 | 24 | 52 |
| Mirandornithes | Podiceps_cristatus_APP039 | Phoenicopterus_ruber_APP037 | 33.1 | 46.6 |
| Columbimorphae | Pterocles_orientalis_GCA_011057875 | Columba_livia_APP015 | 23.9 | 52 |
| Otidimorphae | Tauraco_erythrolophus_APP043 | Ardeotis_arabs_GCA_011801015 | 51.81 | 66.5 |
| Sedentaves | Steatornis_caripensis_LSUMZ_169580 | Nyctibius_griseus_ANSP_183090 | 51.81 | 66.04 |
| Crown Gruiformes | grus_canadensis | rallus_limicola | 53.9 | 66.04 |
| Crown Gruidae | Balearica_regulorum_APP006 | grus_canadensis | 10.3 | 33 |
| Crown Alcini | Uria_aalge_GCA_014363425 | alca_torda_b77512 | 15.99 | 41.01 |
| Crown *Uria* | Uria_aalge_GCA_014363425 | Uria_lomvia_GCA_002289315 | 2.6 | 20.44 |
| Crown Calidridinae | Calidris_pugnax_GCF_001431845 | arenaria_interpres | 11.63 | 33.2 |
| Stem Laromorphae | larus_argentatus_b37399 | stercorarius_pomarinus_b20733 | 20.44 | 47.8 |
| Stem Jacanidae | jacana_jacana | rostratula_benghalensis | 31 | 56 |
| Stem Phaethontiformes | Phaethon_lepturus_APP035 | Eurypyga_helias_APP019 | 56 | 72.17 |
| Stem Phalacrocoracidae | anhinga_anhinga | Phalacrocorax_brasilianus_GCA_002174335 | 24 | 51.81 |
| Austrodyptornithes | Aptenodytes_forsteri_APP005 | diomedea_nigripes | 60.5 | 72.17 |
| Stem Fregatidae | fregata_magnificens | Phalacrocorax_brasilianus_GCA_002174335 | 51.81 | 66.04 |
| *Spheniscus - Eudyptula* split | eudyptula_minor | Spheniscus_magellanicus_GCA_010076225 | 9.2 | 27 |
| *Eudyptes - Megadyptes* split | Eudyptes_moseleyi_GCA_010082375 | Megadyptes_antipodes_GCA_010078485 | 3.06 | 27 |
| Crown Gaviidae | Gavia_stellata_APP022 | gavia_immer | 3.92 | 23.04 |

| | | | | |
|---|---|---|---|---|
| Stem Threskiornithidae | Pelecanus_crispus_APP034 | eudocimus_albus | 53.9 | 66.04 |
| Stem Coliiformes | colius_colius | Campephilus_principalis_AMNH363840 | 62.221 | 72.17 |
| Crown Coliidae | Colius_striatus_APP014 | urocolius_indicus | 5.05 | 23.04 |
| Crown Strigiformes | Tyto_alba_APP045 | Strix_varia_22621 | 25.3 | 61.7 |
| Crown Piciformes | Campephilus_principalis_AMNH363840 | galbula_albirostris | 30 | 62.517 |
| Stem Coracii | Coracias_garrulus_MSB30064 | Merops_ornatus_AMNHdot19747 | 53.9 | 66.04 |
| Stem Todidae | Todus_subulatus_KUNHM8104 | momotus_momota | 30 | 55 |
| Crown Falconidae | falco_mexicanus | herpetotheres_cachinnans2 | 16 | 53.5 |
| *Falco peregrinus - F. tinnunculus* divergence | Falco_peregrinus_APP020 | Falco_tinnunculus_GCA_010332995 | 5.3 | 17.5 |
| Psittacopasserae | Anodorhynchus_hyacinthinus_KU1681 | passer_domesticus_28860 | 53.9 | 66.04 |
| Crown Nestoridae | Nestor_notabilis_APP031 | strigops_habroptila_23361 | 15.9 | 27.29 |
| Stem Pelecanidae | Pelecanus_crispus_APP034 | belaeniceps_rex | 27.29 | 54.18 |
| Eupasseres | passer_domesticus_28860 | Tyrannus_tyrannus_L3328 | 27 | 56 |
| Crown Menuridae | atrichornis_rufescens_554508 | menura_novaehollandiae_76638 | 16 | 34 |
| Stem Meliphagidae | pardalotus_striatus_8886 | Meliphaga_mimikae_ANWCB52704 | 12.9 | 34 |
| *Orthonyx - Pomatostomus* split | orthonyx_temminckii_76694 | pomatostomus_superciliosus_8792 | 17.72 | 34 |
| Stem Neosittidae | mohoua_albicilla_96717 | psophodes_cristatus_6205 | 11.6 | 34 |
| Stem Cracticinae | strepera_graculina_9660 | artamus_cinereus_6183 | 14.5 | 34 |
| Stem *Ammodramus* | ammodramus_savannarum_b8557 | Arremonops_rufivirgatus_JFBM43282 | 7.5 | 18.6 |

**Table S8**. Credible clade recovery and computation time of the ASTRAL trees using the full dataset and six filtered datasets.

| **Data set** | **Filter** | **Number of non-monophyletic clades** | | | | | **Computation time (CPU hours)** | | |
|---|---|---|---|---|---|---|---|---|---|
| | | Order | Family | Genus | High-level | Sum | Gene tree | ASTRAL | Sum |
| Full dataset | | 3 | 20 | 174 | 9 | 206 | 12849 | 55814 | 68663 |
| indv_50_sites_50_loci_50 | 1 | 1 | 16 | 138 | 13 | 168 | 4226 | 19350 | 23576 |
| indv_0_sites_50_loci_70 | 2 | 2 | 17 | 175 | 13 | 207 | 4766 | 22346 | 27112 |
| indv_0_sites_90_loci_50 | 3 | 1 | 13 | 135 | 12 | 161 | 1810 | 28631 | 30442 |
| indv_50_sites_50_loci_70 | 4 | 3 | 17 | 139 | 12 | 171 | 3200 | 11622 | 14822 |
| indv_75_sites_90_loci_70 | 5 | 3 | 17 | 112 | 12 | 144 | 388 | 1452 | 1839 |
| indv_50_sites_90_loci_90 | 6 | 3 | 13 | 136 | 14 | 166 | 312 | 714 | 1026 |

**Table S9**. Summary of non-monophyletic clades for hybrid supertree and hybrid divide-and-conquer analyses. For hybrid supertree approach, the published trees were weighted based on the amount of data used to infer them, same as described for the initial supertree approach, and backbones were always given a weight of one.

| Analysis | Detail | Backbone | Trees in MRP matrix | Number of non-monophyletic clades | | | | | CPU Hour |
|---|---|---|---|---|---|---|---|---|---|
| | | | | Order | Family | Genus | High-level | Sum | |
| **Hybrid supertree** | Published trees + one fasttree (custom weight) | Full fasttree | 206 | 5 | 19 | 48 | 17 | 89 | 2165 |
| | | Filter1 fasttree | 206 | 9 | 27 | 55 | 17 | 108 | 801 |
| | | Filter2 fasttree | 206 | 6 | 21 | 50 | 11 | 88 | 641 |
| | | Filter3 fasttree | 206 | 7 | 26 | 53 | 18 | 104 | 494 |
| | | Filter4 fasttree | 206 | 6 | 19 | 43 | 12 | 80 | 500 |
| | | Filter5 fasttree | 206 | 9 | 28 | 59 | 16 | 112 | 65 |
| | | Filter6 fasttree | 206 | 8 | 26 | 54 | 19 | 107 | 50 |
| | | indv0_sites50_loci50 +GAMMA parsB | 206 | 8 | 17 | 48 | 10 | 83 | 811 |
| | | Fulldata +FreeRates parsD | 206 | 7 | 31 | 48 | 9 | 95 | 3786 |
| | Published trees + one fasttree + family backbone (custom weight) | Full fasttree + FamilyBackbone | 207 | 0 | 4 | 38 | 0 | 42 | 2164 |
| | | Filter1 fasttree + FamilyBackbone | 207 | 0 | 2 | 39 | 0 | 41 | 801 |
| | | Filter2 fasttree + FamilyBackbone | 207 | 0 | 2 | 38 | 0 | 40 | 641 |
| | | Filter3 fasttree + FamilyBackbone | 207 | 0 | 1 | 38 | 0 | 39 | 494 |
| | | Filter4 fasttree + FamilyBackbone | 207 | 0 | 2 | 39 | 0 | 41 | 500 |
| | | Filter5 fasttree + FamilyBackbone | 207 | 0 | 2 | 41 | 0 | 43 | 64 |
| | | Filter6 fasttree + FamilyBackbone | 207 | 0 | 3 | 40 | 0 | 43 | 51 |
| | | indv0_sites50_loci50 +GAMMA parsB + FamilyBackbone | 207 | 0 | 2 | 36 | 0 | 38 | 810 |
| | | **Fulldata +FreeRates parsD + FamilyBackbone (representative)*** | **207** | **0** | **2** | **35** | **0** | **37** | **3786** |
| **Hybrid divide-and-conquer** | 50 subsets + one fasttree (weight 1:1) | Full fasttree | 51 | 0 | 2 | 40 | 1 | 43 | 10122 |
| | | Filter1 fasttree | 51 | 0 | 2 | 41 | 1 | 44 | 8764 |
| | | Filter2 fasttree | 51 | 0 | 2 | 40 | 1 | 43 | 8593 |
| | | Filter3 fasttree | 51 | 1 | 3 | 39 | 1 | 44 | 8461 |
| | | Filter4 fasttree | 51 | 0 | 2 | 41 | 1 | 44 | 8467 |
| | | Filter5 fasttree | 51 | 0 | 2 | 48 | 2 | 52 | 8104 |
| | | Filter6 fasttree | 51 | 0 | 2 | 46 | 1 | 49 | 8101 |

| | | | | | | | | | |
|---|---|---|---|---|---|---|---|---|---|
| | | indv0_sites50_loci50 +GAMMA parsB | 51 | 0 | 2 | 39 | 1 | 42 | 8743 |
| | | **Fulldata +FreeRates parsD (representative)*** | **51** | **0** | **2** | **39** | **1** | **42** | **11721** |
| | 50 subsets + one fasttree + genus backbone (weight 2:1:1) | Full fasttree + GenusBackbone | 102 | 0 | 2 | 39 | 1 | 42 | 10064 |
| | | Filter1 fasttree + GenusBackbone | 102 | 0 | 2 | 38 | 1 | 41 | 8705 |
| | | Filter2 fasttree + GenusBackbone | 102 | 0 | 2 | 39 | 1 | 42 | 8545 |
| | | Filter3 fasttree + GenusBackbone | 102 | 0 | 2 | 38 | 1 | 41 | 8401 |
| | | Filter4 fasttree + GenusBackbone | 102 | 0 | 2 | 39 | 1 | 42 | 8403 |
| | | Filter5 fasttree + GenusBackbone | 102 | 0 | 2 | 39 | 1 | 42 | 7981 |
| | | Filter6 fasttree + GenusBackbone | 102 | 0 | 2 | 39 | 1 | 42 | 7961 |
| | | indv0_sites50_loci50 +GAMMA parsB + GenusBackbone | 102 | 0 | 2 | 39 | 1 | 42 | 8713 |
| | | Fulldata +FreeRates parsD + GenusBackbone | 102 | 0 | 2 | 39 | 1 | 42 | 11690 |

### Supplementary Information

#### Overlap in phylogenomic datasets

Among the 22 selected phylogenomic datasets, we observed extensive overlap in taxon sampling, i.e., multiple individuals of the same species were present in our alignments or the same sample was used in different studies. We arbitrarily selected the representative sample based on the alphabetical order of the studies (Supplementary Table S1). For example, we prioritized the “AllFam” dataset (which sampled representatives for almost all families of birds; Braun et al., 2024 bioRxiv) over Oliveros2019 dataset which sampled representatives for almost all families of passerines (Oliveros et al., 2019). Similarly, the “Harvey2020” dataset (which sampled almost all suboscine passerines; Harvey et al., 2020) also had priority over Oliveros2019 dataset.

#### Tree visualization

The RAxML-NG full tree was first collapsed to the genus level using custom perl scripts. If a genus was not monophyletic, it would appear in the tree more than once. For example, *Neopelma* was paraphyletic, therefore, it was represented by two tips, *Neopelma1* and *Neoplema2*. The genus level trees (Figures 1 and S1) were plotted in R using ape v5.7-1 (Paradis and Schliep, 2019), geiger v2.0.11 (Pennell et al., 2014), ggtree v3.10.0 (Yu et al., 2016), and RColorBrewer v1.1-3 (Neuwirth and Neuwirth, 2014). We also collapsed the fulldata parsD fasttree to the order level and plotted the trees (Figure 5a) using FigTree v1.4.4 (http://tree.bio.ed.ac.uk/software/figtree/) and Inkscape (https://inkscape.org/).

#### Novel relationships introduced in supertree analysis

The supertree method can be prone to limited taxon overlap in the source trees. For example, the three supertree analyses all failed to recover the monophyly of the *Chlamydotis* bustards (*C. macqueenii* and *C. undulata*), likely due to that all the source trees only sampled either one of the sister pair, i.e., their sister relationship was not present in any of the source trees, thus the resulting supertree also fails to support this relationship. This issue was resolved in a test run where we manually added *C. macqueenii* into the Burleigh backbone to be sister with *C. undulata*. The supertree method can also be susceptible to topological errors in the source tree. For example, all initial three supertree analyses rendered Paridae paraphyletic. *Cyanistes caeruleus* (Paridae) was only present in the backbone trees, while Jetz backbone put it together with other Paridae birds but Burleigh backbone erroneously placed it within Muscicapoidea, a different superfamily. As Elachuridae is basal in Muscicapoidea, the resulting supertree grouped *Cyanistes caeruleus* and the monotypic Elachuridae together as sisters. The paraphyly of Paridae was resolved in a test run after we pruned out *Cyanistes caeruleus* from the Burleigh backbone.

#### Comparing levels of polytomy in the divide-and-conquer trees

We found that the two divide-and-conquer trees using the genus backbone (T5 & T6) performed much better than the two trees using the family backbone (T3 & T4) in resolving expected genera (Figure 2). While the two divide-and-conquer trees T5 and T6 appeared to perform well with the exact same expected clades recovered, there were polytomies within families that were heavily sampled, such as Tyrannidae and Thamnophilidae, as well as polytomies among some of the oscine families. We evaluated the level of polytomy in T5 and T6 to assess the effect of different weighting strategies on phylogenetic accuracy. We created a completely unresolved tree for 2758 taxa, i.e., a star phylogeny that contains no topological information for any of the tips, which has the exact same taxon sampling as T5 and T6. We used ete3 (Huerta-Cepas et al., 2016) to calculate the Robinson-Foulds distance between the unresolved tree and T5 (RF = 2714), as well as between the unresolved tree and T6 (RF = 2736). T6 had a higher RF distance to the unresolved tree than T5, therefore T6 contained fewer polytomies.

**Subsets in the divide-and-conquer approach**

The first step of the divide-and-conquer approach was to subset the full supermatrix and estimate a concatenation tree for each subset supermatrix. To identify what might be a good-sized subset to run and how many computer cores to allocate for each analysis, we tested on different combinations between taxon number (the number of taxa included in a subset supermatrix: 50, 100, and 150) and core number (the number of CPUs used for each concatenation analysis: 8, 16, 24, 32). We found that the analysis running with a 150 taxon dataset and 8 CPUs had relatively higher CPU and memory efficiencies. Therefore, we built our random subsets, partially stratified subsets, and fully stratified subsets largely based on this. See Data Availability for taxon sampling in each subset.

To create the partially stratified subsets, we stratified the 2760 taxa (with *Muscipipra vetula* and *Spheniscus mendiculus* included initially) in the full data set into six major groups (Figure S1): (i) 399 taxa representing oscine passerines; (ii) 1282 taxa representing Acanthisittidae (basal in Passeriformes) and suboscines; (iii) 386 taxa representing Australaves excluding all passerines; (iv) 369 taxa representing Afroaves; (v) 162 taxa representing Neoaves excluding all Telluraves (Australaves + Afroaves); and (vi) 162 taxa representing Palaeognathae, Galloanserae and two crocodilians. These groups were built based on clades recovered across many studies, but they were not necessarily monophyletic.

For fully stratified subsets, we divided all taxa in the full data set into 25 groups based on taxonomy. We limited the number of taxa included in each subset under 200 to maximize the computing efficiency given our availability of computer resources. For example, there were four groups representing Tyrannidae (tyr1, tyr2, tyr3, and tyr4), each containing around 110 taxa, which resulted in a full coverage of our sampled taxa for Tyrannidae (414 taxa). For group tyr1, we selected 24 linker taxa from other families of suboscines, oscines, Falconiformes, and Psittaciformes (with fewer taxa selected from more taxonomically distinct groups). Group tyr1 plus the linker taxa made up one fully stratified subset. Since we wanted to ensure all taxa were included in a subset with their congeners, the fully stratified subsets had varying numbers of taxa (ranging from 99 to 191). We plotted the schematic of subsets (Figure S2) using ComplexHeatmap (Gu et al., 2016).

**List of Calibrations used in this study**
We utilized a total of 43 fossil calibrations for node-dating analyses (Supplementary Table S7). For each calibration point, we provided information for associated fossil specimens, age constraints, justification, and data source below.

**Calibrated Node:** Root (Crocodilia – Aves)
**Fossil Specimen:** *Ctenosauriscus koeneni* GZG.V.4191
**Phylogenetic Justification:** *Ctenosauriscus koeneni* has been recovered as a crown archosaur in multiple phylogenetic analyses (e.g. Butler et al. 2011). The holotype specimen selected as the calibration preserves much of the diagnostic vertebral column.
**Minimum Age Constraint:** 246.7 Ma
**Maximum Age Constraint:** 260 Ma
**Age Justification:** The holotype is from the late Olenekian Solling Formation of Germany. We therefore specified the age of the Olenekian-Anisian boundary as a minimum age, which is currently placed at 246.7 Ma based on radiometric dating combined with composite cyclostratigraphy (Gradstein et al. 2020). *Xilousuchus sapingensis* from the Heshanggou Formation of China is close in age (Nesbitt et al. 2010), but some uncertainty remains over whether the age of that taxon is late Olenkian or early Anisian. The maximum age is set to 260 Ma in order to incorporate the late Permian assemblages from which stem representatives of Archosauria such as Proterochampsidae, Euparkeriidae, Erythrosuchidae, and Proterosuchidae are known but no crown archosaurs have been uncovered, following Benton et al. (2015).

**Calibrated Node:** Crown Aves
**Fossil Specimen:** *Asteriornis maastrichtensis* NHMM 2013 008
**Phylogenetic Justification:** In phylogenetic analyses, *Asteriornis* has resolved as a crown bird with strong support (Field et al. 2020). Character states diagnosing *Asteriornis* as a member of Neornithes include edentulous upper and lower jaws and a fused mandibular symphysis, and features diagnosing it as a member of total-clade Galloanserae include a bicondylar mandibular process of the quadrate, a dorsally oriented internal articular process of the mandible, and a dorsoventrally deep and mediolaterally compressed maxillary process of the premaxilla (Field et al. 2020, Crane et al. 2024, Field et al. 2024). Although revised estimates of the age of the Lopez de Bertodano Formation make clear that fossils of *Vegavis iaai* (69.2-68.4 Ma) are slightly older than *Asteriornis*, *Vegavis* has not uniformly been inferred to be a member of Neornithes in phylogenetic analyses (e.g., Field et al. 2020). Although *Vegavis* appears to exhibit a 'neognathous' palate on the basis of a well-preserved pterygoid, recent work has suggested that an unfused pterygoid-palatine complex is plesiomorphic for crown birds (e.g., Benito et al. 2022).
**Minimum Age Constraint:** 66.55 Ma
**Maximum Age Constraint:** 86 Ma
**Age Justification:** A refined age mode for the Maastricht Formation assigns an age of 66.6 Ma (+/- 0.05 Ma) for the middle Gronsveld Member (Vellekoop et al. 2022), so we assign an age of 66.55 Ma as a hard minimum, as opposed to 66.7 Ma as originally recommended by Field et al. (2020). No substantially older crown birds are known from Cretaceous deposits, though numerous avialan-bearing fossil deposits are known from sediments between 86 MYA (the

approximate age of the avialan-rich Niobrara Formation) and the K–Pg boundary. Based on current evidence we therefore consider it unlikely that future discoveries of crown birds will predate 86 MYA.

**Calibrated Node:** Stem Casuariiformes
**Fossil Specimen:** *Emuarius gidju* QM F45460
**Phylogenetic Justification:** Worthy et al. (2014) recovered *Emuarius* as more closely related to *Dromaius* than *Casuarius* in a phylogenetic analysis. Codings for *Emuarius* were based on multiple specimens, and key synapomorphies occur in the skull, tarsometatarsus and scapulocoracoid. A scapulocoracoid (QM F45460) is thus specified as the calibrating specimen.
**Minimum Age Constraint:** 24.459 Ma
**Maximum Age Constraint:** 58.7 Ma
**Age Justification:** The calibrating fossil is from Faunal Zone A at the Hiatus South Site of the Riversleigh locality in Queensland, Australia. Based on biocorrelation to the faunas from the Etadunna and Namba Formations in South Australia (Woodbourne at al. 2014), a minimum age matching the top of Chron 7r is applied, with the numerical date selected from table 28.1 of Gradstein et al. 2020. The maximum is based on the age of the oldest putative palaeognaths, which include middle-late Paleocene lithornithids from North America and the ratite *Diogenornis*, from the early Eocene of Brazil. While the precise phylogenetic relationships of these taxa are debated, none are plausibly nested within crown Casuariiformes.

**Calibrated Node:** Crown Galloanserae
**Fossil Specimen:** *Conflicto antarcticus* MLP 07-III-1-1
**Phylogenetic Justification:** The holotype specimen of *Conflicto* is very well preserved, including a nearly-complete skull and much of the appendicular skeleton. Phylogenetic analyses by Tambussi et al. (2019) recovered *Conflicto* as a stem anseriform with strong support, resolving weakly as the sister taxon of *Nettapterornis oxfordi* from the Eocene London Clay of England. The same phylogenetic position was recovered by Field et al. 2020. Relevant synapomorphies supporting total-clade galloanseran affinities for *Conflicto* include elongate retroarticular processes, and recent work by Crane (2024) suggests a similar morphology of the internal articular process of the mandible with *Asteriornis*. Although a number of analyses (e.g., Clarke et al. 2005) have obtained a position for the marginally older fossil taxon *Vegavis iaai* among crown anseriforms, other studies have failed to consistently recover anseriform, or even neornithine affinities for *Vegavis* (e.g., Field et al. 2020). As such, we elect to use *Conflicto* as an internal calibration for crown Galloanserae.
**Minimum Age Constraint:** 61.1 Ma
**Maximum Age Constraint:** 86 Ma
**Age Justification:** The type specimen of *Conflicto antarcticus* derives from the uppermost López de Bertodano Formation (Level 10 of Montes et al. 2012a,b), which has been assigned a Danian age (~66-61 Ma). The Cretaceous-Palaeogene boundary lies just below level 10, identified on the basis of dinoflagellate biostratigraphy and Iridium traces (Tambussi et al. 2019).

**Calibrated Node:** Stem Phasianidae

**Fossil Specimen:** *Palaeortyx* cf. *gallica* PW 2005/5023a-LS
**Phylogenetic Justification:** Mayr et al. (2006) described apomorphies including the well-developed processus intermetacarpalis that support placement of *Palaeortyx* cf. *gallica* within crown Galliformes, most likely as a stem group representative of Phasianidae. This placement was confirmed by Ksepka et al. (2023). PW 2005/5023a-LS represents a nearly complete skeleton and is selected as the calibrating specimen.
**Minimum Age Constraint:** 24 Ma
**Maximum Age Constraint:** 52 Ma
**Age Justification:** The fossil is from a maar lake deposit at Enspel, near Bad Marienberg in Westerwald, Rheinland-Pfalz, Germany. These deposits are assigned to the MP28 biozone. Numerical ages ranges of the Mammal Paleogene (MP) reference levels are considered approximate (Gradstein et al. 2020), so we selected a minimum age of 24Ma for the calibration which is conservative considering MP29 and MP30 intervene between MP28 and the Paleogene-Neogene boundary at 23.04Ma. The maximum is based on the age of the Green River Formation from which multiple complete skeletons of the stem galliform *Gallinuloides wyomingensis* have been collected. This maximum encompasses other strata that have yielded good material of stem galliforms but no convincing crown galliform material including the Messel Formation and late Eocene horizons at Quercy. The maximum also encompasses the ages of taxa that may possibly represent crown galliforms but require additional study, such as *Schaubortyx.*

**Calibrated Node:** Crown Anseres
**Fossil Specimen:** *Romainvillia stehlini* NMB P.G.38.a
**Phylogenetic Justification:** *Romainvillia* provides the earliest evidence of genuinely “duck-like” forms known (Mayr 2022). Phylogenetic analyses have repeatedly supported a sister group relationship between *Romainvillia* and Anatidae (e.g., Mayr 2008), with synapomorphies for this clade including the inference of a reduced hallux (Mayr 2022). Conversely, *Romainvillia* exhibits plesiomorphies consistent with a phylogenetic position outside of crown Anatidae such as a supracoracoideus nerve foramen (Mayr 2008).
**Minimum Age Constraint:** 33.85
**Maximum Age Constraint:** 66.55
**Age Justification:** The earliest clear evidence of *Romainvillia* derives from the late Eocene of France (MP 20; Mourer-Chauviré 1996). MP20 is regarded as 33.9 +- 0.05 (Gradstein et al. 2020). The latest Cretaceous is set as the maximum, corresponding to the age range of the oldest crown birds. The oldest confirmed crown birds are latest Cretaceous in Age, and no members of Anseres are known from Cretaceous deposits, indicating it is unlikely the divergence between *Anseranas* and Anatidae had occurred before the Paleocene.

**Calibrated Node:** Crown Mirandornithes
**Fossil Specimen:** *Adelalopus hoogbutseliensis* IRScNB Av 71
**Phylogenetic Justification:** *Adelalopus hoogbutseliensis* is the oldest-known representative of Palaeolodidae, a clade of probable stem-Phoenicopteriformes exhibiting a number of phoenicopteriform synapomorphies (e.g., a deep mandibular ramus presumably associated with a thick tongue related to an incipient filter feeding apparatus; Mayr 2022), but lacking

others such as extremely long legs and a distinctively down-curved bill associated with the refinement of the specialised mode of filter feeding exhibited by extant flamingos (Mayr 2022). Character polarity among total-clade Mirandornithes is challenging to assess, given the striking morphological differences between the two extant lineages comprising the clade (flamingos and grebes). As such, the possibility that palaelodids could represent members of stem-Podicipedidae rather than stem-Phoenicopteridae cannot be excluded (Mayr 2022); however, even if this were the case, it would not affect the use of *Adelalopus* to calibrate the flamingo-grebe divergence.
**Minimum Age Constraint:** 33.1 Ma
**Maximum Age Constraint:** 46.6 Ma
**Age Justification:** The fossil bird remains from Hoogbutsel, Belgium, have been assigned an early Oligocene age (MP 21; Aguilar J.-P. et al. 1997; Mayr and Smith 2002). We note that numerical ages ranges of the Mammal Paleogene (MP) reference levels are considered approximate (Gradstein et al. 2020) but consider a minimum age of 33.1 +- 0.05 Ma to be in line with current consensus for MP21. The maximum age is based on the age of the stem-mirandornithean *Juncitarsus* from the Bridger Formation of Wyoming (Olson & Feduccia 1980; Murphey & Evanoff 2007).

**Calibrated Node**: Sedentaves
**Fossil Specimen:** *Prefica nivea* USNM 336278
**Phylogenetic Justification:** Olson (1987) discussed synapomorphies of *Prefica* and *Steatornis*, and a sister group relationship between the two was supported by the phylogenetic analysis of Mayr (2005) and subsequent work (Nesbitt et al. 2011; Ksepka et al. 2013; Chen et al. 2019).
**Minimum Age Constraint:** 51.81 Ma
**Maximum Age Constraint:** 66.04 Ma
**Age Justification:** The fossil is from Fossil Butte Member, Green River Formation, Wyoming, USA. These deposits are late early Eocene, and multicrystal analyses (sanidine) from a K-feldspar tuff (FQ-1) at the top of the middle unit of the Fossil Butte Member, from Fossil-Fowkes Basin (locality: N41º47'32.2'' W110º42'39.6") have yielded an age of 51.97 ± 0.16Ma (Smith et al. 2010). The latest Cretaceous is set as the soft maximum. The oldest confirmed crown birds are latest Cretaceous in Age, and no members of Strisores are known from Cretaceous deposits, indicating it is unlikely the highly nested divergence between oilbirds and other Strisores had occurred before the Paleocene.

**Calibrated Node:** Crown Gruiformes
**Fossil Specimen:** *Pellornis mikkelseni* MGUH 29278
**Phylogenetic Justification:** Various phylogenetic analyses have recovered Messelornithidae as sister taxon to Rallidae+Heliornithidae (Mayr, 2004; Musser et al. 2019) or to Rallidae to the exclusion of Heliornithidae (Bertelli et al. 2011). We utilize the more inclusive placement here, as minimum ages should be applied conservatively.
**Minimum Age Constraint:** 53.9 Ma
**Maximum Age Constraint:** 66.04 Ma
**Age Justification:** The fossil is from the Fur Formation of Denmark. The minimum age is based on a 54.04+/-0.14Ma radiometric date reported for layer +19 of the Fur Formation (Chambers

et al. 2003). The latest Cretaceous is set as the soft maximum. No reliable records of Gruiformes are known from Cretaceous deposits, and the maximum incorporates the possibility that Paleocene taxa such as the poorly known *Messelornis russelli* or the enigmatic *Walbeckornis* belong to crown Gruiformes.

**Calibrated Node:** Crown Gruidae
**Fossil Specimen:** *Balearica exigua* UNSM 53579
**Phylogenetic Justification**: *Balearica exigua* is known from a number of specimens that exhibit distinct similarities to extant *Balearica* across the skeleton, including the skull, beak, femur, tibiotarsus, tarsometatarsus, and humerus (Feduccia and Voorhies, 1992). Most diagnostically, *B. exigua* exhibits inflated frontals, a feature shared with extant *Balearica*, in contrast to the uninflated condition in Gruinae (the crane subfamily including all other cranes in the genera *Leucogeranus*, *Antigone*, and *Grus*). *Balearica* is the only clade of extant Neoaves exhibiting this feature (Mayr, 2018), further supporting referral of *B. exigua* to total-clade Balearicinae.
**Minimum Age Constraint**: 10.3 Ma
**Maximum Age Constraint**: 33.0 Ma
**Age Justification**: The specimens derive from the upper Clarendonian Ash Hollow Formation of the Cap Rock Member, near Orchard, Nebraska (Feduccia and Voorhies, 1992). Although the specimens derive from a 2m-thick volcanic ash bed, and despite previous work on the age of the Ash Hollow Formation (Boellstorf, 1978), precise constraints on the age of this locality are lacking. Considering this uncertainty, and given the upper Clarendonian age of this locality, we assign an age of 10.3 Ma, corresponding to the minimum age of the Clarendonian inclusive of error. The maximum age encompasses the early Oligocene, a time interval in which potential close relatives of Gruideae are known such as the putative limpkin *Badistornis aramus* and *Parvigrus pohli*, which may represent the sister taxon of Aramidae + Gruidae (Mayr, 2005; Musser et al. 2019) or of Gruoidea (Mayr, 2013).

**Calibrated Node:** Crown Otidimorphae
**Fossil Specimen:** *Foro panarium* USNM 336261
**Phylogenetic Justification:** *F. panarium* was supported as the sister taxon to crown Musophagidae on the basis of phylogenetic analyses employing multiple alternative backbone constraints (Field & Hsiang 2018). Character states resolving as unambiguous synapomorphies of an exclusive Musophagidae + *Foro* clade include a furcula unfused at its midline, large tubercula praeacetabularia of the pelvis, os carpi ulnare with crus longum greatly abbreviated, and bill short and stout with broad processus maxillaris of os nasale (Olson 1992). A number of presumed plesiomorphic features unobserved in crown musophagids (e.g., elongate hindlimbs) suggest a phylogenetic placement for *Foro* on the stem of Musophagidae (Field & Hsiang 2018).
**Minimum Age Constraint:** 51.81 Ma
**Maximum Age Constraint:** 66.5 Ma
**Age Justification:** The fossil is from the Fossil Butte Member, Green River Formation, Wyoming, USA. These deposits are late early Eocene in age, and multicrystal analyses (sanidine) from a K-feldspar tuff (FQ-1) at the top of the middle unit of the Fossil Butte Member, from the Fossil–Fowkes Basin have yielded an age of 51.97 +/- 0.16 Ma (Smith et al. 2010). The latest Cretaceous is set as the maximum, corresponding to the age range of the oldest crown birds.

*Foro panarium* is easily the oldest known well supported member of Otidimorphae, indicating that although an Otidimorphae ghost lineage must extend earlier into the Paleogene, a Cretaceous divergence among crown Otidimorphae is unlikely.

**Calibrated Node:** Crown Columbimorphae
**Fossil Specimen:** *Leptoganga sepultus* MNHN Av-2844
**Phylogenetic Justification:** Several taxa referred to stem-Pteroclidae are known from the Quercy fissure fillings, but *Leptoganga* is arguably the most diagnostic of total-clade Pteroclidae in possessing an apomorphic intermetatarsal sesamoid (Mourer-Chauviré 1993). Additional aspects of the osteology of these taxa are closely comparable with extant *Pterocles* (Mayr 2022). Since *Leptoganga* is inferred to represent a stem-group member of Pteroclidae, it serves as an internal calibration for the divergence between Pteroclidae and their extant sister taxon, which in some topologies is found to be Columbidae as long assumed based on morphological similarities, but in other recent topologies could be the rest of Columbimorphae (Columbidae + Mesitornithidae).
**Minimum Age Constraint:** 23.9 Ma
**Maximum Age Constraint:** 52.0 Ma
**Age Justification:** *Leptoganga sepultus* derives from the Quercy fissue fillings, assigned to the MP28 biozone. Numerical ages ranges of the Mammal Paleogene (MP) reference levels are considered approximate (Gradstein et al. 2020), so we selected a minimum age of 24 Ma for the calibration which is conservative considering MP29 and MP30 intervene between MP28 and the Paleogene-Neogene boundary at 23.04 Ma. In addition to being less diagnostic morphologically, other putative stem-pteroclids from Quercy are less well constrained stratigraphically, bolstering the case for selecting *Leptoganga* to calibrate this node. The maximum is based on the age of the Green River Formation, a diverse fossil avifauna from which no fossil members of Columbimorphae have been recovered.

**Calibrated Node:** Crown Alcini
**Fossil Specimen:** *Miocepphus bohaski* USNM 237142
**Phylogenetic Justification**: *Miocepphus bohaski* was recovered as more closely related to *Alle* than any other extant taxon of Alcidae in combined analyses of morphological and molecular data (Smith, 2011a; Smith and Clarke, 2011). Because *Alle* is not sampled in our dataset, we applied this calibration to the next rootward node in the tree.
**Minimum Age Constraint**: 15.99 Ma
**Maximum Age Constraint**: 41.01 Ma
**Age Justification**: The calibrating fossil was recovered from the Popes Creek Sand Member of the Calvert Formation. The age of the Popes Creek Sand Member was considered to be close to the Burdigalian-Aquitanian boundary by Wijnker and Olson (2009). In the absence of radiometric dates, Smith (2015) considered the minimum age of the Burdigalian stage to be a conservative minimum age for calibrations based on this fossil. *Miocepphus bohaski* is currently the oldest vetted record of crown Alcidae. A putatively Eocene (~35 Ma) humerus from Georgia was assigned to Alcidae by Chandler and Parmley (2003), but it is uncertain whether this fossil represents a stem or crown member of Alcidae (Smith and Clarke, 2011). However, because this

specimen was badly worn, was collected from a mine, and is nearly twice as old as the next earliest fossil record of Alcidae, we consider it possible it comes from younger strata that were exposed during mining and deem it unreliable as a minimum age constraint. The maximum age constraint extends to the base of the Bartonian, encompassing putative age of the Georgia humerus as well as the stage below it.

**Calibrated Node:** Crown *Uria*
**Fossil Specimen:** *Uria lomvia* CASG 71892
**Phylogenetic Justification**: Olson (2013) referred this partial postcranial skeleton to the extant species *Uria lomvia* based upon apomorphies of the coracoid and humerus. Smith (2015) advocated for use of this fossil as a calibration, noting it is identical to modern *Uria lomvia* for all phylogenetically informative characters considered in recent phylogenetic analyses of Alcidae.
**Minimum Age Constraint**: 2.6 Ma
**Maximum Age Constraint**: 20.44 Ma
**Age Justification**: Transgressive marine sequences exposed at the fossil locality Tolstoi Point, are interpreted as equivalent to the Bigbendian and Colvillian stages, which have ages of 2.6-3.0 Ma (Brigham-Grette and Carter 1992; Smith, 2015). The maximum age is based on the Burdigalian-Aquitanian, in order to encompass the age range of the oldest fossil specimens of Alcini (*Miocepphus bohaski*).

**Calibrated Node:** Crown Calidridinae
**Fossil Specimen:** *Mirolia brevirostrata* BS 1970 XVIII
**Phylogenetic Justification:** Ballmann (2004) assigned *Mirolia brevirostrate* to Calidridinae based on synapomorphies of the holotype skull (BS 1970 XVIII) and referred postcranial specimens including the short bill and processus retroarticularis and the shape of the processus supracondylaris dorsalis on the humerus.
**Minimum Age Constraint:** 11.63 Ma
**Maximum Age Constraint:** 33.2 Ma
**Age Justification:** The fossil comes from the Nördlinger Ries basin, which has been dated to the Langhian-Serravallian on the basis of faunal correlation (Ballmann, 2004). In the absence of numerical dates, Smith (2015) recommended using the age of the Serravallian-Tortonian boundary as a minimum age constraint for calibrations based on *Mirolia brevirostrata*. The maximum constraint is set to the Eocene-Oligocene boundary, based on the oldest secure records of Scolopaci, which are early Oligocene jacanids from the Jebel Qatrani Formation of Egypt (Rasmussen et al. 1987)

**Calibrated Node:** Stem Laromorphae
**Fossil Specimen:** *Laricola elegans* NMB s.g.18810
**Phylogenetic Justification:** De Pietri et al (2011) recovered *Laricola* as either the sister to Laridae (=Laromorphae) or within Laridae (with *Anous* the sister taxon to all other Laridae). Smith (2015) recommended *Laricola* as a crown Laromorphae calibration, however, the analysis upon which this was based was conducted before new cranial material was described. We conservatively place it as sister to Laromorphae, reflecting this uncertainty.

**Minimum Age Constraint:** 20.44 Ma
**Maximum Age Constraint:** 47.8 Ma
**Age Justification:** The fossil is from Saint-Gérand-le-Puy, France. Quarries at Saint-Gérand-le-Puy span the Oligocene and Miocene, but De Pietri et al (2011) were unable to confirm or refute whether any of the historically collected *Laricola* material comes from the Oligocene age deposits. We thus conservatively select one of the Miocene skulls as the calibrating fossil and apply the upper bound of the Aquitanian for the hard minimum, given the phylogenetic placement of the *Laricola* is supported primarily by cranial characters. The oldest reasonably complete fossil assignable to Charadriiformes is an unnamed Eocene (Lutetian) fossil SMF-ME 2458A+B (Mayr, 2000). The lower bound of the Lutetian is thus used as a maximum.

**Calibrated Node:** Stem Jacanidae
**Fossil Specimen:** *Nupharanassa tolutaria* DPC 2580
**Phylogenetic Justification:** This specimen is represented only by a partial tarsometatarsus. However, this bone bears distinctive apomorphies of Jacanidae including an extremely enlarged distal vascular foramen, broad tendinal groove, and strongly flattened shaft (Rasmussen et al. 1987).
**Minimum Age Constraint:** 31 Ma
**Maximum Age Constraint:** 56 Ma
**Age Justification:** The calibrating fossil was collected from Quarry E of the Jebel Qatrani Formation (Rasmussen et al. 1987). Based on paleomagnetic correlation, the age of Quarry E is estimated to be 31.0-33.2 Ma (Seiffert, 2006). The maximum age is set to the earliest Eocene based on the oldest plausible records of Charadriiformes, which include the potential stem group Charadriiformes *Scandiavis* and *Nahmavis* as well as distal humeri from Virginia and Mongolia that preserve the characteristic shape of the dorsal supracondylar process (Bertelli et al. 2013; Mayr, 2016; Hood et al. 2019; Musser et al. 2020).

**Calibrated Node:** Stem Phaethontiformes
**Fossil Specimen:** *Lithoptila abdounensis* OCP.DEK/GE 1087
**Phylogenetic Justification:** Phylogenetic analyses by Bourdon et al. (2005) and Smith (2010) recover *Lithoptila abdounensis* as a stem representative of Phaethontiformes, and cranial characters preserved in OCP.DEK/GE 1087 support this placement. Although the position of Phaethontidae within Aves is controversial, there is little doubt regarding the placement of *Lithoptila*, which tracks Phaethontidae regardless of the arrangement of other taxa.
**Minimum Age Constraint:** 56 Ma
**Maximum Age Constraint:** 72.17 Ma
**Age Justification:** The fossil was collected from an unspecified quarry, assigned to Bed IIa of the Ouled Abdoun Basin, near Grand Daoui, Morocco, which in turn can be assigned to the Thanetian based on selachians identified in the matrix (Bourdon et al. 2005). As both the precise numerical age of Bed IIa deposits and the precise horizon from which the fossil was collected remain uncertain, the lower age bound for the Thanetian is used as a hard minimum. More fragmentary records of probable Phaethontiformes are known from slightly older (Danian) deposits in New Zealand (Mayr and Scofield 2016). We conservatively rely on *Lithoptila*, but note that these records are encompassed between the minimum and maximum

bounds. The maximum age extends to the base of the Maastrichtian to accommodate the possibility that some of the poorly represented marine birds from the Cretaceous-Paleogene of New Jersey potentially represent tropicbirds (Bourdon et al. 2008).

**Calibrated Node:** Stem Phalacrocoracidae
**Fossil Specimen:** *Oligocorax* (=*Borvocarbo*) *stoeffelensis* PW 2005/5022-LS
**Phylogenetic Justification:** Phylogenetic analysis by Smith (S42) and Mayr (S45) recover *Oligocorax stoeffelensis* as more closely related to Phalacrocorax than to Anhinga. PW 2005/5022-LS preserves a substantial portion of the skeleton, including synapomorphy-bearing elements.
**Minimum Age Constraint:** 24 Ma
**Maximum Age Constraint:** 51.81 Ma
**Age Justification:** The fossil is from a maar lake deposit at Enspel in Germany. These deposits are assigned to the MP28 biozone. Numerical ages ranges of the Mammal Paleogene (MP) reference levels are considered approximate (Gradstein et al. 2020), so we selected a minimum age of 24Ma for the calibration which is conservative considering MP29 and MP30 intervene between MP28 and the Paleogene-Neogene boundary at 23.04Ma. Comparable in age is the Late Oligocene Nambashag from the Australian Etadunna and Namba Formations (S46), which also represents a stem member of Phalacrocoracidae (S45). The maximum is based on the age of the Green River Formation, from which members of Aequornithes such as Limnofregata and Vadaravis have been recovered.

**Calibrated Node:** Austrodyptornithes
**Fossil Specimen:** *Waimanu maneringi* CM zfa35
**Phylogenetic Justification:** Phylogenetic analysis supports the placement of *Waimanu* along the stem penguin lineage (Slack et al. 2006; Ksepka et al. 2012). CM zfa35 is the only published specimen of *Waimanu manneringi.*
**Minimum Age Constraint:** 60.5 Ma
**Maximum Age Constraint:** 72.17 Ma
**Age Justification:** Biostratigraphic evidence, specifically the ranges of *Hornibrookina teuriensis* and *Chaismolithus bidens* indicate the minimum possible age of the type locality is 60.5 Ma (Hornibrook et al. 1989; Slack et al. 2006; Cooper et al. 2004). The maximum is based on the lower bound of the Maastrichtian Stage. Penguin fossils from the Paleocene Chatham Islands are potentially close in age to *Waimanu* but less tightly constrained, and based on morphology appear to represent more crownward (and thus potentially younger) lineages. Southern Hemisphere Maastrichtian marine vertebrate sites have yielded diving birds such as *Polarornis* and hesperornithids, indicating preservation potential for marine diving birds, but no penguin (or procellariiform) remains have been recovered at these sites.

**Calibrated Node:** Stem Fregatidae
**Fossil Specimen:** *Limnofregata azygosternon* USNM 22753
**Phylogenetic Justification:** Phylogenetic analysis supports the placement of *Limnofregata* as the sister taxon to extant *Fregata* (Smith et al. 2010), in agreement with longstanding interpretations of this fossil taxon (Olson, 1977). USNM 22753 is an articulated skeleton

preserving most key synapomorphies that place *Limnofregata azygosternon* on the frigatebird stem lineage.
**Minimum Age Constraint:** 51.81 Ma
**Maximum Age Constraint:** 66.04 Ma
**Age Justification:** The fossil is from the Fossil Butte Member, Green River Formation, Wyoming, USA. These deposits are late early Eocene in age, and multicrystal analyses (sanidine) from a K-feldspar tuff (FQ-1) at the top of the middle unit of the Fossil Butte Member, from Fossil–Fowkes Basin have yielded an age of 51.97 +/- 0.16 Ma (Smith et al. 2010). A few fragmentary records of *Limnofregata* are known from slightly older (~2Ma) deposits of the Wasatch Formation (Stidham, 2015 ) and Namejoy Formation (Mayr, 2016). We conservatively rely on the complete Fossil Butte skeleton, but note that these records are encompassed between the minimum and maximum bounds. The latest Cretaceous is set as the soft maximum. No well-supported material from the core waterbird clade Aequornithes are known from Cretaceous deposits, indicating it is unlikely the highly nested divergence between Fregatidae and Sulidae had occurred before the Paleocene.

**Calibrated Node:** *Spheniscus* – *Eudypula* split
**Fossil Specimen:** *Spheniscus muizoni* MNHN PPI 147
**Phylogenetic Justification:** Synapomorphies listed by Gölich ( 2007) support this fossil taxon as a member of the genus *Spheniscus*, and this placement was supported by several subsequent phylogenetic analyses (e.g. Ksepka et al. 2012)
**Minimum Age Constraint:** 9.2 Ma
**Maximum Age Constraint:** 27 Ma
**Age Justification:** In the original description (Gölich, 2007) an age estimate of 11-13 Ma was provided for this fossil. However, subsequent work (Brand et al. 2011) shows the holotype horizon to be younger in age. The maximum extends into the Late Oligocene, encompassing well-described fossil penguin faunas from the Late Oligocene-Early Miocene of New Zealand and South America which have yielded many articulated and associate skeletons of multiple species of stem lineage penguins but no reliable records of crown penguins.

**Calibrated Node:** *Eudyptes* – *Megadyptes* split
**Fossil Specimen:** *Eudyptes atatu* NMNZ S.046318
**Phylogenetic Justification:** This specimen is the holotype of *Eudyptes atatu*, a fossil taxon recovered a stem representative of the crested penguin genus *Eudyptes* by Thomas et al. (2020). Assignment of this species to *Eudyptes* is supported by derived features including strong sigmoid curvature of jugal bar, presence of shelf of bone bounding the salt gland fossa, greatly deepened temporal fossae, strongly shortened tarsometatarsus (ratio of length to proximal width less than 2.0) and a moderately deep sulcus between metatarsals II and III. The less deepened mandible supports placement outside of the clade formed by all extant *Eudyptes* species.
**Minimum Age Constraint:** 3.06 Ma
**Maximum Age Constraint:** 27.0 Ma
**Age Justification:** The minimum age is based on that for the Late Pliocene Tangahoe Formation, Taranaki, New Zealand (Naish et al. 2005). The Tangahoe Formation has been tightly

constrained between 3.36 and 3.06 Ma using magnetostratigraphic correlation to the d18O timescale from Ocean Drilling Program Site 846, and the presence of Waipipian stage macro- and microfossils (Naish et al. 2005). Because many species of the *Eudyptes* + *Megadyptes* clade occur on islands and have no pre-Holocene fossil records, we used a conservative maximum. The maximum extends into the Late Oligocene, encompassing well-described fossil penguin faunas from the Late Oligocene-Early Miocene of New Zealand and South America which have yielded many articulated and associate skeletons of multiple species of stem lineage penguins but no reliable records of crown penguins.

**Calibrated Node:** Crown Gaviidae
**Fossil Specimen:** *Gavia howardae* USNM 192845
**Phylogenetic Justification:** *Gavia howardae* is known from a large number of isolated bones, which closely resemble *Gavia stellata* in their slender proportions. A coracoid was selected as the calibrating fossil because this element bears a synapomorphy shared with *Gavia stellata*: the coracoid bears a medial notch for the foramen nervi supracoracoidei, whereas all other extant loon species have a completely enclosed foramen (Olson and Rasmussen, 2001).
**Minimum Age Constraint:** 3.92 Ma
**Maximum Age Constraint:** 23.04 Ma
**Age Justification:** The oldest *Gavia howardae* come from the Yorktown Formation of the Lee Creek Mine in North Carolina (Olson and Rasmussen, 2001). Because fossils at Lee Creek Mine are commonly collected from spoil piles, the precise stratigraphic horizon of individual *Gavia howardae* elements remains uncertain. Although they are most likely to be derived from the Sunken Meadow Member (Emerson, 2008), none of the fossils were conclusively placed to that horizon and so we conservatively use the age of the top of the Yorktown Formation as a minimum age. Boessenecker et al. (2018) argued for an age range of 4.9–3.92 Ma for the Yorktown based on strontium isotope (Browning et al. 2009) and calcareous nannoplankton ranges. A number of Miocene fossils, most of them highly incomplete, have been assigned to *Gavia*, along with several stem loon taxa such as *Petralca* and *Colymboides* (Olson, 1985; Göhlich & Mayr, 2018). The maximum age was set to the base of the Miocene to accommodate the possibility that at least some of the material identified to *Gavia* may truly represent crown loons.

**Calibrated Node:** Stem Threskiornithidae
**Fossil Specimen:** *Rhynchaeites* sp. MGUH 20288
**Phylogenetic Justification:** Multiple apomorphies support the placement of *Rhynchaeites* within the total clade Threskiornithidae (Mayr and Bertelli, 2011). Although the characteristic ibis-type bill is not preserved in MGUH 20288, derived characteristics of the hindlimb support assignment to *Rhynchaeites* as well as placement along the stem lineage of Threskiornithidae for this specimen (Mayr and Bertelli, 2011).
**Minimum Age Constraint:** 53.9 Ma
**Maximum Age Constraint:** 66.04 Ma
**Age Justification:** The minimum age is based on a 54.04+/-0.14Ma radiometric date reported for layer +19 of the Fur Formation (Chambers et al. 2003). The latest Cretaceous is set as the soft maximum. No well-established members of the core waterbird clade Aequornithes are

known from Cretaceous deposits, indicating it is unlikely the highly nested divergence between ibises and other waterbirds occurred before the Paleocene.

**Calibrated Node**: Stem Coliiformes
**Fossil Specimen**: *Tsidiiyazhi abini* NMMNH P-54128
**Phylogenetic Justification**: Combined analyses by Ksepka et al. (2017) recovered *Tsidiiyazhi abini* as a stem mousebird, regardless of whether only morphological data are considered or the relationships of extant taxa are constrained using various topologies recovered by recent large-scale molecular studies.
**Minimum Age Constraint**: 62.221 Ma
**Maximum Age Constraint**: 72.17 Ma
**Age Justification**: The fossil was collected from the Ojo Encino Member of the Nacimiento Formation. This horizon falls within magnetochron C27N, constraining the absolute geochronological age to 62.221–62.517 Ma. No fossils older than the latest Cretaceous have yet been firmly identified as crown birds, and no potential mousebirds are represented in the scrappy but increasingly diverse Maastrichtian avifaunas of North America. Since mousebirds are small-bodied and thus potentially less likely to enter the fossil record than larger birds, we extended the maximum age to the base of the Maastrichtian.

**Calibrated Node:** Crown Coliidae
**Fossil Specimen:** *Colius hendeyi* SAM-PQ-L28858
**Phylogenetic Justification:** *Colius hendeyi* is known from a large number of isolated bones (Rich and Haarhoff, 1985). We select the holotype tarsometatarsus as the calibrating specimen. Referred material shows this species shares several features with *Colius* that are absent in *Urocolius* including humerus exceeding ulna in length, which is likely derived within *Colius* based on the wing proportions of stem mousebirds such as *Oligocolius* and *Palaeospiza*. The holotype tarsometatarsus suggests the fossil may be nested within crown *Colius*, as it shares one character only with *Colius leucocephalus* but not other extant *Colius* species: trochlea metartarsi III asymmetrical in distal view, with lateral rim protruding farther caudally (Ksepka, pers. obs.). As *Colius leucocephalus* is not sampled in this study, we apply the calibration to the *Colius* – *Urocolius* split.
**Minimum Age Constraint:** 5.05 Ma
**Maximum Age Constraint:** 23.04 Ma
**Age Justification:** The calibrating fossil was collected from the Varswater Formation at E Quarry, Langebaanweg, South Africa. Global sea-level reconstructions indicate an age of 5.15+/-0.1Ma for this horizon (Roberts et al. 2011). The geographical distribution of mousebirds makes formulating a maximum difficult. Stem mousebirds have a long and dense Northern Hemisphere record extending all the way to the early Paleocene, but crown group mousebirds are restricted to Africa where fossil sampling is poor. The earliest record of mousebirds in Africa is from the late Miocene, but this fragmentary tarsometatarsus cannot be assigned to either of the extant genera and the possibility it thus represents a stem taxon cannot be ruled out (Mourer-Chauviré, 2008). Given that crown mousebirds may have originated in Africa, where collecting effort for fossil birds has been limited, we used the base of the Miocene as a maximum bound.

**Calibrated Node:** Crown Strigiformes
**Fossil Specimen:** *Heterostrix tatsinensis* PIN 3211/35
**Phylogenetic Justification:** *Heterostrix* was originally considered a possible stem owl (Kurochkin and Dyke, 2011). However, Mayr (2022) noted that the holotype specimen has a fully ossified arch for the extensor tendons of the toes, which is absent in Tytonidae and stem owls and considered a synapomorphy of Strigidae.
**Minimum Age Constraint:** 25.3 Ma
**Maximum Age Constraint:** 61.7 Ma
**Age Justification:** Magnetostratigraphy and radiometric dates provide an age of 25.3Ma (Höck et al. 1999; Sun and Windley, 2015) for the boundary between the Hsanda Gol Formation and the overlying Loh Formation. Because the precise horizon from which PIN 3211/35 was collected was not reported, we use this date as a minimum age constraint. The Late Eocene *Aurorornis* resembles *Heterostrix* and may potentially represent an older record of Strigidae (Mayr, 2022). However, because the ossified loop is incomplete in this taxon and we were unable to evaluate the specimen directly, we conservatively retain *Heterostrix* as the reference fossil for the minimum age. The maximum age is based on the oldest reported stem owl, *Ogygoptynx wetmorei*. This taxon is known from the Tiffanian NALMA which overlaps the Thanetian-Selandian, so we specified the upper age bound of the Selandian as the maximum age constraint.

**Calibrated Node:** Crown Piciformes
**Fossil Specimen:** *Rupelramphastoides knopfi* SMF Av 500
**Phylogenetic Justification:** Mayr (2005, 2006) provided evidence from synapomorphies of the tarsometatarsus and ulna that clearly support placement of this fossil within total clade Pici. However, uncertainty remains over whether this taxon belongs within the crown Pici or is outside this clade. Conservatively, it is used as a calibration for the Pici-Galbulae split.
**Minimum Age Constraint** 30 Ma
**Maximum Age Constraint:** 62.517 Ma
**Age Justification:** The fossil is from Frauenweiler, Germany. The Frauenweiler locality is considered to be MP22 (Micklich and Hildebrandt, 2005). Numerical ages ranges of the Mammal Paleogene (MP) reference levels are considered approximate (Gradstein et al. 2020), so we selected a minimum age of 30Ma for the calibration which is generally considered younger than the MP22-MP23 boundary. The maximum is based on the oldest described member of Afroaves, the stem mousebird *Tsidiiyazhi abini*. This results in a very wide age range, but in doing so extends the bounds to include a number of enigmatic Eocene semi-zygodactyl and zygodactyl birds that may could potentially represent Piciformes such as the Sylphornithidae, Gracilitarsidae, and *Eutreptodactylus*.

**Calibrated Node:** Stem Coracii
**Fossil Specimen:** *Septencoracias morsensis* MGUH.VP 9509
**Phylogenetic Justification:** Phylogenetic analyses place *Septencoracias* along the stem lineage leading to the clade Coracioidea (rollers and ground rollers) (Bourdon et al. 2016; Mayr et al. 2022). Interestingly, some derived features are shared between Meropidae (bee-eaters) and

*Septencoracias* but not other Coracioidea. Here we accept the phylogenetic evidence for a close relationship between *Septencoracias* and rollers, but we note that if new evidence later places *Septencoracias* closer to bee-eaters the calibrated node would remain the same.
**Minimum Age Constraint:** 53.9 Ma
**Maximum Age Constraint:** 66.04 Ma
**Age Justification:** The fossil is from the Fur Formation of Denmark. The minimum age is based on a 54.04+/-0.14Ma radiometric date reported for layer +19 of the Fur Formation (Chambers et al. 2003). A specimen of *Septencoracias* was also described from the contemporaneous London Clay Formation (Mayr et al. 2022), and stem Coracii are abundant in the slightly younger Green River Formation and Messel Formation. The latest Cretaceous is set as the soft maximum. No members of the "landbird" clade Telluraves are known from Cretaceous deposits, indicating it is unlikely the highly nested Coracioidea – Meropidae divergence had occurred before the Paleocene.

**Calibrated Node:** Stem Todidae
**Fossil Specimen:** *Palaeotodus itardiensis* SMF Av505
**Phylogenetic Justification:** Mayr and Knopf (2007) identified derived characters of Todidae including the scapi claviculárum of the furcula being very thin, the proximal end of the humerus reaching far ventrally and being inflected so that almost the entire caput humeri is situated farther ventrally than the ventral margin of the shaft, a carpometacarpus with a large processus intermetacarpalis, a greatly elongated and slender tarsometatarsus measuring almost the length of the humerus, and the plantar surface of trochlea metatarsi III bearing a marked sulcus.
**Minimum Age Constraint:** 30 Ma
**Maximum Age Constraint:** 55 Ma
**Age Justification:** The fossil is from Frauenweiler south of Wiesloch (Baden-Württemberg, Germany), former clay pit of the Bott-Eder GmbH (“Grube Unterfeld”). The Frauenweiler locality is considered to date to MP22 (32Ma) (Micklich and Hildebrandt, 2005). Numerical age ranges of the Mammal Paleogene (MP) reference levels are considered approximate (Gradstein et al. 2020), so we selected a minimum age of 30Ma for the calibration which is generally considered younger than the MP22-MP23 boundary. The oldest reported Coraciiformes (*sensu* Yuri et al. 2015) are from the early Eocene. Given this limit and the absence of Todidae in Lagerstätten such as the Green River, Messel, London Clay, and Fur Formations which otherwise preserve an abundance of small birds, a maximum of 55Ma is specified.

**Calibrated Node:** Crown Falconidae
**Fossil Specimen:** *Pediohierax ramenta* USNM 13898
**Phylogenetic Justification:** Phylogenetic analysis (Li et al. 2014) supports placement of *Pediohierax ramenta* as a crown member of Falconidae. All remains of this taxon are isolated bones, and the apomorphies supporting placement as sister to *Falco* occur in the humerus and tarsometatarsus. Therefore, a tarsometatarsus is chosen as the calibrating specimen.
**Minimum Age Constraint:** 16 Ma
**Maximum Age Constraint:** 53.5 Ma

**Age Justification:** The fossil is from the Merychippus Quarry, Sand Canyon Member of the Sheep Creek Formation, Nebraska. The Sheep Creek Formation is assigned to the Hemmingfordian North American Land Mammal Age. Thus, the end of the Hemmingfordian is used as a minimum age for the calibration. There are many "raptorial" birds of uncertain affinities in the fossil record, which potentially represent Falconiformes, Accipitriformes, or some separate clade. The maximum extends back to the Eocene to include the taxon *Masillaraptor* which was originally described from the Messel Formation and recently reported from the London Clay. This taxon is the oldest well-represented potential representative of Falconidae though its placement is far from resolved as it shares derived traits with many raptorial clades (Mayr, 2009).

**Calibrated Node:** Divergence between *Falco tinnunculus* and *Falco peregrinus*
**Fossil Specimen:** *Falco hezhengensis* IVPP V14586.
**Phylogenetic Justification:** Phylogenetic analysis (Li et al. 2014) supports placement of *Falco hezhengensis* as the sister taxon to *Falco tinnunculus* and *Falco rupicoloides*. This species is known from a nearly complete skeleton that shares two synapomorphies with extant kestrels: weak proximal projection of the trochanteric crest of the femur and a very deep infracotylar fossa
in the tarsometatarsus (Li et al. 2014).
**Minimum Age Constraint:** 5.3Ma
**Maximum Age Constraint:** 17.5Ma
**Age Justification:** The fossil is from Liushu Formation, and the lithology suggests it was derived from the *Hipparion* fauna beds (Li et al. 2014). Because the precise stratigraphic horizon from within the Liushu Formation remains uncertain, we conservatively used a date of 5.3Ma for the minimum age, corresponding to the age given for the upper bound of this unit by Deng et al. (2013). The maximum age is based on that of *Thegornis musculosus*, the oldest substantiated crown member of Falconidae which is dated to the Santacrucian South American Land Mammal Age, providing a maximum age 17.5Ma.

**Calibrated Node:** Crown Psittacopasserae
**Fossil Specimen:** *Primoscens minutus* NHMUK A4681
**Phylogenetic Justification:** *Primoscens* is the first described species of the Zygodactylidae (Mayr, 2022). Multiple phylogenetic analyses have recovered Zygodactylidae as sister taxon to crown Passeriformes (e.g. Mayr, 2008).
**Minimum Age Constraint:** 53.9 Ma
**Maximum Age Constraint:** 66.04 Ma
**Age Justification:** The fossil is from the Walton Member (Division A2) of the London Clay Formation at Walton-on-the-Naze, England. Although NHMUK A4681 is highly incomplete (single carpometacarpus), multiple more complete records in private collections confirm the presence of Zygodactylidae at this locality (Mayr 2022). The Walton Member correlates to the upper part of Chron C24r, and the minimum age is based on the revised estimate for the top of this Chron (Gradstein et al. 2020: Table 28.1). The latest Cretaceous is set as the soft maximum. No members of the "landbird" clade Telluraves (to which Psittacopasserae belong) are known

from Cretaceous deposits, indicating it is extremely unlikely the highly nested parrot-songbird divergence had occurred before the Paleocene.

**Calibrated Node:** Crown Nestoridae
**Fossil Specimen:** *Nelepsittacus minimus* NMNZ S.52404
**Phylogenetic Justification:** Worthy et al. (2011) reported several apomorphies that support a placement for *Nelepsittacus* closer to *Nestor* than to *Strigops.* A unique apomorphy, the foramen vasculare distale being bounded on its dorsal facies by a ridge extending proximal of it, creating a shallow groove extending proximal of the foramen, is observed in NMNZ S.52404.
**Minimum Age Constraint:** 15.9 Ma
**Maximum Age Constraint:** 27.29 Ma
**Age Justification:** The fossil is from Bed HH2b, Manuherikia River section, located 21.02–21.31 m above the base of the Bannockburn Formation (Worthy et al. 2011). The Bannockburn Formation is considered to be Altonian in age (Worthy et al. 2011). The numerical age is thus based on the upper boundary of the Altonian Stage. Given the sparse record of fossil parrots in general and also of non-aquatic birds in New Zealand, it is difficult to formulate a maximum. The oldest potential crown parrot fossils globally are Late Oligocene or Early Miocene in age (Mayr, 2010), thus we specify the upper bound of the Late Oligocene as a maximum here.

**Calibrated Node:** Stem Pelecanidae
**Fossil Specimen:** *Pelecanus* sp. NT-LBR-039
**Phylogenetic Justification:** Louchart et al. (2011) noted this skull is essentially identical to that of extant *Pelecanus*, and particularly shows features that do not occur in combination in any other modern birds such as the spatulate bill with ventral ridges, exceptionally long and thin mandible, and presence of a syndesmotic intraramal hinge.
**Minimum Age Constraint:** 27.29 Ma
**Maximum Age Constraint:** 54.18 Ma
**Age Justification:** The calibrating fossil is a Rupelian limestone in Luberon, France. Because no finer stratigraphic data was noted for the fossil, we conservatively use the upper bound of the Rupelian as a minimum age constraint. The late Eocene *Eopelecanus* predates the Luberon skull but is known only from a tibiotarsus (El Adli et al. 2021). We consider the skull to be a substantially more secure record, and so use its age for the minimum age constraint while also ensuring the maximum constraint accommodates the possibility *Eopelecanus* is a stem pelican. The putative late Eocene pelican *Protopelicanus* is known only from a femur, but its assignment to Pelecanidae is considered erroneous (Olson, 1985). The oldest putative fossil records of Balaenicipitidae are an ulna and incomplete tarsometatarsus from the Jebel Qatrani Formation of Egypt, the former of uncertain age and the latter from the early Oligocene (Rasmussen et al., 1987). The maximum age is based on the oldest secure record of Pelecaniformes, the early Eocene stem ibis *Rhynchaeites*.

**Calibrated Node:** Eupasseres
**Fossil Specimen:** Eupasseres indet. SMF Av 504
**Phylogenetic Justification:** The fossil includes a partial wing. The presence of a distally protruding fingerlike process at the cranial edge of metacarpal III is an apomorphy supporting

assignment of SMF Av 504 to at least the stem suboscine lineage (Mayr and Manegold, 2006). Additionally, the hatchet-shaped phalanx II-1 is similar to suboscines and differs from oscines, Acanthisittidae, and Zygodactylidae. This feature is potentially another apomorphy for suboscines, though its distribution has not yet been fully documented.
**Minimum Age Constraint:** 27.0 Ma
**Maximum Age Constraint:** 56.0 Ma
**Age Justification:** The exact horizon from which this specimen was collected was not specified, but the Luberon fossil deposits as a whole are considered to fall within the MP21-MP25 age range. We conservatively use the estimated age of MP25 as a minimum date (see Figure 28.12 of (Gradstein et al. 2020). The oldest reported stem Passeriformes are from the early Eocene. Furthermore, no crown Passeriformes of any type are found in Eocene deposits such as the Green River Formation, Messel Formation, London Clay Formation, or Fur Formation, each of which otherwise preserves an abundance of small bird fossils. These deposits are all from the Northern Hemisphere. Eupasseres appear to have originated in the Southern Hemisphere, which has a much poorer fossil record for small birds. Nevertheless, several Oligocene-Miocene fossils of basal members of Eupasseres have been described from European deposits, which indicate that the clade was not entirely restricted to the Southern Hemisphere early in their evolution and the Early Eocene serves as a conservative maximum.

**Calibrated Node:** Crown Menuridae
**Fossil Specimen:** *Menura tyawanoides* QM F.20887 (AR 11466), carpometacarpus
**Phylogenetic Justification:** *Meruna tyawanoides* was referred to the genus *Menura* by the following features: ligamental attachment of the pisiform process prominent and single, external ligamental attachment prominent, carpometacarpus is proportionally stout, prominence present on metacarpal II about midway between proximal and distal ends on the external border, external carpal trochlea less pronounced, intermetacarpal tuberosity with wide base, and ligamental groove on the external surface of the distal half of carpometacarpus is deep and well defined (Boles, 1995).
**Minimum Age Constraint:** 16.0 Ma
**Maximum Age Constraint:** 34.0 Ma
**Age Justification:** The fossil is from the Upper Site (System B), Riversleigh World Heritage Area, Queensland, Australia. The principal fossil-bearing sites from this locality can be divided into three 'systems' with systems B and C yielding most fossil passerines. The site producing *M. tyawanoides* derives from the Upper Site of System B, which is interpreted as belonging to Faunal Zone B (early Miocene). Faunal Zone B is inferred to span 16Ma-20Ma based on biocorrelation (reviewed by Woodhead et al. 2016). Arena et al. (2016) assigned the Upper Site specifically to B3, the youngest subdivision, based on refinement of regional biostratigraphy. Given this evidence, we use 16Ma as the minimum age for this divergence. We base the maximum on the upper range of the possible age of the oldest confirmed crown Eupasseres, which are from the Oligocene (MP21-MP25) of Luberon, France (Mayr and Manegold, 2006).

**Calibration:** Stem Meliphagidae
**Fossil Specimen:** Meliphagidae indet. QM F20622

**Phylogenetic Justification:** The referral of partial tarsometatarsus total-clade Meliphagidae is based on a suite of character states, including an enlarged canalis for m. flexor hallucis longus, a triangular-shaped trochlea metatarsi II, a large and deep fossa metatarsi I extending at least to the midline of the tarsometatarsus, and the medial side of trochlea metatarsi III projecting more than the lateral side (Boles, 2005). Although some other Australasian taxa including Pomatostomidae exhibit generally similar tarsometatarsi, these taxa can be easily distinguished from Meliphagidae.
**Minimum Age:** 12.9 Ma
**Maximum Age:** 34.0 Ma
**Age Justification:** The Ringtail site forms part of Riversleigh System C ( Travouillon et al, 2006), and based on species assemblage biocorrelation is thought to be of Middle Miocene age (12-15). The Model 1 isochron equilibrium age obtained for this site by Woodhead et al. (2016) corresponds to 13.56Ma ± 0.66Ma; thus, we recommend a minimum age of 14.22Ma for this specimen. We base the maximum on the upper range of the possible age of the oldest confirmed crown Eupasseres, which are from the Oligocene (MP21-MP25) of Luberon, France (Mayr and Manegold, 2006).

**Calibration:** *Orthonyx – Pomatostomus* split
**Fossil Specimen:** *Orthonyx kaldowineryi* QM 56329
**Phylogenetic Justification:** Following Nguyen et al. (2014), this fossil tarsometatarsus can be referred to *Orthonyx* on the basis of the two apomorphiesL impressio ligamentum collateralis lateralis low but very large and plantar surface of shaft immediately proximal to the incisura intertrochlearis lateralis deeply excavated, whereas the plantar surface immediately proximal to the incisura intertrochlearis medialis is very shallowly excavated.
**Minimum Age:** 17.72 Ma
**Maximum Age:** 34.0 Ma
**Age Justification:** Neville's Garden Site represents Faunal Zone B at Riversleigh. This site represents an area of cave deposits comprising both karst formations and a pool deposit (Woodhead et al 2016). Two U/Pb radiometric dates (17.85 ± 0.13 Ma vs 18.24 ± 0.29 Ma) from Neville's Garden Site are associated with speleothems that include flowstones wrapping around fossils ( Woodhead et al 2016). These stalagmites are interpreted as contemporary in age with the fossils and the youngest possible date, inclusive of error, is used for a minimum age calibration: 17.72. A specimen (QM F30244) from the stratigraphically older Hiatus Site is also referred to *O. kaldowineryi*, but is less complete and thus less confidently referred. Attempts to directly date the Hiatus Site specifically have been unsuccessful thus far, but this specimen is potentially ~24Ma. We base the maximum on the upper range of the possible age of the oldest confirmed crown Eupasseres, which are from the Oligocene (MP21-MP25) of Luberon, France (Mayr and Manegold, 2006).

**Calibration:** Stem Neosittidae
**Fossil Specimen:** *Daphoenositta trevorworthyi* QM F57897
**Phylogenetic Justification:** This fossil tibiotarsus was referred to *Daphoenositta* of the monogeneric Neosittidae (sittellas) by (Nguyen, 2014) based on a diagnostic combination of character states including distal margin of pons supratendineus located proximally of condylus

medialis by a distance about equal to a third of condyle length, lateral and medial bony ridges for attachment of retinaculum m. fibularis long and low, bony ridges for retinaculum m. fibularis distally level with tuberositas retinaculi extensoris lateralis and tuberositates retinaculorum extensorium elongate.
**Minimum Age:** 11.6 Ma
**Maximum Age:** 34.0 Ma
**Age Justification:** Rick's Sausage Site is not precisely dated, but is regarded as middle Miocene in age (Archer et al., 1997; Travouillon et al., 2006, 2011), and assigned to Faunal Zone C on the basis of stage-of-evolution biocorrelation. Our recommended minimum age accommodates uncertainty in the age of the upper part of this zone, and is thus placed at 11.6Ma (Woodhead et al. 2016). We base the maximum on the upper range of the possible age of the oldest confirmed crown Eupasseres, which are from the Oligocene (MP21-MP25) of Luberon, France (Mayr and Manegold, 2006).

**Calibration:** Stem Cracticinae
**Fossil Specimen:** *Kurrartapu johnnguyeni* QM F56251
**Phylogenetic Justification:** QM F56251 comprises a tarsometatarsus. Nguyen et al (2013) identified a unique apomorphy uniting *Kurrartapu* with *Cracticus* + *Strepera* (note, these authors included *Melloria quoyi* within the genus *Cracticus*). The incomplete ossification of the retinaculum extensorium tarsometatarsi is not known in any other passerine group except for this clade. Thus, despite the limited material, the fossil taxon *Kurrartapu* can be placed at least as the sister taxon of the clade uniting *Cracticus, Strepera,* and *Melloria* to the exclusion of other Artamidae.
**Minimum Age:** 14.5 Ma
**Maximum Age:** 34.0 Ma
**Age Justification:** The Price is Right Site site from which QM F56251was collected has been interpreted as belonging to Faunal Zone B (early Miocene). However, a review of regional biostratigraphy by Woodhead et al. (2016) found this site to be one of the more poorly constrained Riversleigh localities, with a possible range of encompassing the upper part of Faunal Zone B (B2 or B3) or even the lower part of Faunal Zone C (C1). We thus use the estimated upper age boundary of lower Faunal Zone C (14.5Ma: figure 6 of Woodhead et al. 2016) as a minimum age. An indeterminate scapula from the Miocene St Bathans Fauna (Worthy et al. 2007) potentially represents an older record of Cracticinae, but is only slightly older than QM F56251. We base the maximum on the upper range of the possible age of the oldest confirmed crown Eupasseres, which are from the Oligocene (MP21-MP25) of Luberon, France (Mayr and Manegold, 2006).

**Calibration:** Stem *Ammodramus*
**Fossil Specimen:** *Ammodramus hatcheri* (=*Paleostruthus hatcheri*) USNM 6647
**Phylogenetic Justification:** Steadman (1981) considered USNM 6647 to be more closely related to *Ammodramus savannarum* than other sparrows based on a combination of six characters: (1) crista tomialis slightly concave in ventral aspect, (2) os premaxillare tapering abruptly, with anterior tip very sharp, (3) os premaxillare shallow in lateral aspect, with no distinct bend in the medial os nasale, (4) nares large, rounded anteriorly, (5) posterior border of ventral surface of

os premaxillare only slightly anterior to junction of lateral os nasale and os maxillare, and 6) medial groove on ventral surface of os premaxillare distinct, but not deep. We concur that the morphology precisely matches that of extant *Ammodramus* and differs from other sparrows.
**Minimum Age:** 7.5 Ma
**Maximum Age:** 18.6 Ma
**Age Justification:** The fossil was collected from Quarry E of the Long Island local fauna sites, Phillips County, Kansas, USA. An age of Hh1 (early Hemphilian) is established for Long Island Quarry (Janis et al. 2008). This mammal age spans 7.5-9.0 Ma (Gradstein et al 2012: figure 29.9). The lower end of this age range is used for the minimum age. Although the fossil record of sparrows is poor, the regional record of passerines provides the context for formulating a maximum age. The fossil record of crown Passeriformes in North America is shallow. Aside from *Ammodramus hatcher*i, only a handful of Miocene songbird fossils have been reported including *Palaeoscinis turdirostris* (Howard, 1957), *Miocitta galbreathi* (Brodkorb, 1972), and several still undescribed fossils from the Truckee Formation (Ksepka et al., 2013). The oldest reported North American songbirds are putative records of Parulidae from the Early Miocene (Hemingfordian) Thomas Farm locality, which remain undescribed (Becker, 1987). This distribution suggests that New World sparrows had not originated prior to the Miocene. We use the upper age boundary of the Hemingfordian (18.6Ma; Gradstein et al., 2012: figure 29.9), which conservatively encompasses all North American crown passerine records, as a maximum.

##### References for fossil calibrations


Aguilar, J.-P. et al. (106 authors). 1997. Synthèses et tableaux de corrélations / Syntheses and corrélation tables. In: Aguilar, J.-P., Legendre, S. & Michaux, J. (Editors), Actes du Congres BiochroM'97. Mémoires et Travaux de l'Ecole Pratique des Hautes Etudes, Institut de Montpellier, 21: 769-805.

Archer M, Hand S, Godthelp H, Creaser P 1997. Correlation of the Cainozoic sediments of the Riversleigh World Heritage fossil property, Queensland, Australia. Mémoires et travaux de l'Institut de Montpellier(21): 131-152.

Arena DA, Travouillon KJ, Beck RM, Black KH, Gillespie AK, Myers TJ, Archer M, Hand SJ 2016. Mammalian lineages and the biostratigraphy and biochronology of Cenozoic faunas from the Riversleigh World Heritage Area, Australia. Lethaia 49(1): 43-60.

Ballmann P 2004. Fossil Calidridinae (Aves: Charadriiformes) from the Middle Miocene of the Nordlinger Ries. Bonner Zoologische Beitrage: 101-114.

Benito J, Kuo PC, Widrig KE, Jagt JW, Field DJ. 2022. Cretaceous ornithurine supports a neognathous crown bird ancestor. Nature 612(7938): 100-105.

Benton MJ, Donoghue PC, Asher RJ, Friedman M, Near TJ, Vinther J 2015. Constraints on the timescale of animal evolutionary history.

Bertelli S, Chiappe LM, Mayr G 2011. A new Messel rail from the Early Eocene Fur Formation of Denmark (Aves, Messelornithidae). . Journal of Systematic Palaeontology 9: 551-562.

Bertelli S, Lindow B, Dyke GJ, Mayr G 2013. Another charadriiform-like bird from the lower Eocene of Denmark. Paleontological Journal 47: 1282-1301.

Boellstorff J 1978. Chronology of some late Cenozoic deposits from the central United States and the Ice Ages.

Boessenecker SJ, Boessenecker RW, Geisler JH 2018. Youngest record of the extinct walrus Ontocetus emmonsi from the Early Pleistocene of South Carolina and a review of North Atlantic walrus biochronology. Acta Palaeontologica Polonica 63(2).

Boles WE 1995. The world's earliest songbird (Aves: Passeriformes). Nature 374: 21-22.

Bourdon E, Bouya B, Iarochene M 2005. Earliest African neornithine bird: a new species of Prophaethontidae (Aves) from the Paleocene of Morocco. Journal of Vertebrate Paleontology 25: 157-170.

Bourdon E, Kristoffersen AV, Bonde N 2016. A roller-like bird (Coracii) from the Early Eocene of Denmark. Scientific Reports 6(1): 34050.

Bourdon E, Mourer-Chauviré C, Amaghzaz M, Bouya B 2008. New specimens of *Lithoptila abdounensis* (Aves, Prophaethontidae) from the Lower

Paleogene of Morocco. Journal of Vertebrate Paleontology 28(3): 751-761.

Brand L, Urbina M, Chadwick A, DeVries TJ, R. E 2011. A high resolution stratigraphic framework for the remarkable fossil cetacean assemblage of the Miocene/Pliocene Pisco Formation, Peru. Journal of South American Earth Sciences 31: 414-425.

Brigham-Grette J, Carter L 1992. Pliocene marine transgressions of northern Alaska: circumarctic correlations and paleoclimatic interpretations. Arctic: 74-89.

Brodkorb P 1972. Neogene fossil jays from the Great Plains. Condor 74: 347-349.

Browning JV, Miller KG, McLaughlin Jr PP, Edwards LE, Kulpecz AA, Powars DS, Wade BS, Feigenson MD, Wright JD 2009. Integrated sequence stratigraphy of the postimpact sediments from the Eyreville core holes, Chesapeake Bay impact structure inner basin. The ICDP-USGS deep drilling project in the Chesapeake Bay impact structure: Results from the Eyreville core holes: Geological Society of America Special Paper 458: 775-810.

Butler RJ, Brusatte SL, Reich M, Nesbitt SJ, Schoch RR, Hornung JJ 2011. The sail-backed reptile Ctenosauriscus from the latest Early Triassic of Germany and the timing and biogeography of the early archosaur radiation. PLoS One 6(10): e25693.

Chambers LM, Pringle M, Fitton G, Larsen LM, Pedersen AK, Parrish R 2003. Recalibration of the Palaeocene-Eocene boundary (P-E) using high precision U-Pb and Ar-Ar isotopic dating. Geophysical Research Abstracts, EGS-AGU-EUG Joint Assembly, Nice, 6th-11th April 2003: 9681–9682.

Chandler RM, Parmley D 2003. The earliest North American record of auk (Aves: Alcidae) from the Late Eocene of central Georgia. Oriole 68: 7-9.

Chen A, White ND, Benson RB, Braun MJ, Field DJ 2019. Total-evidence framework reveals complex morphological evolution in nightbirds (Strisores). Diversity 11(9): 143.

Clarke JA, Tambussi CP, Noriega JI, Erickson GM, Ketcham RA. 2005. Definitive fossil evidence for the extant avian radiation in the Cretaceous. Nature 433(7023): 305-308.

Cooper RA 2004. The New Zealand Geological Timescale. Lower Hutt, Institute of Geological and Nuclear Sciences.

Crane A, Benito J, Chen A, Musser G, Torres CR, Clarke JA, Lautenschlager S, Ksepka DT, Field DJ. 2024. Taphonomic damage obfuscates interpretation of the retroarticular region of the Asteriornis mandible. Geobios.

De Pietri VL, Costeur L, Güntert M, Mayr G 2011. A revision of the Lari (Aves, Charadriiformes) from the early Miocene of Saint-Gérand-le-Puy (Allier, France). Journal of Vertebrate Paleontology 31(4): 812-828.

Deng T, Qiu Z-X, Wang B-Y, Wang X, Hou S-K 2013. Late Cenozoic biostratigraphy of the Linxia basin, northwestern China. Fossil mammals of Asia: 243-273.

El Adli JJ, Wilson Mantilla JA, Antar MSM, Gingerich PD 2021. The earliest recorded fossil pelican, recovered from the late Eocene of Wadi Al-Hitan, Egypt. Journal of Vertebrate Paleontology 41(1): e1903910.

Feduccia A, Voorhies MR 1992. Crowned cranes (Gruidae: Balearica) in the Miocene of Nebraska. Nat. Hist. Mus. Los Angel. Cty. Sci. Ser 36: 239-248.

Field DJ, Hsiang AY. 2018. A North American stem turaco, and the complex biogeographic history of modern birds. BMC Evolutionary Biology, 18: 1-16.
Field DJ, Benito J, Chen A, Jagt JW, Ksepka DT. 2020. Late Cretaceous neornithine from Europe illuminates the origins of crown birds. Nature 579(7799): 397-401.
Field DJ, Benito J, Werning S, Chen A, Kuo PC, Crane A, Widrig KE, Ksepka DT, Jagt JW. 2024. Remarkable insights into modern bird origins from the Maastrichtian type area (north-east Belgium, south-east Netherlands). Netherlands Journal of Geosciences 103: p.e15.
Göhlich UB 2007. The oldest fossil record of the extant penguin genus *Spheniscus* - a new species from the Miocene of Peru. Acta Paleontologica Polonica 52: 285-298.
Göhlich UB, Mayr G 2018. The alleged early Miocene Auk Petralca austriaca is a Loon (Aves, Gaviiformes): restudy of a controversial fossil bird. Historical Biology 30(8): 1076-1083.
Gradstein FM, Ogg JG, Schmitz MD, Ogg GM 2020. Geologic time scale 2020, Elsevier.
Höck V, Daxner-Höck G, Schmid HP, Badamgarav D, Frank W, Furtmüller G, Montag O, Barsbold R, Khand Y, Sodov J 1999. Oligocene-Miocene sediments, fossils and basalts from the Valley of Lakes (Central Mongolia)—an integrated study. . Mitteilungen der Geologischen Gesellschaft 90: 83-125.
Hood SC, Torres CR, Norell MA, Clarke JA 2019. New fossil birds from the earliest Eocene of Mongolia. American Museum Novitates 2019(3934): 1-24.
Hornibrook NdB, Brazier RC, Strong CP 1989. Manual of New Zealand Permian to Pleistocene foraminiferal biostratigraphy. Paleontological Bulletin(56).
Ksepka DT, Balanoff AM, Bell MA, Houseman MD 2013. Fossil grebes from the Truckee Formation (Miocene) of Nevada and a new phylogenetic analysis of Podicipediformes (Aves). Palaeontology 56: 1149-1169.
Ksepka DT, Clarke JA, Nesbitt SJ, Kulpe F, Grande L 2013. Fossil evidence of wing shape in a stem relative of swifts and hummingbirds (Aves, Pan-Apodiformes). Proceedings of the Royal Society B 280: 20130580.
Ksepka DT, Early CM, Dzikiewicz K, Balanoff AM 2023. Osteology and neuroanatomy of a phasianid (Aves: Galliformes) from the Miocene of Nebraska. Journal of Paleontology 97(1): 223-242.
Ksepka DT, Fordyce RE, Ando T, Jones CM 2012. New fossil penguins (Aves: Sphenisciformes) from the Oligocene of New Zealand reveal the skeletal plan of stem penguins. Journal of Vertebrate Paleontology 32: 235-254.
Ksepka DT, Stidham TA, Williamson TE 2017. Early Paleocene landbird supports rapid phylogenetic and morphological diversification of crown birds after the K-Pg mass extinction. Proc Natl Acad Sci U S A 114(30): 8047-8052.
Kurochkin E, Dyke G 2011. The first fossil owls (Aves: Strigiformes) from the Paleogene of Asia and a review of the fossil record of Strigiformes. Paleontological Journal 45: 445-458.
Li Z, Zhou Z, Deng T, Li Q, Clarke JA 2014. A falconid from the Late Miocene of northwestern China yields further evidence of transition in Late Neogene steppe communities. Auk 131(3): 335-350.
Louchart A, Tourment N, Carrier J 2011. The earliest known pelican reveals 30 million years of evolutionary stasis in beak morphology. Journal of Ornithology 152(1): 15-20.
Mayr G 2000. Charadriiform birds from the early Oligocene of Céreste (France) and the middle Eocene of Messel (Hessen, Germany). Géobios 33: 625-636.
Mayr G 2001. A cormorant from the late Oligocene of Enspel, Germany (Aves, Pelecaniformes, Phalacrocoracidae). Senckenbergiana Lethaea 81(329-333).
Mayr G, Smith R. 2002. Avian remains from the lowermost Oligocene of Hoogbutsel (Belgium). Bull Inst Roy Sci Nat Belg 72:139–150
Mayr G 2004. Phylogenetic relationships of the early Tertiary Messel rails (Aves, Messelornithidae). Senckenbergiana Lethaea 84: 317-322.

Mayr G 2005. A tiny barbet-like bird from the lower Oligocene of Germany: the smallest species and earliest substantial fossil record of the pici (woodpeckers and allies). The Auk 122(4): 1055-1063.
Mayr G 2005. A new cypselomorph bird from the middle Eocene of Germany and the early diversification of avian aerial insectivores. Condor 107(2): 342-352.
Mayr G 2006. First fossil skull of a Palaeogene representative of the Pici (woodpeckers and allies) and its evolutionary implications. Ibis 148(4): 824-827.
Mayr G 2008. Phylogenetic affinities of the enigmatic avian taxon *Zygodactylus* based on new material from the early Oligocene of France. Journal of Systematic Palaeontology 6: 333-334.
Mayr G. 2008. Phylogenetic affinities and morphology of the late Eocene anseriform bird *Romainvillia stehlin*i Lebedinsky, 1927. Neues Jahrb Geol Palaontol Abh 248:365–380
Mayr G 2009. A well-preserved skull of the "falconiform" bird *Masillaraptor* from the middle Eocene of Messel (Germany). Palaeodiversity 2: 315-320.
Mayr G 2010. Mousebirds (Coliiformes), parrots (Psittaciformes), and other small birds from the late Oligocene/early Miocene of the Mainz Basin, Germany. Neues Jahrbuch für Geologie und Paläontologie-Abhandlungen: 129-144.
Mayr G 2013. Parvigruidae (Aves, core Gruiformes) from the early Oligocene of Belgium. Palaeobiodiversity and Palaeoenvironments 93: 77-89.
Mayr G 2016. The world's smallest owl, the earliest unambiguous charadriiform bird, and other avian remains from the early Eocene Nanjemoy Formation of Virginia (USA). PalZ 90: 747-763.
Mayr G 2022. Paleogene Fossil Birds (Second Edition). Heidelberg, Springer. 239 p.
Mayr G 2022. A partial skeleton of Septencoracias from the early Eocene London Clay reveals derived features of bee-eaters (Meropidae) in a putative stem group roller (Aves, Coracii). Palaeobiodiversity and Palaeoenvironments 102(2): 449-463.
Mayr G, Bertelli S 2011. A record of *Rhynchaeites* (Aves, Threskiornithidae) from the early Eocene Fur Formation of Denmark, and the affinities of the alleged parrot *Mopsitta*. Palaeobiodiversity and Palaeoenvironments 91: 229-236.
Mayr G, Knopf CW 2007. A tody (Alcediniformes: Todidae) from the early Oligocene of Germany. The Auk 124(4): 1294-1304.
Mayr G, Manegold A 2006. A small suboscine-like passeriform bird from the early Oligocene of France. The Condor 108(3): 717-720.
Mayr G, Poschmann M, Wuttke M 2006. A nearly complete skeleton of the fossil galliform bird Palaeortyx from the late Oligocene of Germany. Acta Ornithologica 41(2): 129-135.
Mayr G, Scofield RP 2016. New avian remains from the Paleocene of New Zealand: the first early Cenozoic Phaethontiformes (tropicbirds) from the Southern Hemisphere. Journal of Vertebrate Paleontology 36: e1031343.
Micklich N, Hildebrandt L 2005. The Frauenweiler clay pit (''Grube Unterfeld''). . Kaupia: Darmstädter Beiträge zur Naturkunde 14: 113–118.
Montes M, Nozal F, Santillana SN, Marenssi SA, Olivero E. 2012a. Mapa Geológico de la Isla Marambio (Seymour); escala 1:20.000. Serie Cartográfica Geocientífica Antártica IGME-IAA. Madrid: Instituto Geológico y Minero de España; Buenos Aires-Instituto Antártico Argentino (Mapa Geomorfológico de la Serie, Isla Marambio.
Montes M, Santillana S, Nozal F, Beamud E, Marenssi S. 2012b. Estratigrafía del Maastrichtiense terminal-Paleoceno inferior de la Península Antártica 169: 230–233.
Mourer-Chauviré C. 1993. Les gangas (Aves, Columbiformes, Pteroclidae) du Paleogène et du Miocène inférieur de France. Palaeovertebrata 22: 73-98.
Mourer-Chauviré C. 1996. Paleogene avian localities of France. In: Mlíkovský J. (ed.): Tertiary avian localities of Europe. Acta Universitatis Carolinae, Geologica 39: 567-598.

Mourer-Chauviré C 2008. Birds (Aves) from the Early Miocene of the Northern Sperrgebiet, Namibia. Mem. Geol. Surv. Namibia 20: 147-167.
Murphey PC, Evanoff E. 2007. Stratigraphy, fossil distribution, and depositional environments of the upper Bridger Formation (middle Eocene), southwestern Wyoming. Wyoming State Geological Survey Report of Investigations No. 57.
Musser G, Clarke JA 2020. An exceptionally preserved specimen from the Green River Formation elucidates complex phenotypic evolution in Gruiformes and Charadriiformes. Frontiers in Ecology and Evolution 8: 559929.
Musser G, Ksepka DT, Field DJ 2019. New material of Paleocene-Eocene Pellornis (Aves: Gruiformes) clarifies the pattern and timing of the extant gruiform radiation. Diversity 11(7): 102.
Naish TR, Wehland F, Wilson GS, Browne GH, Cook RA, Morgans HE, Rosenberg M, King PR, Smale D, Nelson CS 2005. An integrated sequence stratigraphic, palaeoenvironmental, and chronostratigraphic analysis of the Tangahoe Formation, southern Taranaki coast, with implications for mid-Pliocene (c. 3.4–3.0 Ma) glacio-eustatic sea-level changes. Journal of the Royal Society of New Zealand 35(1-2): 151-196.
Nesbitt SJ, Ksepka DT, Clarke JA 2011. Podargiform affinities of the enigmatic *Fluvioviridavis platyrhamphus* and the early diversification of Strisores (''Caprimulgiformes'' + Apodiformes). PLoS One e26350.
Nesbitt SJ, Liu J, Li C 2010. A sail-backed suchian from the Heshanggou Formation (Early Triassic: Olenekian) of China. Earth and Environmental Science Transactions of the Royal Society of Edinburgh 101(3-4): 271-284.
Nguyen J, Worthy T, Boles W, Hand S, Archer M 2013. A new cracticid (Passeriformes: Cracticidae) from the Early Miocene of Australia. Emu 113(4): 374-382.
Nguyen JM, Boles WE, Worthy TH, Hand SJ, Archer M 2014. New specimens of the logrunner Orthonyx kaldowinyeri (Passeriformes: Orthonychidae) from the Oligo-Miocene of Australia. Alcheringa: An Australasian Journal of Palaeontology 38(2): 245-255.
Ogg JG, Ogg G, Gradstein FM 2008. The Concise Geologic Time Scale. Cambridge, Cambridge University Press. 177 p.

Olson SL, Feduccia A. 1980. Relationships and evolution of flamingos (Aves: Phoenicopteridae). Smithson Contrib Zool 316:1–73.
Olson S, Rasmussen P 2001. Miocene and Pliocene birds from the Lee Creek Mine, North Carolina. Smithsonian Contributions to Paleobiology 90: 233-365.
Olson SL 1977. A Lower Eocene frigatebird from the Green River Formation of Wyoming (Pelecaniformes, Fregatidae). Smithsonian Contributions to Paleontology 35: 1-33.
Olson SL 1985. The fossil record of birds. In: Farner DS, King JR, Parkes KC ed. Avian Biology. New York, Academic Press. Pp. 79-238.
Olson SL 1987. An early Eocene oilbird from the Green River Formation of Wyoming (Caprimulgiformes: Steatornithidae). Documents des Laboratoires de Geologie de Lyon 99: 57-69.
Olson SL. 1992. A new family of primitive landbirds from the lower Eocene Green River formation of Wyoming. In: Campbell KE, editor. Papers in avian paleontology honoring Pierce Brodkorb. Natural History Museum of Los Angeles County science series 36: 137–60.
Rasmussen DT, Olson SL, Simons EL 1987. Fossil birds from the Oligocene Jebel Qatrani Formation, Fayum Province, Egypt. Smithsonian Contributions to Paleobiology 62: 1-20.
Rich PV, Haarhoff PJ 1985. Early Pliocene Coliidae (Aves, Coliiformes) from Langebaanweg, South Africa. Ostrich 56: 20-41.

Roberts DL, Matthews T, Herries AIR, Boulter C, Scott L, Musekiwa C, Mthembi P, Browning C, Smith RMH, Haarhoff P and others 2011. Regional and Global Context of the Late Cenozoic Langebaanweg (LBW) Palaeontological Site: West Coast of South Africa. Earth Science Reviews.

Seiffert ER 2006. Revised age estimates for the later Paleogene mammal faunas of Egypt and Oman. Proceedings of the National Academy of Sciences 103(13): 5000-5005.

Slack KE, Jones CM, Ando T, Harrison GL, Fordyce RE, Arnason U, Penny D 2006. Early penguin fossils, plus mitochondrial genomes, calibrate avian evolution. Molecular Biology and Evolution 23(6): 1144-1155.

Smith ME, Chamberlain KR, Singer BS, Carroll AR 2010. Eocene clocks agree: Coeval 40Ar/39Ar, U-Pb, and astronomical ages from the Green River Formation. Geology 38: 527–530.

Smith NA 2011. Systematics and evolution of extinct and extant Pan-Alcidae (Aves, Charadriiformes): combined phylogenetic analyses, divergence estimation, and paleoclimatic interactions. Unpublished thesis, The University of Texas at Austin, Austin. 748 p.

Smith NA 2011. Taxonomic revision and phylogenetic analysis of the flightless Mancallinae (Aves, Pan-Alcidae). ZooKeys 91: 1-116.

Smith NA 2015. Sixteen vetted fossil calibrations for divergence dating of Charadriiformes (Aves, Neognathae). Palaeontologia Electronica 18.1.4FC: 1-18.

Smith ND 2010. Phylogenetic analysis of Pelecaniformes (Aves) based on osteological data: implications for waterbird phylogeny and fossil calibration studies. PLoS ONE 5(10): e13354.

Steadman D 1981. A re-examination of Palaeostruthus hatcheri (Shufeldt), a late Miocene sparrow from Kansas. Journal of Vertebrate Paleontology 1(2): 171-173.

Stidham TA 2015. A new species of *Limnofregata* (Pelecaniformes: Fregatidae) from the Early Eocene Wasatch Formation of Wyoming: implications for palaeoecology and palaeobiology. Palaeontology 58: 239-249.

Sun J, Windley BF 2015. Onset of aridification by 34 Ma across the Eocene-Oligocene transition in Central Asia. Geology 43(11): 1015-1018.

Tambussi CP, Degrange FJ, De Mendoza RS, Sferco E, Santillana S. 2019. A stem anseriform from the early Palaeocene of Antarctica provides new key evidence in the early evolution of waterfowl. Zoological Journal of the Linnean Society 186(3): 673-700.

Thomas DB, Tennyson AJD, Scofield RP, Heath TA, Pett W, Ksepka DT 2020. Ancient crested penguin constrains timing of recruitment into seabird hotspot. Proceedings of the Royal Society B: Biological Sciences 287: 20201497.

Travouillon KJ, Archer M, Hand SJ, Godthelp H 2006. Multivariate analyses of Cenozoic mammalian faunas from Riversleigh, northwestern Queensland. Alcheringa: An Australasian Journal of Palaeontology 30(S1): 323-349.

Vellekoop J, Kaskes P, Sinnesael M, Huygh J, Déhais T, Jagt JW, Speijer RP, Claeys P. 2022. A new age model and chemostratigraphic framework for the Maastrichtian type area (southeastern Netherlands, northeastern Belgium). Newsletters on Stratigraphy, 55(4): 479-501.

Woodburne MO, Goin FJ, Raigemborn MS, Heizler M, Gelfo JN, Oliveira EV 2014. Revised timing of the South American early Paleogene land mammal ages. Journal of South American Earth Sciences 54: 109-119.

Woodhead J, Hand SJ, Archer M, Graham I, Sniderman K, Arena DA, Black KH, Godthelp H, Creaser P, Price E 2016. Developing a radiometrically-dated chronologic sequence for Neogene biotic change in Australia, from the Riversleigh World Heritage Area of Queensland. Gondwana Research 29(1): 153-167.

Worthy TH 2011. Descriptions and phylogenetic relationships of a new genus and two new species of Oligo-Miocene cormorants (Aves: Phalacrocoracidae) from Australia. Zoological Journal of the Linnean Society 163(1): 277-314.

Worthy TH, Hand SJ, Archer M 2014. Phylogenetic relationships of the Australian Oligo–Miocene ratite *Emuarius gidju* Casuariidae. Integrative Zoology 2014( 9): 148–166.

Worthy TH, Hand SJ, Nguyen JMT, Tennyson AJD, Worthy JP, Scofield RP, Boles WE, Archer M 2010. Biogeographical and phylogenetic implications of an early Miocene wren (Aves: Passeriformes: Acanthisittidae) from New Zealand. Journal of Vertebrate Paleontology 30: 479-498.

Worthy TH, Tennyson AJD, Scofield RP 2011. An Early Miocene diversity of parrots (Aves, Strigopidae, Nestorinae) from New Zealand. Journal of Vertebrate Paleontology 31(5): 1102-1116.

Yuri T, Kimball RT, Harshman J, Bowie RCK, Braun MJ, Chojnowski JL, Han K-L, Hackett SJ, Huddleston CJ, Moore WS and Reddy, S. 2013. Parsimony and model-based analyses of indels in avian nuclear genes reveal congruent and incongruent phylogenetic signals. Biology 2: 419-444.

#### Other references

Braun, E.L., Oliveros, C.H., White Carreiro, N.D., Zhao, M., Glenn, T.C., Brumfield, R.T., Braun, M.J., Kimball, R.T. and Faircloth, B.C., 2024. Testing the mettle of METAL: A comparison of phylogenomic methods using a challenging but well-resolved phylogeny. bioRxiv, 2024-02. https://doi.org/10.1101/2024.02.28.582627.

Gu, Z., Eils, R., Schlesner, M., 2016. Complex heatmaps reveal patterns and correlations in multidimensional genomic data. Bioinformatics 32, 2847–2849. https://doi.org/10.1093/bioinformatics/btw313.

Harvey, M.G., Bravo, G.A., Claramunt, S., Cuervo, A.M., Derryberry, G.E., Battilana, J., Seeholzer, G.F., McKay, J.S., O'Meara, B.C., Faircloth, B.C., Edwards, S.V., Pérez-Emán, J., Moyle, R.G., Sheldon, F.H., Aleixo, A., Smith, B.T., Chesser, R.T., Silveira, L.F., Cracraft, J., Brumfield, R.T., Derryberry, E.P., 2020. The evolution of a tropical biodiversity hotspot. Science 370, 1343–1348. https://doi.org/10.1126/science.aaz6970.

Huerta-Cepas, J., Serra, F., Bork, P., 2016. ETE 3: reconstruction, analysis, and visualization of phylogenomic data. Mol. Biol. Evol. 33, 1635–1638. https://doi.org/10.1093/molbev/msw046.

Neuwirth, E., Neuwirth, M.E., 2014. Package "RColorBrewer." ColorBrewer palettes.

Oliveros, C.H., Field, D.J., Ksepka, D.T., Barker, F.K., Aleixo, A., Andersen, M.J., Alström, P., Benz, B.W., Braun, E.L., Braun, M.J. and Bravo, G.A., 2019. Earth history and the passerine superradiation. Proceedings of the National Academy of Sciences, 116, 7916-7925. https://doi.org/10.1073/pnas.1813206116.

Paradis, E., Schliep, K., 2019. ape 5.0: an environment for modern phylogenetics and evolutionary analyses in R. Bioinformatics 35, 526–528. https://doi.org/10.1093/bioinformatics/bty633.

Pennell, M.W., Eastman, J.M., Slater, G.J., Brown, J.W., Uyeda, J.C., FitzJohn, R.G., Alfaro, M.E., Harmon, L.J., 2014. geiger v2.0: an expanded suite of methods for fitting macroevolutionary models to phylogenetic trees. Bioinformatics 30, 2216–2218. https://doi.org/10.1093/bioinformatics/btu181.

Yu, G., Smith, D.K., Zhu, H., Guan, Y., Lam, T.T.Y., 2016. ggtree: An R package for visualization and annotation of phylogenetic trees with their covariates and other associated data. Methods Ecol. Evol. 8, 28–36. https://doi.org/10.1111/2041-210X.12628.
